# Drought-Induced Epigenetic Memory in the Cambium of Poplar Trees persists and primes future stress responses

**DOI:** 10.1101/2025.10.14.681991

**Authors:** Alexandre Duplan, Yu-Qi Feng, Gwenvaël Laskar, Bao-Dong Cai, Vincent Segura, Alain Delaunay, Isabelle Le Jan, Christian Daviaud, Anouar Toumi, Francoise Laurans, Mamadou Dia Sow, Odile Rogier, Patrick Poursat, Harold Duruflé, Véronique Jorge, Leopoldo Sanchez, Hervé Cochard, Isabel Allona, Jörg Tost, Régis Fichot, Stéphane Maury

## Abstract

Understanding how perennial plants such as trees perceive, integrate and memorize repeated environmental stresses like water deficit is crucial in the context of climate change.

We investigated short-term and trans-annual memory of water deficit in cambium derived tissues of poplars (*Populus* spp.) using two contrasting genotypes and four genetically modified epitypes with altered DNA methylation machinery.

We found persistent changes in hormone profiles, gene expression and DNA methylation one week after stress relief, consistent with the definition of a multi-layered molecular short-term stress memory. These signatures revealed distinct adaptive strategies between genotypes and marked variability between epitypes, demonstrating that both genetic and epigenetic backgrounds drive stress memory.

Trees exposed to water deficit in Year 1 displayed distinct physiological and molecular responses upon re-exposure in Year 2. The more sensitive genotype showed greater molecular plasticity, whereas the more tolerant genotype exhibited higher stability. A limited set of candidate genes was reactivated upon re-exposure together with persistent drought-induced CG methylation changes, supporting a role in long-term stress imprinting and potential priming.

Our findings highlight the vascular cambium as a key persistent reservoir for short- to trans-annual stress memory in trees. They suggest that mitotically stable CG DNA methylation dynamics, shaped by genetic predisposition and acting through cis- and trans-regulatory routes, help fine-tune growth–survival strategies over longer time frames. This contrasts with the predominantly short-term stress memory described in annual species. These insights open perspectives for harnessing epigenetic variation in tree breeding and management under increasing drought frequency.

## Introduction

Trees, as sessile and perennial organisms, must continually adjust to environmental fluctuations to grow, reproduce, and survive. Acclimation, defined as phenotypic adjustment to changing conditions (i.e. plasticity; Nicotra et al., 2010), is critical for sustaining fitness across diverse environments. While genetic variation underlies much phenotypic diversity, recent research highlights the role of epigenetic mechanisms in shaping plant acclimation (Kakoulidou et al., 2021; Westerband et al., 2021). Epigenetics refers to heritable modifications in gene expression that occur without changes in DNA sequence (Richards, 2006). In the context of climate change, such dynamic modifications are particularly relevant for long-lived woody species exposed to recurrent trans-annual stresses, where memory operates within the same individual across successive growing seasons rather than being reset each generation, as in annuals such as Arabidopsis. Epigenetic regulation may therefore buffer environmental variability across years and decades (Gallusci et al., 2023), with implications for breeding and conservation of stress-resilient forest resources (Holeski et al., 2012; Plomion et al., 2018).

Among epigenetic marks, DNA methylation is widely studied for its role in genome stability and transcriptional control (Mhiri et al., 2022; Muyle et al., 2022; Ramakrishnan et al., 2022). In plants, methylation occurs in CpG, CHG, and CHH contexts (Zhang et al., 2018). Its establishment relies on the RNA-directed DNA methylation (RdDM) pathway, in which small RNAs guide DRM2 to specific loci for de novo methylation (Xie et al., 2025). Maintenance involves context-specific machinery: MET1 preserves CpG methylation, CMT3 and CMT2 maintain CHG and CHH methylation, and DDM1 facilitates access of methyltransferases to heterochromatin, whereas active demethylation is mediated by ROS1 and DML enzymes. After DNA replication, parental methylation patterns are restored with high fidelity, particularly in symmetric CG sites, while non-CG methylation depends on histone H3K9me2 recognition and RdDM activity, resulting in greater dynamism. Promoter methylation generally represses transcription (Niederhuth et al., 2016; Zhang et al., 2006), whereas gene body CpG methylation stabilizes housekeeping gene expression and limits aberrant transcription and TE activity (Muyle et al., 2022). Thus, methylome dynamics integrate stability and flexibility, coordinating development and stress responses across timescales (Lloyd & Lister, 2022; Ramakrishnan et al., 2022). The mitotic stability of CG methylation makes it a strong candidate substrate for long-term somatic memory in perennial meristems, while feedback loops between CHG/CHH methylation, H3K9me2, and small RNAs link DNA methylation to broader chromatin regulation. Consequently, genome-wide methylome mapping captures heritable chromatin states and has become central in plant epigenetics.

Water availability strongly constrains plant productivity and increasing drought frequency drives forest dieback worldwide (Hammond et al., 2022). Severe drought causes mortality through hydraulic failure and carbon imbalance (McDowell et al., 2022), whereas sublethal drought may leave persistent traces shaping future resistance and recovery (ecological memory; Gessler et al., 2020). Stress memory describes how prior exposure leaves molecular, physiological, or epigenetic imprints that modify subsequent responses (Crisp et al., 2016; Hilker et al., 2016). Such memory operates at multiple scales, from somatic persistence within tissues over months or years to transgenerational transmission (Brenya et al., 2022; Farkas & Dobránszki, 2024; Gallusci et al., 2023), and involves coordinated regulation across genome, epigenome, proteome, and metabolome (Galviz et al., 2020; Harris et al., 2023).

Most mechanistic studies have focused on *Arabidopsis thaliana* and short-term priming responses (Harris et al., 2023). Priming refers to pre-conditioning by mild stress, chemical cues, or symbioses that enhances later responses (Lämke & Bäurle, 2017; Mauch-Mani et al., 2017) and often involves histone modifications and non-coding RNAs. In annuals, memory typically resolves within days or, in limited cases, across one generation, reflecting life-cycle constraints. While Arabidopsis has provided major insights (Kakoulidou et al., 2021), the extent to which these mechanisms apply to long-lived trees exposed to repeated stresses over decades remains unclear (Amaral et al., 2020; Gallusci et al., 2023). In perennials, memory must span short-term responses, trans-annual persistence within the same individual, and potentially transgenerational transmission, without complete organismal reset.

In poplar, drought-induced epigenetic memory has been demonstrated in the shoot apical meristem (SAM), involving DNA methylation, hormonal shifts, and transcriptomic changes (Lafon-Placette et al., 2018; Sow et al., 2021; Trontin et al., 2021). Such modifications may persist for months (Le Gac et al., 2018) and transmit mitotically to developing leaves (Le Gac et al., 2019). The SAM is now considered a hotspot of methylation plasticity (Yang & Johannes, 2025). However, whether comparable mechanisms operate in other meristems, such as the vascular cambium responsible for radial growth, and how genetic background influences their stability, remain largely unresolved.

Meristematic tissues are central to plasticity and memory. Persistent changes in transcription factors, metabolites, DNA methylation, and histone marks occur during recovery (Crisp et al., 2016), and phytohormones interact closely with chromatin regulation (Lafon-Placette et al., 2018; Liu et al., 2016; Ojolo et al., 2018; Peirats-Llobet et al., 2016; Yamamuro et al., 2016). As a stem-cell–containing tissue with continuous mitotic activity, the vascular cambium integrates developmental and environmental signals through hormonal gradients (Ding et al., 2024; Fukuda, 2004; Mähönen et al., 2006; Nieminen et al., 2015; Planas-Riverola et al., 2019; Smetana et al., 2019) and epigenetic mechanisms including DNA methylation, histone variants, and non-coding RNAs (Groover, 2025; Inácio et al., 2022). Its initials integrate signals, while wood derivatives provide long-term records, making the cambial zone a candidate reservoir of stress memory (Maury et al., 2019).

In this study, we investigated short-term and trans-annual drought-induced epigenetic memory in cambium-derived tissues of poplar (Populus spp.). We combined two *P. nigra* genotypes from distinct European origins with engineered epitypes of *P. tremula* × *P. alba* altered in methylation and demethylation pathways, including *DDM1* and *DMLs* (Conde et al., 2017a, 2017b; Sow et al., 2021; Vigneaud et al., 2023; Zhu et al., 2013; Xu & Law, 2024). This design allowed us to disentangle genetic and epigenetic contributions to drought memory. By integrating ecophysiology, hormone profiling, transcriptomics, and DNA methylation analyses across two timescales (one week vs. one year), we tested whether the vascular cambium functions as a meristematic reservoir of drought memory, with mitotically stable CG methylation providing a potential molecular backbone, and whether genetic background modulates its establishment and persistence. Overall, our work addresses how perennial meristems integrate environmental information, a key question for understanding acclimation and resilience of forest trees under recurrent drought.

## Material and Methods

### Plant material, growth conditions and experimental design

We conducted experiments on two *Populus nigra* L. genotypes (thereafter referred as *Pn* genotypes) and four *Populus tremula* L. × *Populus alba* L. (INRA 717-1B4) DNA methylation epitypes (thereafter referred as *Ptm* × *Pa* epitypes; see Table 1 and Fig. 1). The *Pn* genotypes (Table 1) were chosen based on available plant material and data previously reported from a wide collection of 1160 cloned genotypes, originating from 14 natural populations across 11 river catchments in 4 European countries, installed in common gardens in France and Italy (more details on the origin of these genotypes are available in the <u>GnpIS Information System</u> (Steinbach et al., 2013). The *Ptm* × *Pa* epitypes were chosen based on previous experiments to include representative transgenic constructs affecting distinct methylation pathways (Table 1).

**Fig. 1.**
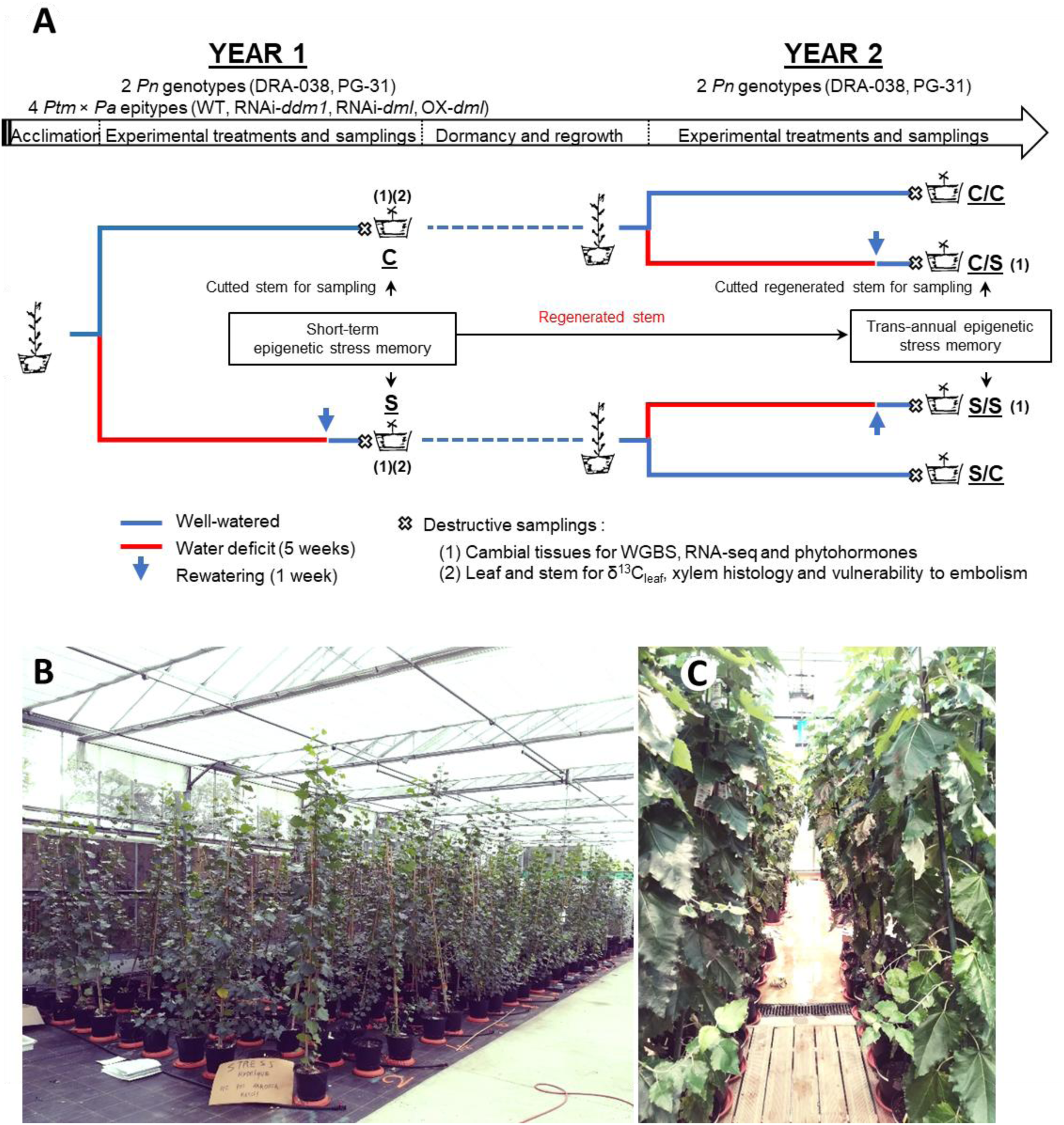
(A) Schematic representation of the experimental design. Experiments were conducted independently on two *P. nigra* (*Pn*) genotypes and four *P. tremula* × *P. alba* (*Ptm* × *Pa*) epitypes. For *Pn* genotypes, experiments were repeated in Year 2 to investigate long-term (trans-annual) memory of water stress. After sampling of cambium from stem in Year 1, *Pn* trees were kept in greenhouse all winter for dormancy, and the regenerated stem in Year 2 was subjected to a new set of treatment and sampling. Abbreviations for treatments: C, control Year 1; S, stressed Year 1; C/C, control Year 1 / control Year 2; C/S, control Year 1 / stressed Year 2; S/S, stressed Year 1 / stressed Year 2; S/C, stressed Year 1 / control Year 2. (B) and (C) Overview of potted trees of *Pn* and *Ptm* × *Pa*, respectively, at the end of the experiment.

**Table 1.**
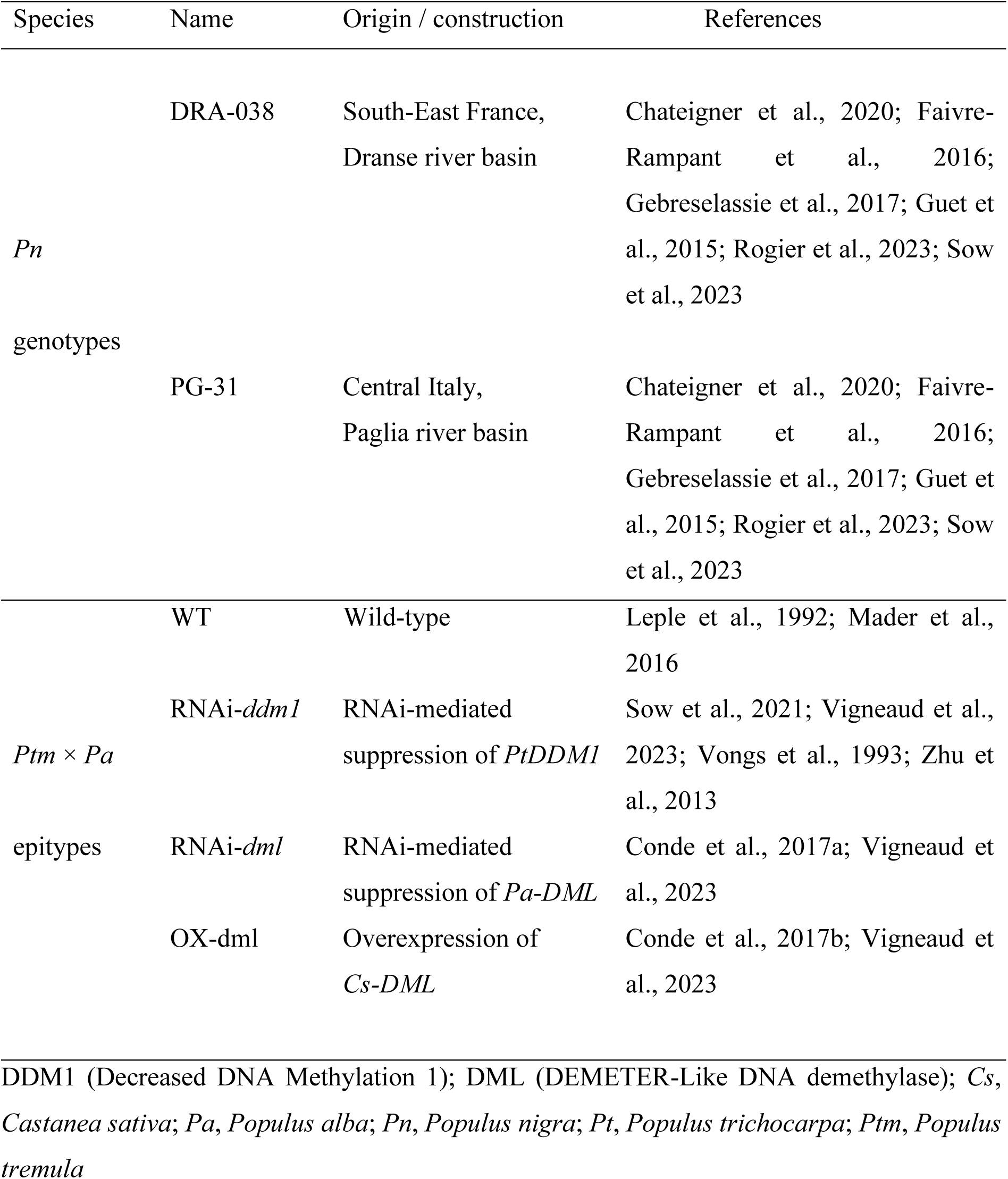
Denomination and characteristics of the plant material used in the study.

| Species | Name | Origin / construction | References |
| --- | --- | --- | --- |
| *Pn*<br>genotypes | DRA-038 | South-East France,<br>Dranse river basin | Chateigner et al., 2020; Faivre-Rampant et al., 2016; Gebreselassie et al., 2017; Guet et al., 2015; Rogier et al., 2023; Sow et al., 2023 |
|  | PG-31 | Central Italy,<br>Paglia river basin | Chateigner et al., 2020; Faivre-Rampant et al., 2016; Gebreselassie et al., 2017; Guet et al., 2015; Rogier et al., 2023; Sow et al., 2023 |
| *Ptm* × *Pa*<br>epitypes | WT | Wild-type | Leple et al., 1992; Mader et al., 2016 |
|  | RNAi-*ddm1* | RNAi-mediated<br>suppression of *PtDDM1* | Sow et al., 2021; Vigneaud et al., 2023; Vongs et al., 1993; Zhu et al., 2013 |
|  | RNAi-*dml* | RNAi-mediated<br>suppression of *Pa-DML* | Conde et al., 2017a; Vigneaud et al., 2023 |
|  | OX-dml | Overexpression of<br>*Cs-DML* | Conde et al., 2017b; Vigneaud et al., 2023 |

DDM1 (Decreased DNA Methylation 1); DML (DEMETER-Like DNA demethylase); *Cs*, *Castanea sativa*; *Pa*, *Populus alba*; *Pn*, *Populus nigra*; *Pt*, *Populus trichocarpa*; *Ptm*, *Populus tremula* The experiment on *Pn* genotypes was initiated from homogeneous dormant cuttings sampled at the end of winter from current-year branches of trees growing at a nearby common garden. Cuttings were rooted in 10 L pots with a potting mix (Klassmann® RHP 25-564) complemented with Osmocote® PG Mix (1 kg/m^3^ of N-P-K 80/35/60) and were grown in a heated greenhouse for two months under controlled conditions [photoperiod 16/8 (day/night, h), air temperature cycles 20/15 (day/night, °C) and daily PPFD of 950 µmol.m^-2^.s^-1^]. Early growing saplings were then transferred to an unheated greenhouse receiving natural daylight and following the course of the natural photoperiod until the end of the experiment in early September. The experiment on *Ptm* × *Pa* epitypes was initiated from *in vitro* micropropagated plantlets. Young acclimated plantlets were first rooted in 2 L pots before being transplanted in 10 L pots using the same soil substrate as for the *Pn* experiment. Plants were then grown in a GMO greenhouse under the same ambient conditions as for *Pn* genotypes.

Experiments on *Pn* and *Ptm* × *Pa* epitypes followed the same experimental design adapted from previous work (Sow et al., 2021). Treatments were initiated on 3-month-old plants and lasted for 6-weeks. Half of the plants were assigned to the well-watered treatment (Control, C) while the other half was assigned to a 5-week water deficit followed by a one-week rewatering (Stressed, S). Plants from the C treatment were maintained close to field capacity during the whole experiment using an automatic drip irrigation system connected to tensiometers (model INT4, Kriwan Industrie-Electronik GmbH, Germany). Plants from the S treatment were maintained between 10 and 20% of soil relative extractable water (REW) based on daily individual measurements of soil water content (Bouyer et al., 2023; Sow et al., 2021) before being rewatered and maintained close to field capacity for one additional week before sampling (see sup. Fig. S1). The total number of plants amounted to 24 for *Pn* genotypes (2 treatments × 2 genotypes × 6 clonal replicates) and 48 for *Ptm* × *Pa* epitypes (2 treatments × 4 epitypes × 6 clonal replicates).

To explore the trans-annual epigenetic memory of the water deficit applied in Year 1, we decided to keep the *Pn* genotypes for one additional year (Year 2). After sampling in Year 1, the aerial stem was cut, allowing the trees to undergo winter dormancy and subsequently regenerate a new stem from the perennial root system in the following growing season. We could not manage such exploratory experiments for the transgenic *Ptm* × *Pa* epitypes because of technical limitations of keeping our transgenic trees in the GMO greenhouse for over a year. In Year 2, these regenerated stems from half of the C plants from Year 1 were assigned to the control well-watered treatment (C/C) or the water deficit treatment followed by rewatering (C/S), while half of the S plants subjected to water deficit in Year 1 (S) were similarly assigned either to the control well-watered treatment (S/C) or the water deficit treatment followed by rewatering (S/S). The treatments applied and the methods used in Year 2 were identical to Year 1 (Fig. 1).

### Plant-level physiological monitoring

Physiological monitoring concentrated on the core experiments conducted in Year 1 on both *Pn* genotypes and *Ptm* × *Pa* epitypes (see below and Fig. 1). Measurements in Year 2 (trans-annual memory in *Pn* genotypes) were restricted to soil water status and growth.

Tree growth was monitored during the experiment by measuring stem height and diameter of all plants every two to three days. Leaf net assimilation rate (A_net_) and stomatal conductance to water vapour (g_s_) were assessed 8-9 times throughout the experiment to evaluate the effects of water deficit on leaf gas exchange physiology (*n* = 3 random replicates per genotype/epitype per treatment at each date). Measurements were performed on fully mature leaves in the top third of the plants using a Li-6400 open path portable photosynthesis system (Li-Cor, Lincoln, NE, USA) equipped with a LED light source (Li-6400-02B) under saturating chamber conditions (photosynthetic photon flux density of 1500 µmol.m^-2^.s^-1^, ambient CO_2_ concentration of 400 ppm and reference vapour pressure deficit maintained close to 1 kPa). Leaf-level intrinsic water-use efficiency (_i_WUE) was computed directly from instantaneous leaf gas exchange as the ratio between A_net_ and g_s_ and assessed retrospectively by measuring bulk leaf carbon isotope composition (δ^13^C_leaf_) on mature leaves formed during the 5-week water deficit period. We also measured daily minimum leaf water potential (Ψ_leaf-min_) repeatedly during the experiment (*n* = 3 random replicates per genotype/epitype per treatment at each date) to assess differences in water relations. Finally, we evaluated xylem resistance to drought-induced embolism in order to evaluate intrinsic resistance to drought-induced hydraulic dysfunction (Fichot et al., 2015). Measurements were performed at the end of the experiment on stems of all well-watered control plants (*n* = 6 per genotype/epitype) using the cavitron centrifugation technique Cochard et al., 2005.

### Sampling of cambium derived tissues

Tissue sampling for molecular analyses was performed in Year 1 (*Pn* genotypes and *Ptm* × *Pa* epitypes, short-term memory) and in Year 2 (*Pn* genotypes, trans-annual memory) at the end of the 1-week rewatering following water deficit (*n* = 3 replicates per genotype/epitype per treatment) (see Fig. 1). We sampled approximately 80 cm-long stem sections starting at 20 cm above the ground (Chateigner et al., 2020). The bark was split using a sterile scalpel and manually detached from the stem before gently scratching tissues on both sides (bark *vs*. stem) thus leading to two distinct tissue fractions. Histological analyses showed that the tissue fraction from the stem side mostly corresponded to young differentiating secondary xylem while the tissue fraction from the bark side contained most of the cambial cells and secondary phloem derivatives (Fig. S2). Sampled tissue fractions were immediately hand-ground in liquid nitrogen using pestles and mortars and stored separately at -80_∘_C pending further milling and individual extraction of genomic DNA and RNA.

### Phytohormone quantification

Phytohormones in cambium-derived tissues (*n* = 3 replicates per genotype/epitype per treatment) were quantified from 20 mg dry powder from pooled tissue fractions using the LC-MS technique (Thermo Scientific Ultimate 3000 UHPLC coupled with a TSQ Quantiva, Waltham, Massachusetts, USA). The following 15 phytohormones were analysed: Auxin (IAA), Abscisic Acid (ABA), 4’-dihydrophaseic acid (DPA; ABA derivative), Jasmonic acid (JA), 12-oxo-phytodienoic acid (OPDA, JA derivative), Salicylic Acid (SA), *cis*-Zeatin-O-glucoside Cytokinin (cZOG), Isopentenyl adenine riboside Cytokinin (iPR), *cis*- or *trans*-Zeatin riboside Cytokinin (cZR or tZR), Gibberellins (GA8, GA12, GA19), castasterone (CS), 6-Deoxocastasterone (6-deoxoCS).

For GA analysis, samples were extracted with 1 mL 80% (v/v) methanol. [^2^H_2_] GA1, [^2^H_2_] GA3, [^2^H_2_] GA4, [^2^H_2_] GA5, [^2^H_2_] GA6, [^2^H_2_] GA7, [^2^H_2_] GA8, [^2^H_2_], [^2^H_2_] GA12, [^2^H_2_] GA15, [^2^H_2_] GA19, [^2^H_2_] GA20, [^2^H_2_] GA24, [^2^H_2_] GA34, [^2^H_2_] GA44, [^2^H_2_] GA51 and [^2^H_2_] GA53 were added to plant samples as internal standards for quantification. The samples were extracted at 4°C for 12 h. The analysis of GA content was performed after sample pre-treatment (Chen et al., 2012). For brassinosteroids (BR) analysis, samples were extracted with 1 mL acetonitrile. Then [^2^H_3_]BL, [^2^H_3_]CS, [^2^H_3_]6-deoxoCS were added as internal standards for quantification. The samples were extracted at 4°C for 12 h. The analysis of BR content was performed after sample pre-treatment (Ding et al., 2013). Finally, for IAA, ABA, JA, SA and cytokinins (CK), samples were extracted with 1 mL acetonitrile. Then [^2^H_2_]IAA, [^2^H_6_]ABA, [^2^H_3_]PA, [^2^H_3_]DPA, [^2^H_3_]ABAGE, [^2^H_4_]7′−OH-ABA, [^2^H_3_]neoPA, [^2^H_4_]SA, [^2^H_6_]iP, [^2^H_6_]iPR, [^2^H_6_]iP9G, [^2^H_5_]tZ, [^2^H_5_]tZR, [^2^H_5_]tZ7G, [^2^H_5_]tZ9G, [^2^H_5_] tZOG, [^15^N_4_]cZ , [^2^H_3_]DHZ, [^2^H_3_]DHZR, [^2^H_3_]DHZ9G, [^2^H_7_]DHZOG were added as internal standard for quantification. The samples were extracted at 4°C for 12 h. The analysis was performed after sample pre-treatment (Cai et al., 2015).

### Sequencing analysis: Transcriptome and Methylome

Poplar transcriptome analysis has previously been described (Sow et al., 2021; Vigneaud et al., 2023). Briefly, total RNA was extracted independently from stem and bark fractions and assessed for integrity before equimolar pooling per individual to represent cambium-derived tissues. RNA-seq was performed on three biological replicates per treatment and genotype/epitype following rRNA depletion and strand-specific library preparation. Libraries were sequenced in paired-end mode on an Illumina NovaSeq platform. Reads were quality-filtered and mapped to the *Populus trichocarpa* v4.1 reference genome. Transcript quantification and differential expression analyses were performed using Salmon and DESeq2. Genes with adjusted p-values ≤ 0.05 and |log2 fold change| ≥ 0.5 were considered differentially expressed. Alternative splicing analyses were conducted using rMATS and SUPPA2. Full pipeline details, parameters, and software versions are provided in Supplementary Methods.

Methylome analysis has previously been described in detail (Sow et al., 2021; Vigneaud et al., 2023; Lesur et al., 2024). An equimolar pool of 2 µg DNA at 100 ng.µL^-1^ was then prepared for each treatment and genotype/epitype from the gDNA samples of the three individual replicates and the 2 tissue fractions (see above). The gDNA pools were then used for whole genome bisulfite sequencing (WGBS) using the Ultralow Methyl-Seq System (Tecan Genomics) (CEA laboratory, CNRGH, Evry, France) (Dugé de Bernonville et al., 2022). Libraries were sequenced in paired-end mode to an average coverage of ∼40×. Methylation analyses were performed using previously established workflows (Sow et al., 2021; Vigneaud et al., 2023; Lesur et al., 2024). Differentially methylated cytosines were identified using methylKit with a minimum coverage of 10×, methylation difference ≥ 5%, and q-value ≤ 0.05. Genes or transposable elements containing at least one differentially methylated cytosine were defined as differentially methylated. Gene Ontology enrichment was conducted using *Arabidopsis thaliana* orthologs with Metascape v3.5 (Zhou et al., 2019), using default parameters. *p-values* were computed with a hypergeometric test, and the Benjamini-Hochberg correction was used for the correction of multiple-testing. Detailed bioinformatic procedures are provided in Supplementary Methods.

### Statistical analysis

Exploration and statistical analyses of physiological data were carried out using the R software (R Core Team https://doi.org/10.59350/t79xt-tf203). All tests were considered significant at *P* < 0.05. Vulnerability curves relating xylem water potential to the percent loss of xylem hydraulic conductance were processed using the R *fitplc* package (Duursma & Choat, 2017) to estimate the water potential generating 50% of embolism (P_50_). Differences between treatments or genotypes/epitypes were tested for all traits using analysis of variance followed by Tukey’s HSD post-hoc test in case of significant differences.

## Results

### Growth and water relations

General patterns for soil REW, growth and physiological traits in Year 1 were similar between *Pn* genotypes and *Ptm* × *Pa* epitypes (Fig. 2, Fig. S1), indicating that although performed independently, experiments were comparable in terms of treatment intensity and general effects on plant physiology. For all genotypes/epitypes, water deficit induced significant growth reductions and changes in leaf physiology (Fig. 2, Fig. S1). Height growth remained steady, although lower than controls, throughout the experiment (Fig. S1). Radial growth seriously inflected after 20 days of water deficit but in most cases did not cease completely (Fig. S1).

**Fig. 2.**
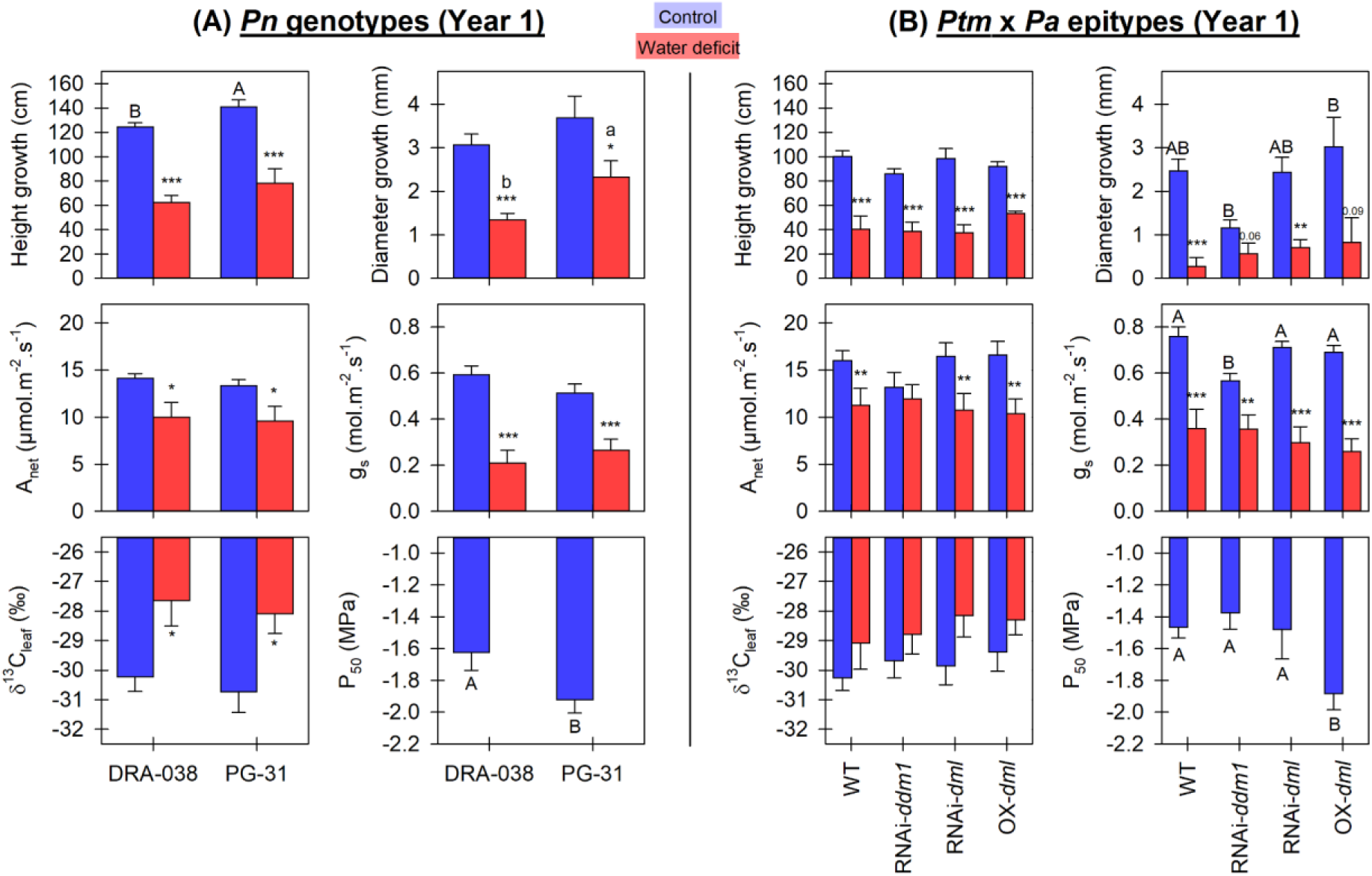
Treatment effects on growth and physiological variables in *Populus nigra* (*Pn*) genotypes (A) and *Populus tremula* × *Populus alba* (*Ptm* × *Pa*) epitypes (B) in Year 1. Abbreviations: A_net_, net CO_2_ assimilation; g_s_, stomatal conductance to water vapour; δ^13^C_leaf_, bulk leaf ^13^C isotopic composition (more negative values indicate lower intrinsic water-use efficiency); P_50_, xylem water potential inducing 50% of embolism (more negative values indicate higher intrinsic tolerance to water deficit). P_50_ was estimated only under control conditions. Bars represent mean values ± SE (*n* = 6 for growth, δ^13^C and P_50_ at the end of the experiment; *n* = 8-9 for A_net_ and g_s_ spread out throughout the 5-weeks period of water deficit). See Materials and Methods for additional information. Different letters (capital letters for controls, lower case letters for water deficit) indicate significant differences between genotypes within treatments following Tukey’s HSD post-hoc test; asterisks indicate significant effects of water deficit (* 0.01 < *P* ≤ 0.05; ** 0.001 < *P* ≤ 0.01; *** *P* ≤ 0.001).

In *Pn* genotypes, PG-31 displayed overall faster growth than DRA-038, especially radial growth under water deficit, and showed significantly higher tolerance to embolism (i.e. lower P_50_) (Fig. 2A). Ψ_min-leaf_ remained high and comparable across genotypes and treatments (−1.25 ± 0.06 MPa on average) suggesting comparable isohydric behaviours. In addition, the two genotypes did not differ significantly in leaf physiological traits and responded similarly to water deficit (Fig. 2A). Based on these observations, PG-31 was considered to be intrinsically more tolerant to water deficit than DRA-038.

*Ptm* × *Pa* epitypes did not differ significantly in height growth regardless of the treatment while differences were more apparent for radial growth (Fig. 2B). Under control conditions, OX-*dml* showed the fastest radial growth while RNAi-*ddm1* was the slowest (Fig. 2B); under water deficit conditions, differences were not significant, probably because of the important growth reductions observed in all lines, but modified lines all tended to show superior radial growth than the WT with OX-*dml* still showing the fastest growth (Fig. 2B). OX-*dml* was also significantly more tolerant to drought-induced embolism (Fig. 2B). Ψ_min-leaf_ remained comparable between epitypes but was slightly reduced in response to water deficit (−0.78 ± 0.04 MPa *vs*. -0.99 ± 0.04 MPa, respectively, *P* < 0.001) suggesting a slightly less isohydric behaviour of *Ptm* × *Pa* epitypes compared to *Pn* genotypes. Leaf traits were not significantly different between epitypes, except for g_s_ of RNAi-*ddm1* under control conditions, and showed similar variations in response to water deficit. Based on these observations, OX-*dml* was the most intrinsically tolerant epitype to water deficit.

We repeated the water deficit experiment in Year 2 on the two *Pn* genotypes to explore stress memory and potential priming. Saplings that experienced water deficit in Year 1 and maintained under control conditions in Year 2 (S/C) exhibited lower growth than saplings from the C/C control treatment in the genotype DRA-038 (Fig. 3), suggesting a legacy effect of water deficit on growth in Year 2 at least in this genotype. In contrast, the growth of saplings that experienced water deficit the two consecutive years (S/S) was not impaired compared to saplings that only experienced water deficit in Year 2 (C/S) in both genotypes (Fig. 3), suggesting that the negative legacy effect of water deficit on growth possibly inherited from Year 1 (see above for DRA-038) was not additive under a repeated stress.

**Fig. 3.**
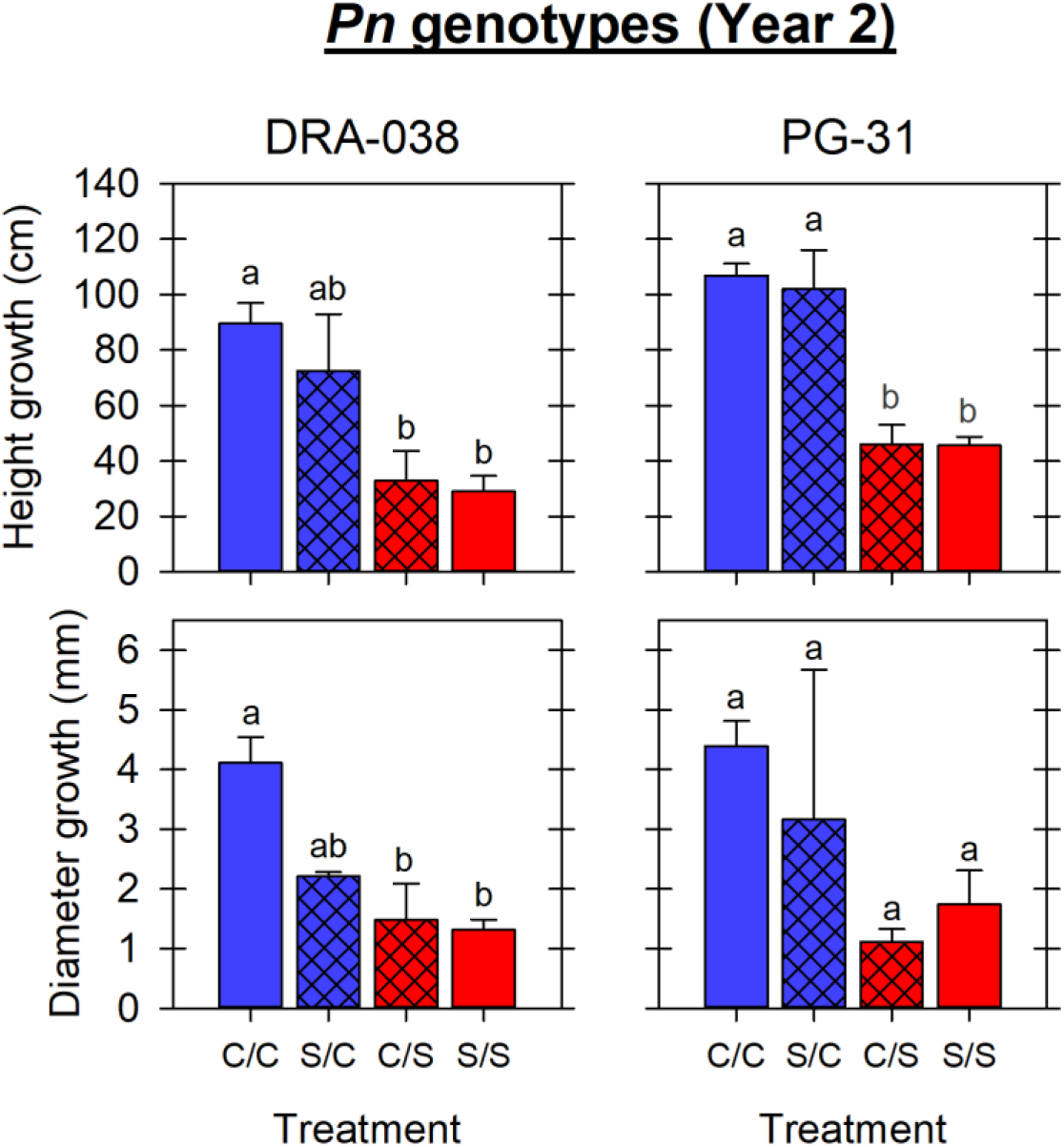
Effects of the repeated water deficit in Year 2 on growth of the two *Populus nigra* (*Pn*) genotypes. Blue bars are for control (C) trees in Year 2 while red bars are for water-stressed (S) trees; hatched bars are for trees that received a different treatment in Year 1 and Year 2. Abbreviations: C/C, control Year 1 / control Year 2; S/C, stressed Year 1 / control Year 2; C/S, control Year 1 / stressed Year 2; S/S, stressed Year 1 / stressed Year 2. Different letters indicate significant differences (*P* < 0.05) between treatments following Tukey’s HSD post-hoc tests.

### Hormonal balance in Cambium-Derived Tissues

Concentrations of 15 phytohormonal compounds were measured in the cambium-derived tissues by LC-MS (Fig. 4). For the *Pn* genotypes, principal component analysis (PCA) revealed that the first three components explained 83.8% of the total variance with principal component (PC) 1 (39.9% of the variance) strongly positively associated with CS, DPA, cZR and GA19 and negatively with JA (Fig. 4A). The PC2 (29.5% of the variance) was mostly positively associated with cZOG and SA and negatively to GA12 (Fig. 4A). PC3 explained 14.4% of the variation, with 6-deoxoCS, tZR, JA, and ABA contributing the most (Fig. S3). Projection of individuals onto the principal plane (Fig. 4B) clearly differentiated genotypes, treatments and years. Genotype DRA-038 was overall more responsive than PG-31, (12/15 hormones significantly different between treatments *vs*. 9/15 in Year 1; 15/15 *vs*. 8/15 in Year 2) (Fig. 4E and Table S1). In Year 1 (S *vs*. C), common responses (i.e. in the same direction) across genotypes were observed for ABA, JA, cZOG, tZR, GA12, and GA19 while opposite responses (i.e. in opposite direction) could be observed for CS (Fig. 4E, Table S1). In Year 2 (S/S *vs*. C/S), common responses across genotypes were observed for JA, OPDA, iPR, and GA19 while opposite responses could be observed for SA, cZOG, tZR and GA12 (Fig. 4E, S Table S1). In addition, the comparison of S *vs*. C patterns (Year 1) with C/S *vs*. S/S patterns (Year 2) revealed a few opposite switches between years for DRA-038 (tZR, cZR and GA19) or PG-31 (cZOG, GA12 and GA19) (Fig. 4E, Table S1).

**Fig. 4.**
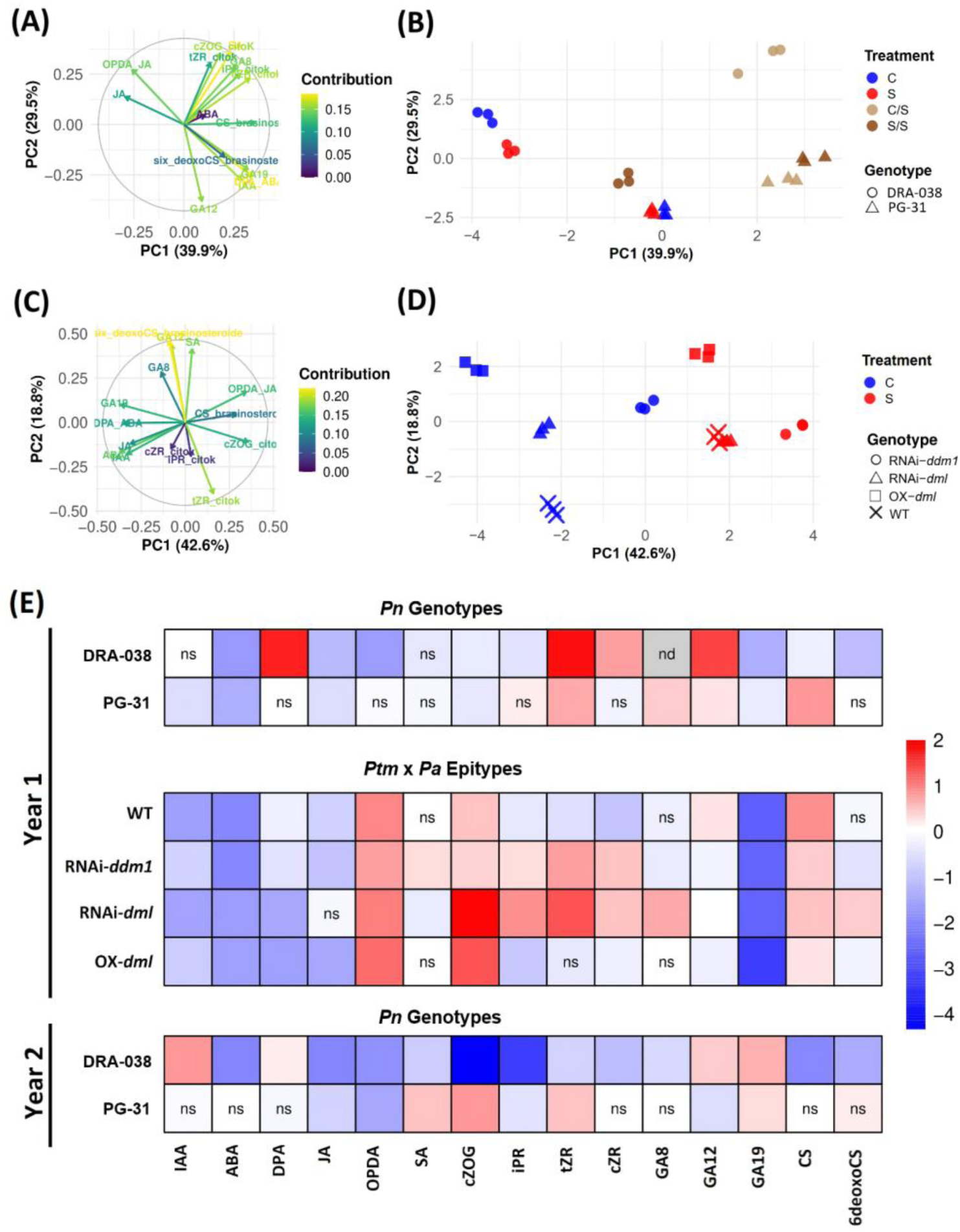
Principal component analysis (A-D) and differential analysis (E-F) of the 15 phytohormonal compounds measured by LC-MS in cambium-derived tissues of *Populus nigra* (*Pn*) genotypes (DRA-038 and PG-31) and *Populus tremula* × *P. alba* (*Ptm* × *Pa*) epitypes (WT, OX-*dml*, RNAi-*dml*, RNAi-*ddm1*). A and C) Correlation circles in the two main components for *Pn* genotypes and *Ptm* × *Pa* epitypes, respectively. B and D) Projections of individuals in the main plane for *Pn* genotypes and *Ptm* × *Pa* epitypes for each treatment, respectively. E) Heatmap of hormone variation in response to water deficit for *Pn* genotypes and *Ptm* × *Pa* epitypes. Colours represent the effect size estimate computed as the log2FoldChange (log2FC) of hormone concentrations (in Year 1, red indicates higher concentrations in the water deficit (S) condition relative to the control (C) condition, while blue indicates lower concentrations; in Year 2, red indicates higher concentrations in the S/S treatment relative to the C/S treatment, while blue indicates lower concentrations, grey indicates no data). Abbreviations: C, control in Year 1; S, stressed in Year 1; C/S, control in Year 1 and stressed in Year 2; S/S, stressed in Years 1 and 2. Auxin (IAA), abscisic acid (ABA), 4’-dihydrophaseic acid (DPA), jasmonic acid (JA), 12-oxo-phytodienoic acid (OPDA), salicylic acid (SA), *cis*-zeatin-O-glucoside cytokinin (cZOG), isopentenyl adenine riboside cytokinin (iPR), *cis*- or *trans*-zeatin riboside cytokinin (cZR or tZR), gibberellin (GA), castasterone (CS), 6-deoxocastasterone (6deoxoCS). **Transcriptome analysis** Differentially expressed transcripts (DETs)

For the *Ptm* × *Pa* epitypes, PCA revealed that the first three components explained 76.6% of the total variance with PC1 (42.6% of the variance) positively associated with GA19, ABA, DPA, IAA, and JA and negatively with OPDA and cZOG Fig. 4C). PC2 (18.8% variance) was associated positively with 6-deoxoCS, GA12, and SA and negatively with tZR (Fig. 4C). PC3 explained 15.2% of the variation, with iPR and cZR contributing most (Fig. S4). Projection of individuals onto the first PCs clearly differentiated epitypes and treatments (Fig. 4D). While epitypes showed distinct projections in control conditions, they tended to group together in stressed conditions, except for OX-*dml*. About half of the compounds (7/15) showed common responses to water deficit among the epitypes (IAA, ABA, DPA, OPDA, cZOG, GA 19 and CS) (Fig. 4E), while the other half showed epitype-specific or non-significant changes. The most notable changes for each epitype (*vs*. WT) were higher contents in gibberellins (GA8 and GA19) and lower content in cytokinin (tZR) for OX-*dml*, variable amounts of cytokinins for RNAi-*dml*, higher amounts of CS and lower amounts of ABA and GA19 for RNAi- *ddm1* (Fig. S5).

Among all genotypes and epitypes, ABA and JA consistently decreased in response to water deficit, indicating a common signature across genetic backgrounds (Fig. 4E). Quantitatively, the main discriminator between genotypes and epitypes was the baseline level of DPA, which was ∼10-fold higher in *Pn* genotypes compared to *Ptm × Pa* epitypes (Table S1).

In *Pn* genotypes, 54% to 70% of the 52,000 annotated poplar transcripts were detected across conditions and years, with the highest proportion observed in DRA-038 under the C/S treatment in Year 2 (Fig. S6, Fig. 5A, 5B). In Year 1, 292 and 122 DETs were observed between C and S treatments in DRA-038 and PG-31, respectively (Fig. 5A), indicating a stronger transcriptional responsiveness or memory in DRA-038 as previously observed for hormones. In Year 2, we found 141 and 136 DETs between C/S and S/S treatments in DRA-038 and PG-31, respectively (Fig. 5A, 7C). Functional annotation revealed that DETs in Year 1 in both genotypes mainly involved metabolic and developmental processes and phosphatidylinositol signalling (Fig. 5B). In Year 2, PG-31 displayed additional tags related to cambium activity and epigenetic regulation (Fig. 5B), suggesting a shift toward regulatory processes upon re-exposure.

**Fig. 5.**
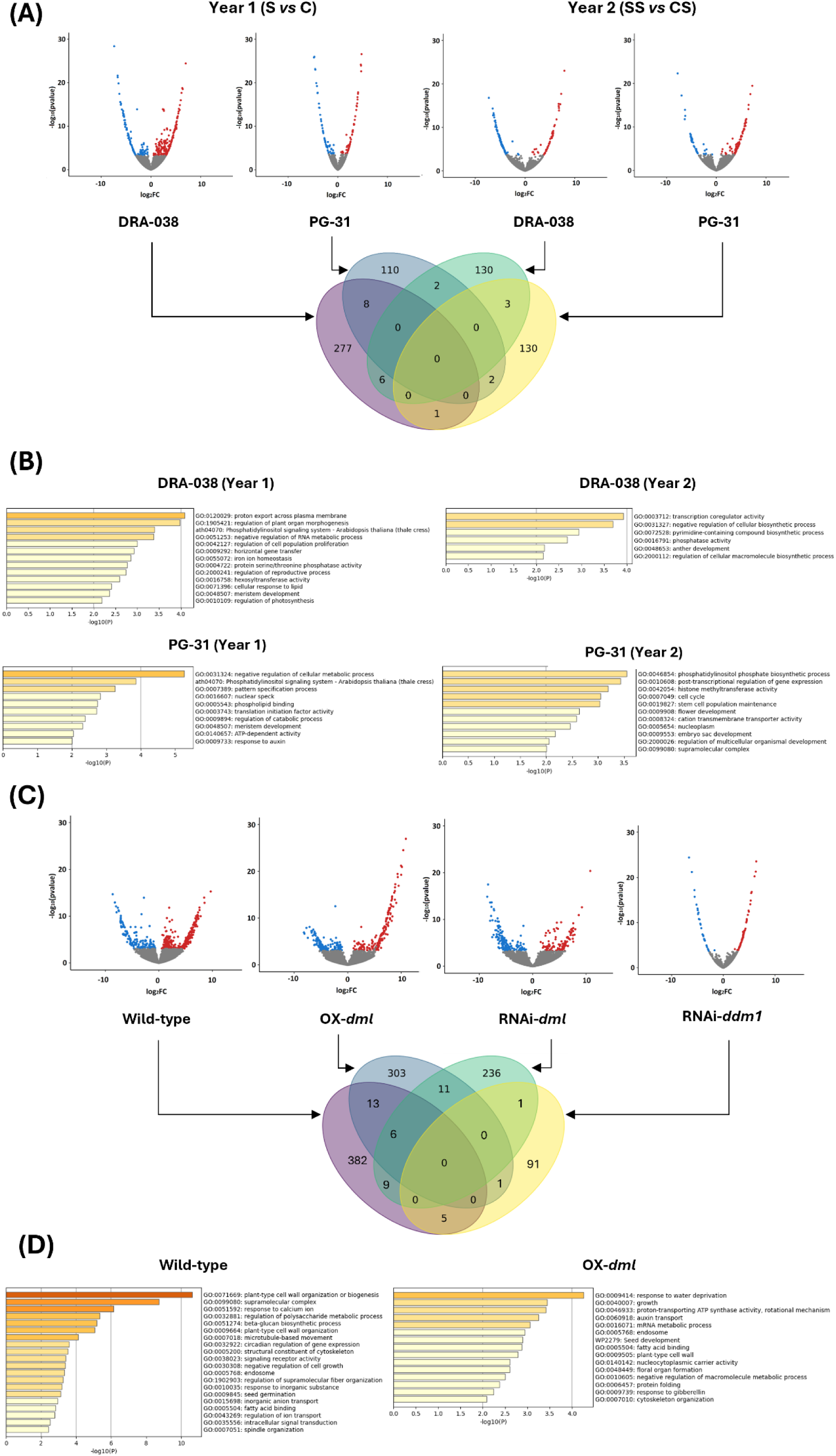
Analysis of differentially expressed transcripts (DETs) in *Pn* genotypes and *Ptm* × *Pa* epitypes under water deficit conditions. (A) Volcano plots showing DETs between stress (S) and control (C) conditions for the two *Pn* genotypes (PG-31 and DRA-038) in Year 1 (S vs. C) and Year 2 (SS vs. CS). Venn diagram indicates the overlap of DETs identified between genotypes and years. (B) Gene Ontology (GO) enrichment analysis of DETs identified in PG-31 and DRA-038 in Year 1 and Year 2. (C) Volcano plots of DETs between stress and control conditions for the *Ptm* × *Pa* epitypes (Wild-type, OX-*dml*, RNAi-*dml* and RNAi-*ddm1*). The Venn diagram shows the overlap of DETs among epitypes. (D) GO enrichment analysis of DETs identified in Wild-type and OX-*dml* epitypes. For all volcano plots, colored dots represent DETs passing the significance threshold (FDR < 0.05) and a log₂ fold change threshold of ±0.5 (blue: downregulated; red: upregulated), while grey dots indicate non-differentially expressed transcripts. For GO enrichment analyses in epitypes, OX-*dml* was selected as the most water-deficit-resistant epitype based on ecophysiological data, Fig. 2). Metascape analyses for RNAi-*dml* and RNAi-*ddm1* are available in Fig. S10).

The number of genotypic DETs (DRA-038 *vs*. PG-31) varied from 297 (Year 2 in S/S condition) to 1930 (Year 1 in S condition) depending on treatments and years (Fig. S7). Importantly, 254 genotypic DETs were conserved between C (Year 1) and C/S (Year 2) treatments, all with a comparable expression pattern (up or down-regulated) except one (AMP-dependent synthetase and ligase *Potri.017G138350*); 168 genotypic DETs were conserved between S (Year 1) and S/S (Year 2) treatments, all with a comparable expression pattern supporting the persistence of genotype-specific transcriptional signatures across years.

The majority of Differentially Spliced Events (DSEs) were genotype-specific (Fig. S8), with a conservation of 21 DSEs in PG-31 between Year 1 and Year 2. Across *Pn* genotypes, 351 stress-induced and 287 genotype-specific DSEs (mostly A3SS, SE, and A5SS types; Fig. S9 A; see materials and methods for DSE classification) were detected, representing a small fraction of the known poplar DSEs (over 9 000 presents in the reference genome *P. trichocarpa* V4.1). Only nine overlapped with DETs, six of which were involved in chromatin remodelling and hormone (Auxin and JA) signalling pathways (Table 2; Table S2) highlighting a limited but functionally relevant convergence between transcriptional and post-transcriptional regulation.

**Table 2.**
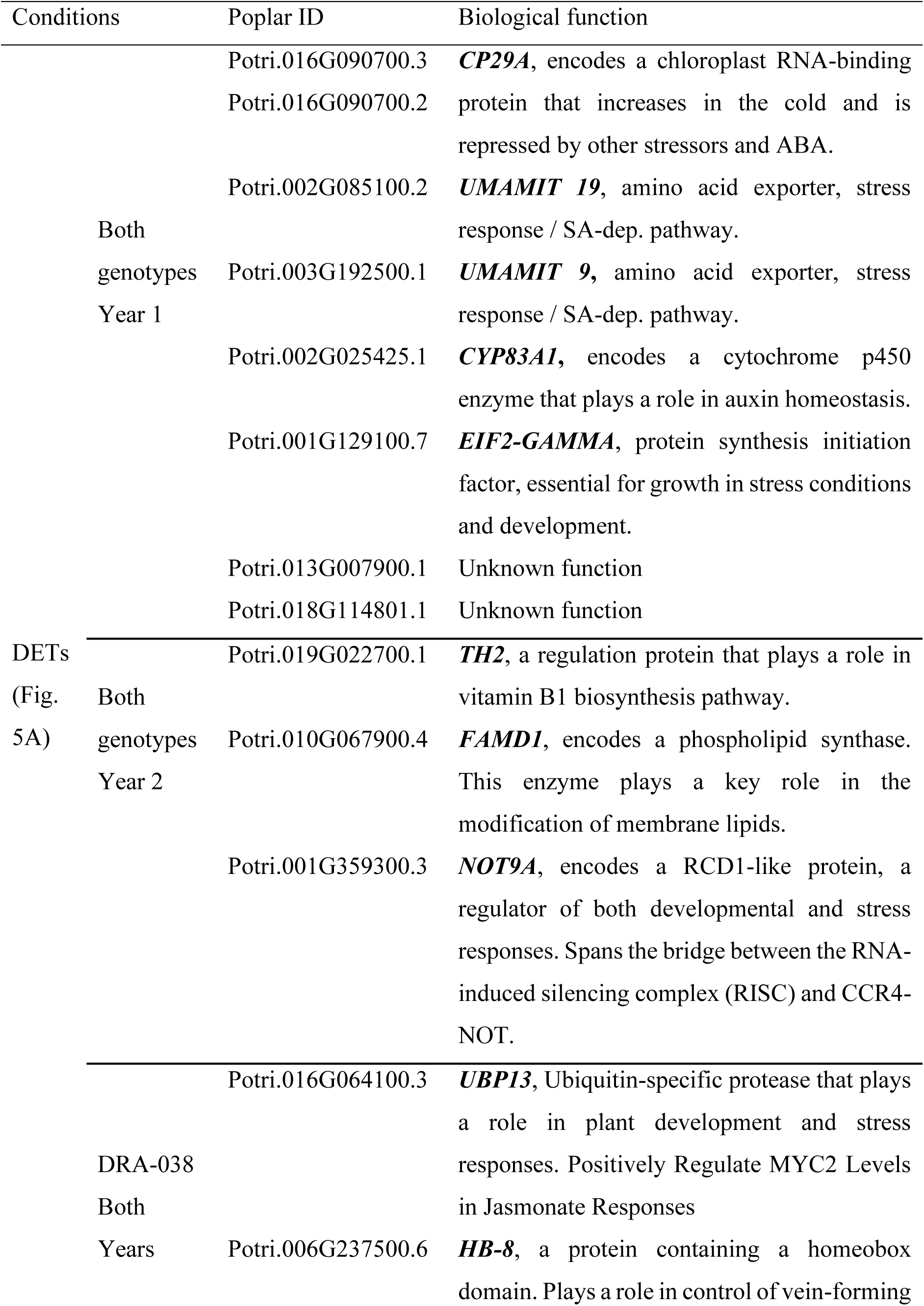
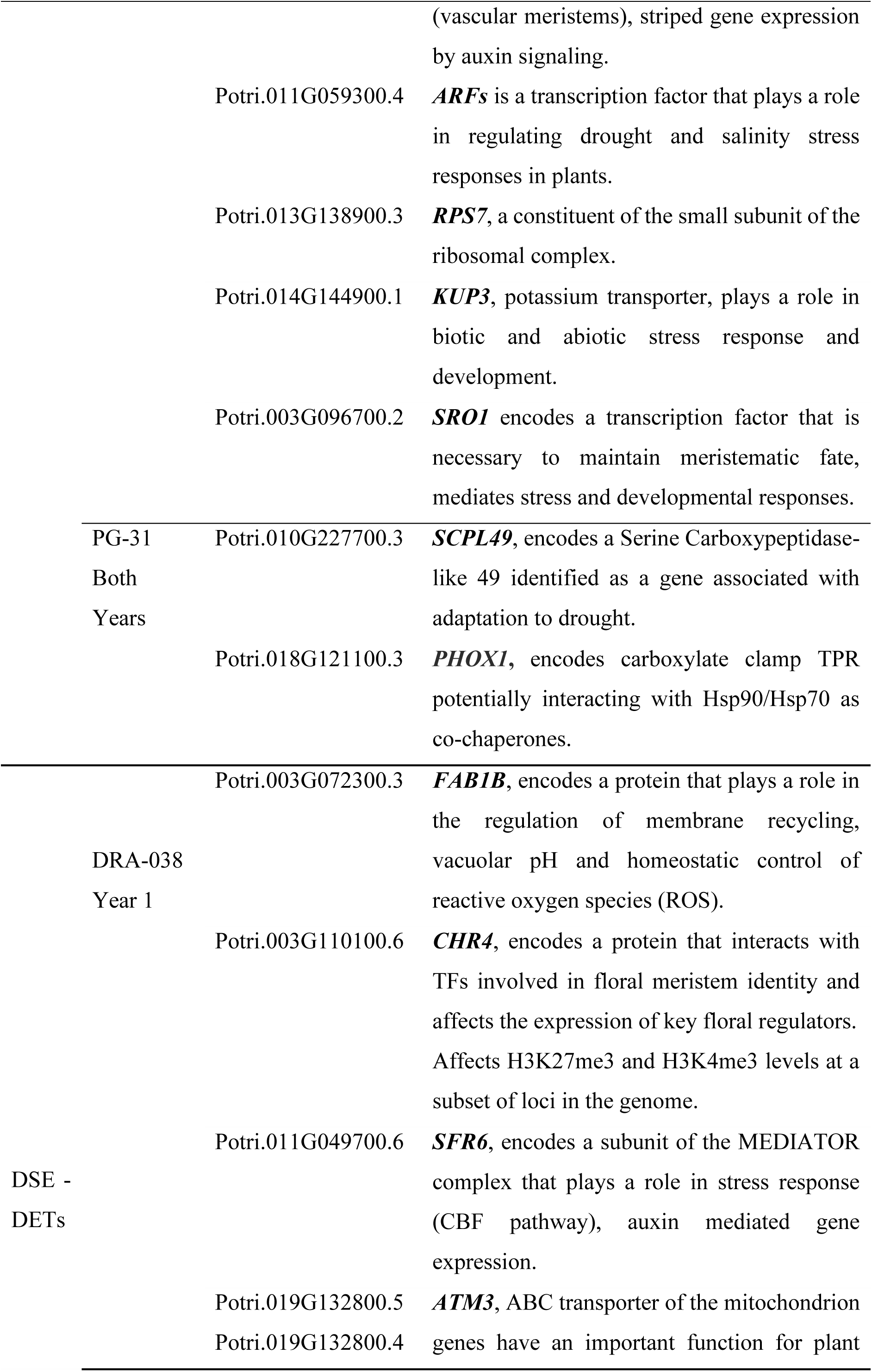
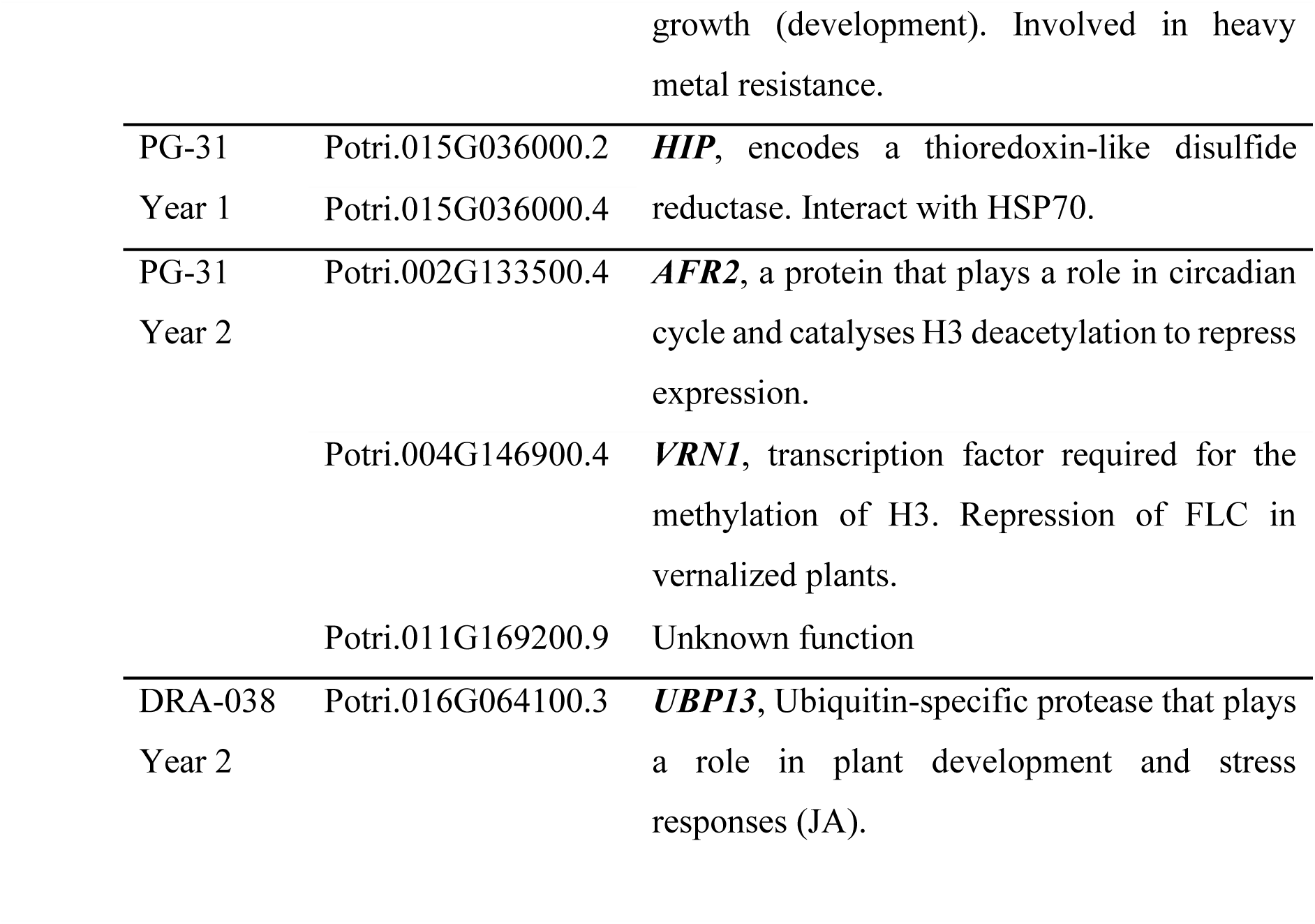
Overview of the candidate genes found in the differential expression (DET) or differential splicing (DSE) analyses carried out by genotype in *Pn* (. **Fig 5A; see also table S2. for more details).**

In *Ptm × Pa* epitypes, transcript detection ranged from 16189 (OX-*dml*, S condition) to 28883 (OX-*dml*, C condition) (Fig. 5C, Fig S6, Fig. S11). Differential expression analysis revealed from 98 (RNAi-*ddm1*) to 415 (WT) DETs in response to water deficit (Fig. 5C), with 46 overlapping with those detected in *Pn* genotypes (Fig. S12), indicating partial conservation of drought-responsive pathways across genetic backgrounds. Most stress-induced DETs were line-specific and related to cell wall modification or growth processes (Fig. 5D, Fig. S10). The OX-*dml* line showed more up-regulated DETs and specific enrichment for “response to water deprivation”, “auxin transport” and “response to gibberellins” (Fig. 5C, 5D), consistent with its ecophysiological resilience profile.

While the majority of DSEs were epitype-specific (Fig. S8) with 257 DSEs detected (Fig. S8, Fig. S9A), of which 12 and 30 were also classified as DETs, we could also detect 181 stress-induced DSEs. GO enrichment analysis shows genes related to “positive regulation of reproductive processes”, including several developmental transcription factors involved in developmental or hormonal signalling (Table S5 and Fig. S9B), suggesting that alternative splicing may contribute to stress-responsive developmental reprogramming.

Overall, transcriptomic analyses indicate that both short-term and trans-annual drought memory are supported by a limited yet persistent subset of DETs and DSEs, largely genotype-or epitype-specific but corresponding to a possible functionally hub of regulatory nodes underlying cambial drought memory.

### Methylome analysis

#### Variation in methylation contexts

In *Pn* genotypes, stress-induced DMCs were primarily found in the CG context, followed by CHG and CHH, with overall a balanced proportion of hyper- and hypomethylation (Fig. 6A). PG-31 showed a markedly higher number of DMCs compared to DRA-038, although DRA-038 showed the strongest relative increase in CG DMCs from Year 1 to Year 2 (22-fold) (Fig. 6A).

**Fig. 6.**
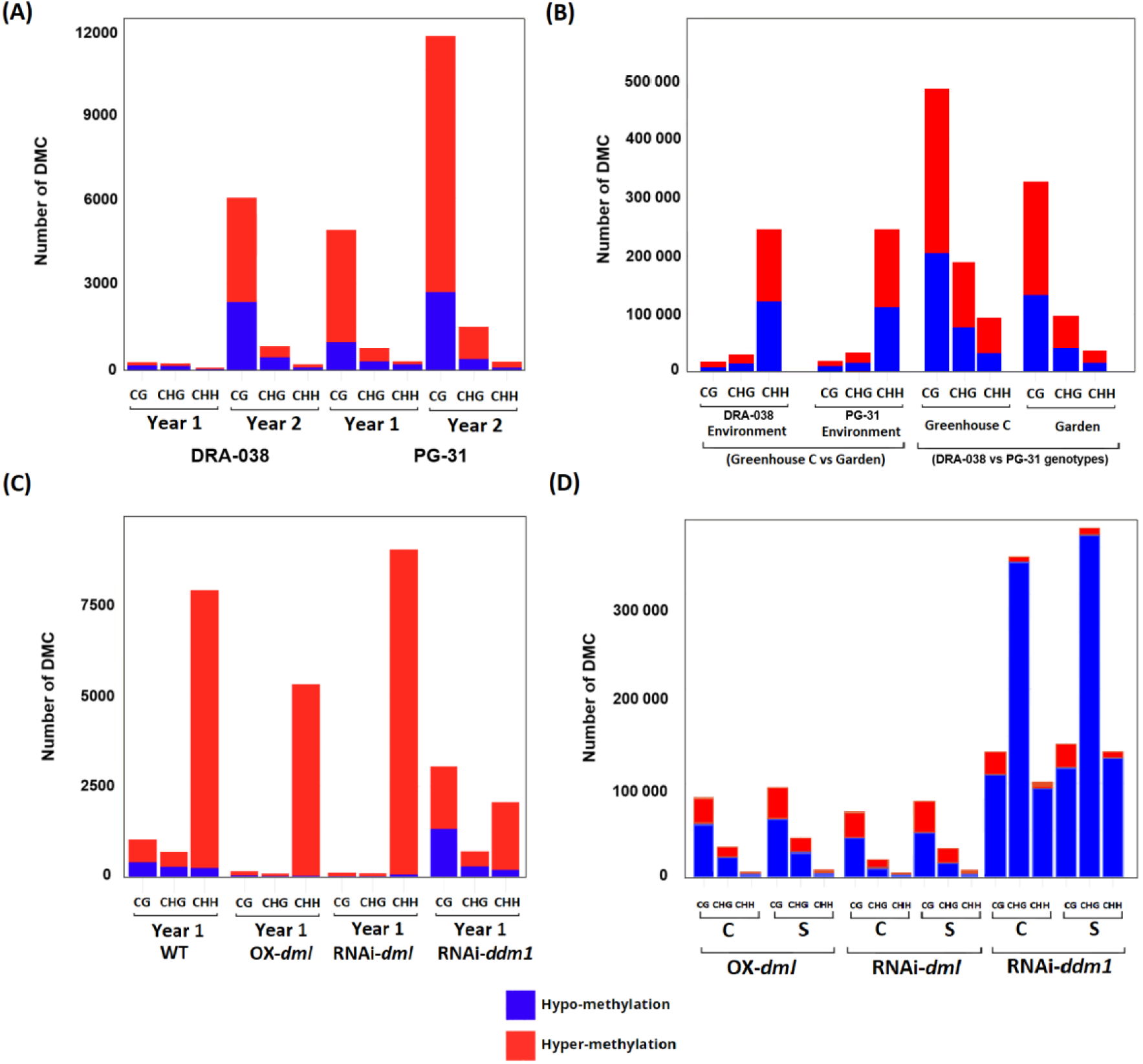
Number of Differentially Methylated Cytosines (DMCs) per methylation context (CG, CHG, CHH). A) DMCs in *Pn* genotypes (DRA-038 and PG-31) in response to treatments in Year 1 (S *vs*. C) and Year 2 (S/S *vs*. C/S). B) DMCs between control greenhouse conditions (this study) and previously published data from an outdoor common garden in the two *Pn* genotypes (DRA-038 and PG-31) and DMCs between genotypes located in the greenhouse (this study) or outdoor common garden (Sow et al. 2023). DMCs in the common garden were obtained from three years old clonal copies of the same genotypes installed in an outdoor common garden (details on the experimental setup are available in Sow et al. 2023). C) DMCs in *Ptm* × *Pa* epitypes (WT, OX-*dml*, RNAi-*dml*, RNAi-*ddm1*) in response to treatment in Year 1 (S *vs*. C). D) DMCs between *Ptm* × *Pa* WT and other epitypes (OX-*dml*, RNAi-*dml* or RNAi-*ddm1*) under control (C) or stressed (S) conditions.

We compared the DMCs identified in the two *Pn* genotypes under controlled greenhouse conditions (present study) with DMCs identified in the same two genotypes but grown in an outdoor common garden (Sow et al., 2023) (Fig. 6B). Environmentally-induced DMCs (greenhouse *vs*. common garden) were predominantly found in the CHH context (up to 248 000), with partial conservation (45-65%) across genotypes (Fig. 6B; Fig. S15). In contrast, genotype-specific DMCs (PG-31 versus DRA-038) were mainly found in CG (up to 490 000) compared to CHG and CHH contexts and showed low conservation (<5% for the 3 contexts) across growing environments, highlighting strong genotype-by-environment interactions (Fig. 6B; Fig. S15).

In *Ptm* × *Pa* epitypes, stress-induced DMCs in WT were distributed across contexts, with 1030 CG and 686 CHG DMCs, values intermediate between DRA-038 (266 CG / 226 CHG) and PG-31 (4889 CG / 757 CHG) in Year 1. In contrast, CHH DMCs were markedly higher (7912 in WT vs. 62 in DRA-038 and 299 in PG-31, corresponding to ∼127× more than DRA-038 and 26× more than PG-31; Fig. 6C). Thus, the strong enrichment in WT was specific to the CHH context, while CG and CHG levels fell within the range observed in *Pn* genotypes. RNAi-*ddm1* differed from other lines by presenting more DMCs in CG (3036 CG, close to PG-31 Year 1 values) and elevated CHH DMCs (2068 CHH), with a relatively more balanced proportion of hyper- and hypomethylation. Line-specific DMCs (i.e. comparison to WT) were 10–30 times more abundant than stress-induced DMCs, especially in RNAi-*ddm1*, and were primarily found in CG except for the RNAi-*ddm1* line showing more DMCs in CHG (Fig. 6D).

#### Methylation Changes in Genes and Transposable Elements

In *P. nigra* genotypes, 15.8% (Year 2) to 33.4% (Year 1) of stress-related DMCs were localized in TEs (differentially methylated TEs, DMTEs) (Fig. 7A). These DMTEs were genotype-specific (Fig. S16), with PG-31 exhibiting stronger stress-induced TE hypermethylation than DRA-038 (Fig. 7B). In contrast, 46.3–61.3% of stress-induced DMCs were located in genes (differentially methylated genes, DMGs), predominantly within exons (Fig. 7A; Fig. S17). DMG numbers ranged from 221 (DRA-038, Year 1) to 4,523 (PG-31, Year 2) (Fig. S18). Functional enrichment highlighted broad metabolic categories, including ATP-dependent and catalytic activities (Fig. S19), consistent with global metabolic adjustment to water deficit. Notably, shared DMGs increased from 19 (Year 1) to 1,119 (Year 2) (Fig. S18), suggesting progressive convergence of methylation signatures upon stress re-exposure.

**Fig. 7.**
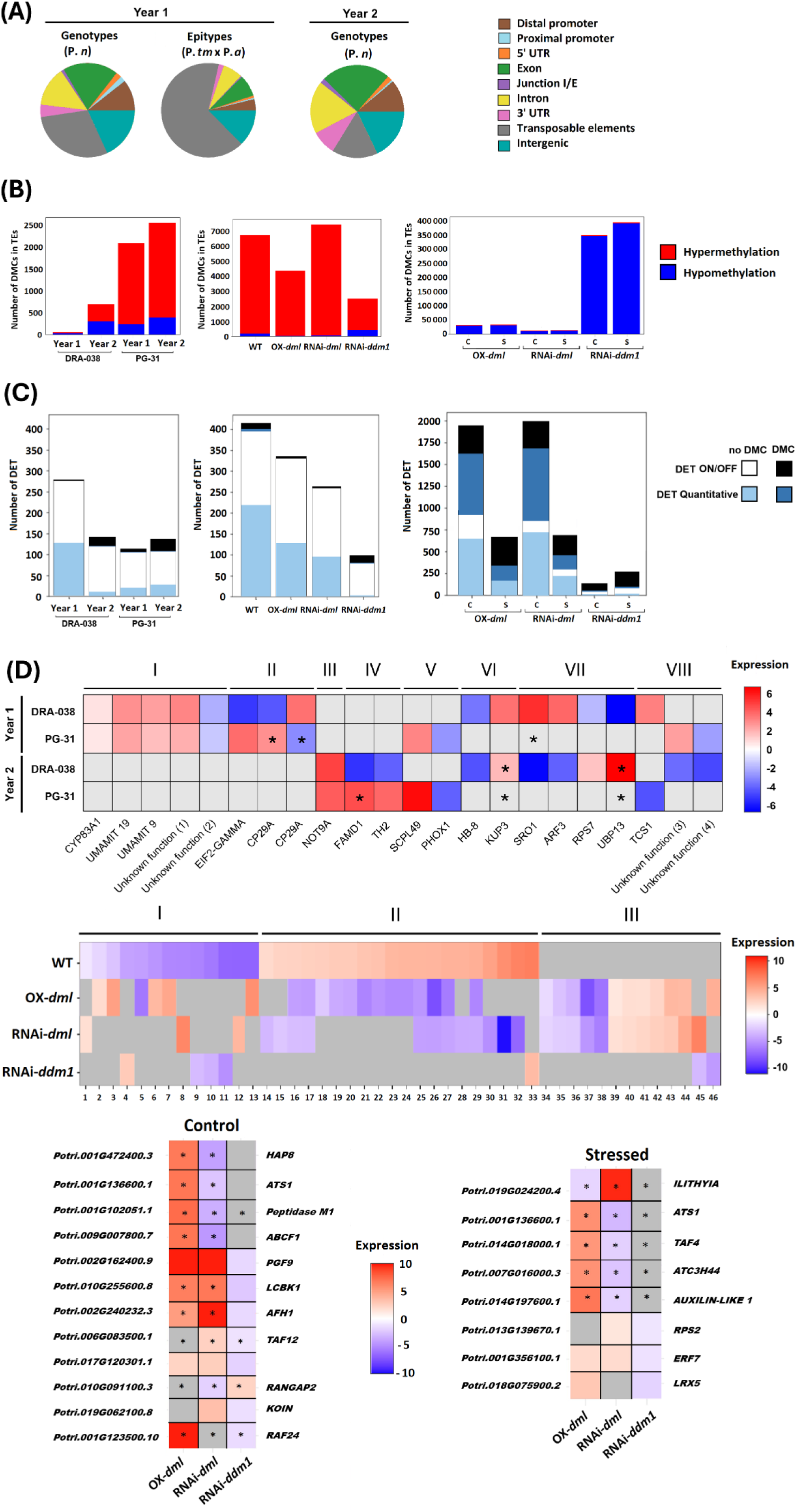
Characterization of stress-induced differentially methylated cytosines (DMCs) and correlation with differentially expressed transcripts (DETs) in *Pn* genotypes and *Ptm* × *Pa* epitypes. (A) Genomic distribution of stress-induced DMCs across genomic features (distal promoter, proximal promoter, 5′UTR, exon, junction I/E, intron, 3′UTR, transposable elements and intergenic regions) identified in *Pn* genotypes (DRA-038 and PG-31) and in *Ptm × Pa* epitypes (WT, OX-*dml*, RNAi-*dml*, RNAi-*ddm1*). For genotypes, DMCs were identified in Year 1 (S vs. C) and Year 2; genes containing DMCs were annotated (Table S6). (B) Number of stress-induced DMCs located in transposable elements (TEs). In *Pn* genotypes, DMCs were quantified in Year 1 (S vs. C) and Year 2 (S/S vs. C/S). In *Ptm × Pa* epitypes, stress-induced DMCs were quantified in Year 1 (S vs. C). In addition, the number of line-specific DMCs (comparison relative to WT) in TEs is shown for each epitype under control (C) and stressed (S) conditions. Hypermethylated and hypomethylated DMCs are indicated in red and blue, respectively. (C) Number of stress-induced DETs in *Pn* genotypes (Year 1 and Year 2) and in *Ptm × Pa* epitypes (Year 1). For epitypes, line-specific DETs (line vs. WT) are shown under control (C) and stressed (S) conditions and classified as either “ON/OFF” (transcripts detected exclusively in one condition and absent in the other) or “Quantitative” (transcripts detected in both conditions with significant differential expression; log₂FC > 0.5, p < 0.05). DETs were further categorized according to the methylation status of their associated gene (presence or absence of DMCs). Gene-level annotations are provided in Table S7. (D) Heatmaps of differentially expressed transcripts. Top panel: Heatmap of common DETs identified in *Pn* genotypes across both years. Blue indicates decreased expression and red increased expression (log₂FC). Asterisks (*) denote genes identified as differentially methylated under the corresponding conditions. Transcripts are grouped into eight clusters (I–VIII): I) similar regulation in both genotypes in Year 1; II) opposite regulation between genotypes in Year 1; III) similar regulation in both genotypes in Year 2; IV) opposite regulation between genotypes in Year 2; V) similar regulation across both years for PG-31; VI) similar regulation across both years for DRA-038; VII) opposite regulation across both years for DRA-038; VIII) genotype-and year-dependent expression patterns. Annotated DETs are listed in Table 2 and Table S2. “Unknown (1)” to “Unknown (4)” correspond to Potri.013G007900.1, Potri.018G114801.1, Potri.009G005300.9 and Potri.010G167100.23, respectively. Middle panel: Heatmap of the 46 stress-induced DETs (S vs. C) identified in at least two of the four epitypes. Gene annotations are provided in Table S3. Expression changes are represented as log₂FC (blue: downregulation; red: upregulation). Bottom panel: Heatmaps of epitype-specific DETs (line vs. WT) displaying opposite expression patterns under control (C) or stressed (S) conditions. Asterisks (*) indicate genes associated with DMCs. Candidate gene annotations are provided in Table S4.

In Year 1, only eight DETs were shared between DRA-038 and PG-31 (Fig. 7C; Table 2), indicating that short-term transcriptional memory relies on a limited core of conserved stress-responsive genes. These included two UMAMIT amino acid transporters (salicylic acid pathway) and CYP83A1 (auxin metabolism), all similarly regulated (cluster I), whereas three DETs—including EIF2-GAMMA and two alternative splicing events of CP29A—displayed opposite regulation between genotypes (cluster II; Fig. 7C; Table S2).

In Year 2, only three DETs were shared (clusters III–IV), including an RCD1-like protein (NOT9a), a regulator of stress and developmental responses via RNA-induced silencing complex activity (Fig. 7C; Table 2). Across years, trans-annual conservation remained limited (two DETs in PG-31; six in DRA-038), involving genes linked to abiotic stress, vascular meristem development, and auxin or jasmonate signaling (Table S2). Importantly, conserved DETs in PG-31 showed amplified responses in Year 2, whereas those in DRA-038 exhibited variable direction or magnitude, supporting genotype-dependent reinforcement versus reprogramming dynamics under repeated drought.

Most DETs across genotypes and years were not associated with DMCs (Fig. 7C). The small subset of overlapping DET–DMC genes was primarily detected in Year 2 and belonged to the ON/OFF category (24 in DRA-038; 30 in PG-31), suggesting discrete cis-regulatory switches upon stress re-exposure. These included Potri.005G182100 (H3K4 histone methyltransferase) and Potri.017G070500 (5′–3′ exonuclease), both exhibiting alternative splicing between oppositely expressed isoforms (Table S7).

In *P. tremula × P. alba* epitypes, 65.9% of stress-induced DMCs localized to TEs—mainly LTR-Gypsy and Copia elements—and were predominantly hypermethylated (Fig. 7A, 7B; Fig. S16; Fig. S20). RNAi-*ddm1* displayed abundant hypomethylated DMTEs, consistent with DDM1’s role in heterochromatic methylation and TE silencing.

Only 16.5% of stress-induced DMCs were gene-associated, enriched for ATP-dependent activity, gene expression, and cellular localization (Fig. S21). Overlap between stress-induced DMGs and DETs remained low (3 genes in RNAi-*dml* to 19 in WT), suggesting that most methylation changes act indirectly or modulate chromatin context rather than directly driving transcription. By contrast, 50–70% of line-specific DETs corresponded to DMGs (Fig. 7B), indicating tight methylation–transcription coupling when core methylation machinery is perturbed. In OX-*dml*, ON/OFF DET-DMGs (324 genes) were enriched in metabolic and epigenetic regulation categories, whereas quantitatively expressed DET-DMGs (701 genes) were associated with metabolic processes and ncRNA regulation (Fig. S22).

Focusing on stress-induced DETs shared by at least two epitypes identified 46 genes involved in transcriptional regulation, auxin response, and xylem development (Fig. 7C; Table S3). Three clusters emerged: (I) downregulated in WT, (II) upregulated in WT, and (III) uniquely regulated in modified lines. Notably, OX-*dml* and RNAi-*dml* frequently showed reversed expression patterns relative to WT, underscoring the impact of altered demethylation capacity.

Line-specific comparisons to WT identified 137 (RNAi-*ddm1*, control) to 1,999 (RNAi-*dml*, control) DETs (Fig. S13; Fig. S14). Among shared DETs, only 20 displayed opposite regulation (Fig. 7D). For instance, RAF24 (osmotic stress response) was upregulated in OX-*dml* but downregulated in RNAi-*ddm1*, while ILITHYIA showed opposite patterns between RNAi-*dml* and OX-*dml* under stress. Under drought, OX-*dml* exhibited a higher proportion of upregulated DETs, including three strongly induced genes of unknown function on chromosome 8.

No direct association was detected between stress-induced DMCs and differential splicing events (DSE) (Fig. S23). However, over 78% of genotype- or epitype-specific DSE overlapped with DMCs (Fig. S24A). GO enrichment identified “chromatin remodeling” as the top category, suggesting functional interplay between DNA methylation, chromatin dynamics, and alternative splicing in stress memory. As an illustrative example, Potri.019G052500 (314 DMCs) exhibited a switch from hypo- to hypermethylation at a spliced exon (Fig. S24C).

## Discussion

Previous studies in Populus demonstrated that water deficit induces persistent molecular changes in shoot apical meristems (SAMs) that remain detectable after one week of recovery (Lafon-Placette et al., 2018; Le Gac et al., 2018; Sow et al., 2021), in line with the concept of short-term somatic stress memory (Crisp et al., 2016; Hilker et al., 2016). Our work extends this perspective to the vascular cambium, the meristematic tissue responsible for radial growth and wood formation, a likely reservoir for long-term epigenetic imprinting (Maury et al., 2019). Most importantly, we identified a few molecular changes that persisted one year later in trees subjected to a second water deficit, suggesting a stable epigenetic memory that could prime future drought responses. Whether this trans-annual memory does confer priming remains to be elucidated, but the tissue specificity and molecular depth underscore its central role in orienting stress-induced acclimation in the vascular cambium.

### Cambium-derived tissues retain drought-associated molecular signatures consistent with a short-term somatic memory

Prolonged exposure to five weeks of water deficit significantly altered tree carbon and water relations, with reduced stomatal conductance, increased water-use efficiency, and decreased primary and secondary growth, without complete growth arrest. Stress intensity was moderate, allowing recovery within one week of rewatering.

Hormonal profiles retained clear stress imprints. Drought decreased ABA, JA, and IAA, while increasing brassinosteroids (BRs) and GA₁₂ across genotypes and epitypes. The persistence of reduced ABA, a central stress hormone, indicated growth recovery rather than ongoing stress signaling (Kuromori et al., 2018; Tong et al., 2014), consistent with short-term post-stress memory (Crisp et al., 2016; Hilker et al., 2016). Because hormone quantification likely integrated xylem and phloem derivatives, these values reflect a composite vascular signature rather than a strictly local meristematic signal. Nonetheless, the hormonal patterns aligned with cambial growth regulation: IAA promotes xylem differentiation (Ruzicka et al., 2015; Smetana et al., 2019), cytokinins stimulate phloem identity (Mähönen et al., 2006), and BRs and GAs synergize with auxin to enhance xylem expansion (Planas-Riverola et al., 2019), whereas ABA and JA restrict cambial activity (Fukuda, 2004; Nieminen et al., 2008).

Transcriptomic analysis revealed selective reprogramming after rewatering. Only ∼0.2% of transcripts (200–250 genes) were differentially expressed, contrasting with the extensive responses typical of acute drought (Cohen et al., 2010; Wilkins et al., 2009), but consistent with post-stress responses previously reported in poplar SAMs (Lafon-Placette et al., 2018; Sow et al., 2021). DETs were enriched in cell wall organization, hormone pathways, and signaling processes, supporting active growth-related remodeling during recovery. Their persistence one week after rewatering further supports short-term molecular memory (Crisp et al., 2016; Hilker et al., 2016).

At the epigenetic level, thousands of DMCs were detected across CG, CHG, and CHH contexts, with genotype- and epitype-specific patterns (see below). CG and CHG methylation are generally associated with stable expression states, whereas CHH methylation is more dynamic and often TE-linked (Dowen et al., 2012; Talarico et al., 2024). The maintenance of these methylation signatures after rewatering suggests they extend beyond transient stress footprints and may contribute to short-term somatic memory. Although only a small subset of DETs (≤22) overlapped with nearby DMCs, these included functionally relevant genes such as ERD family members (Alves et al., 2011) and UBP13, which modulates MYC2 levels in jasmonate signaling (Zhang et al., 2024). Despite limited numerical overlap, this convergence supports coordinated epigenetic–transcriptional regulation during recovery.

Overall, our data indicate that the vascular cambium integrates prior drought exposure through persistent, multilayered adjustments, establishing a molecular basis for short-term somatic memory. Future work combining temporal and cell-type–resolved omics with hormone localization will be required to resolve causal links between hormonal dynamics and epigenetic reprogramming.

### Genetic and epigenetic backgrounds both drive molecular short-term memory

Short-term memory varied across *P. nigra* genotypes and *P. tremula × P. alba* epitypes, indicating that both genetic and epigenetic variation shape drought responses. In *P. nigra*, PG-31 displayed higher drought tolerance and sustained growth compared with DRA-038. Yet their molecular strategies diverged: DRA-038 showed stronger hormonal and transcriptional shifts, whereas PG-31 exhibited fewer transcriptional changes but markedly more DMCs. Increased GA12 and cytokinins, together with elevated DPA in DRA-038, suggested cambial reactivation with active ABA turnover (Ding et al., 2024; Wybouw et al., 2024). However, reduced UBP13 and increased SRO1 expression (Park et al., 2022; Luo et al., 2022; Xiong et al., 2022; Jaspers et al., 2010; Teotia et al., 2009; Teotia & Lamb, 2011) indicated a restrained recovery prioritizing meristem integrity, consistent with the “growth–survival dilemma” (Claeys & Inzé, 2013). Thus, DRA-038 followed a conservative, transcriptionally plastic strategy, whereas PG-31 relied more heavily on epigenetically stabilized regulation. Such divergence likely reflects population-specific epigenomic landscapes shaped by evolutionary history (Vanden Broeck et al., 2024).

In contrast, *P. tremula × P. alba* epitypes differed only in targeted methylation machinery modifications yet displayed distinct drought phenotypes. OX-*dml* combined improved growth with broader transcriptional activation and elevated GAs, consistent with release from methylation-mediated repression (Xie et al., 2025). Its altered mycorrhization capacity (Vigneaud et al., 2023) further supports a systemic role for active demethylation. Upregulated genes involved auxin transport, water deprivation responses, and xylem development (Wybouw et al., 2024). RAF24, both overexpressed and differentially methylated, integrates chromatin remodeling and osmotic stress responses (Fàbregas et al., 2020; Zhou et al., 2025). RAF24 emerges as a candidate regulator of somatic memory, pending functional validation, while repression of ILLYTHIA suggested resource allocation away from immune regulation toward growth. Collectively, these data indicate that active demethylation reshapes regulatory networks and facilitates epigenetic priming (Zhu et al., 2025; Lin et al., 2020).

Methylation landscapes differed markedly between backgrounds. In *P. nigra*, stress-induced DMCs predominantly occurred in genic CG sites, consistent with mitotically stable modifications. In *P. tremula × P. alba*, stress-responsive DMCs were enriched in CHH contexts within transposable elements (Liang et al., 2014; Sow et al., 2021), whereas many epitype-specific DMCs still occurred in genic CGs. CHH methylation, deposited via RdDM (DRM2) or CMT2 (Stroud et al., 2014; Xie et al., 2025; Xu & Law, 2024), represses stress-activated TEs (Ramakrishnan et al., 2022), suggesting a rapid genome-stabilizing response to drought-induced chromatin perturbation. CHH levels increased in RNAi-*dml* but decreased in OX-*dml* and RNAi-*ddm1*, consistent with antagonism between DML activity and RdDM (Sow et al., 2021; Xu & Law, 2024). However, a truncated CMT2 allele was identified exclusively in our INRA 717-1B4 clone (Fig. S25), previously linked to CHH variability and stress tolerance in *Arabidopsis* (Shen et al., 2014b), provides a plausible genetic basis for enhanced CHH plasticity. Functional validation will be required to confirm causality.

Overlap among stress-induced DETs, DSEs, and DMCs was largely genotype- or epitype-specific. More than 70% of genotype- and epitype-specific splicing events co-localized with DMCs in chromatin remodeling genes (López Sánchez et al., 2016; Wang et al., 2016; Zhang et al., 2020), although few overlapped directly with DETs. Given that DNA methylation operates through both cis and trans mechanisms, limited direct correspondence does not preclude functional coupling.

Altogether, our data support a model in which somatic memory arises from methylation contexts shaped by genetic predisposition and epitype-specific regulation, through multilayered interactions among hormones, transcription (including splicing), and DNA methylation. Determining whether this memory enhances resilience or reflects adaptive trade-offs remains central to understanding perennial plasticity under recurrent climatic stress.

### Somatic stress memory lasts more than a year and may contribute to priming

Recent models propose that plants retain stress experiences through multi-layered imprints involving transcription and epigenetics, forming a persistent molecular memory (Farkas & Dobránszki, 2024). In long-lived perennials, such memory may shape responses across developmental phases and seasons (Gallusci et al., 2023). Our multi-year experiment in *P. nigra* provides empirical support for this hypothesis.

S/C (Stress Year 1 / Control Year 2) trees of the more sensitive genotype DRA-038 showed reduced growth compared to C/C controls in Year 2, suggesting the persistence of a physiological debt, whereas S/C trees of the more tolerant genotype PG-31 were unaffected. In contrast, primed trees subjected to a second water deficit (S/S) showed comparable growth to unprimed trees (C/S) in Year 2, indicating a potential stress-induced acclimation especially in DRA-038. Interestingly, although very different in Year 1, hormonal profiles of primed DRA-038 trees clearly converged toward those of PG-31 in Year 2, with higher auxin and GA and lower cytokinins, possibly favoring xylem differentiation and contributing to partial growth compensation. Differential growth and hormonal patterns were further accompanied by distinct molecular profiles. DETs observed between C/S and S/S trees in Year 2 could be assigned to two distinct categories: type I (sustained from Year 1) and type II (enhanced upon re-exposure to water deficit), which was consistent with the definitions of stress memory in annuals (according to Bäurle & Trindade, 2020). We found six memory DETs in the genotype DRA-038, including reversed expression between Year 1 and Year 2 of *UBP13* and *SRO1*, suggesting a shift from conservative to growth-oriented strategies (see above discussion). Alternatively, we only found two memory DETs in the genotype PG-31, indicating molecular stability rather than flexibility. The limited number of candidates likely reflects the distinction between chromatin-based cis memory and trans memory mediated by transcription factors or hormones (Farkas & Dobránszki, 2024).

At the methylome level, Year 2 saw a strong increase in CG methylation consistent with their role in stable memory (Zhang et al., 2018). Most importantly, DRA-038 accumulated more than 20-fold more DMCs than PG-31, converging toward PG-31 Year 1 levels and mirroring hormonal patterns. In addition, the number of DETs colocalizing with DMCs (including *UBP13*) also clearly increased in Year 2, suggesting a possible role for epigenetic regulation of transcriptional long-term memory. Altogether, our data provide empirical evidence that cambial somatic memory is multi-layered (Groover, 2025) and genotype-dependent. Importantly, since trees were cut back and regenerated (stem) between years, our findings also demonstrate that this memory persisted over annual timescales through cell division (Gallusci et al., 2023). The higher molecular plasticity observed in the intrinsically more sensitive genotype DRA-038 contrasted with the higher stability observed in the intrinsically more tolerant genotype PG-31 and likely reflected distinct adaptive strategies. This supports the concept of epigenetic priming where previous stress history may enhance future responsiveness.

## Conclusion

Our study demonstrates that the vascular cambium and its derivatives retain molecular imprints of past drought across growing seasons, establishing trans-annual somatic memory through coordinated hormonal, transcriptional, and epigenetic remodeling. These signatures reflect active reprogramming rather than passive persistence and vary with genotype. Moreover, epitype-specific responses indicate that DNA methylation itself contributes to stress memory, likely through combined cis- and trans-mediated effects that propagate across molecular layers.

Unlike the transient cambium of *Arabidopsis*, the long-lived tree cambium integrates environmental cues over years, providing a unique window into durable molecular memory. Meristematic cells are particularly suited to store stress signatures through hormonal shifts, transcriptional memory, and mitotically stable CG methylation marks (Maury et al., 2019). Consistent with this, most cambial DMC-containing genes (94.5%) overlap with methylation-responsive genes previously reported in poplar meristems (Fig. S26; Lafon-Placette et al., 2018; Sow et al., 2021, 2023), reinforcing the functional relevance of these epigenetic changes. While histone-based mechanisms often dominate short-term priming in annuals, our results support DNA methylation, particularly in CG context, as a robust substrate for long-term stress memory in trees (Le Gac et al., 2018; Trontin et al., 2024).

These findings suggest that post-drought recovery in perennials involves memory-mediated remodeling rather than a simple return to baseline (Gessler et al., 2020). Whether such memory enhances resilience or entails physiological costs (Backhaus et al., 2014) remains unresolved but is central to redefining resilience and plasticity in forests facing recurrent drought. Future integration of temporal multi-omics, gene-network reconstruction, and targeted gene perturbation (e.g. CRISPR–Cas9) will be required to establish causal regulatory pathways underlying drought-induced somatic memory.

## Acknowledgements

This work benefited from the equipment and services coordinated by Y. Marie for the RNA-seq analyses conducted at the IGenSeq platform (ICM, Paris, France). Scientific and technical support was provided by the Epigenomic Environmental Platform of IHPE (http://ihpe.univperp.fr/plateforme-epigenetique/; Cristian Chapparo, Perpignan, France). We thank the GENOTOUL platform for providing computational storage space. Isotopic and elemental analyses were performed by the SILVATECH platform (Silvatech, INRAE, 2018; Structural and Functional Analysis of Tree and Wood Facility, DOI: 10.15454/1.5572400113627854E12), involving UMR 1434 SILVA, 1136 IAM, 1138 BEF, and EA 4370 LERMAB at the INRAE Grand-Est Nancy research center. The SILVATECH facility is supported by the French National Research Agency (ANR) through the Laboratory of Excellence ARBRE (ANR-11-LABX-0002-01). We would also like to thank the Tree Cell Engineering Laboratory (LICA) (https://www6.inrae.fr/in-sylva-france_eng/Services/In-Lab/LICA), where the GM plants were grown and characterized. Finally, we are grateful to the GBFOR facility (INRAE, 2018; Forest Genetics and Biomass Facility, DOI: 10.15454/1.5572308287502317E12) for its contribution to the experimental design setup and sample collection. We also acknowledge the GDR 3E network (Epigenetics in Ecology and Evolution, coordinated by Christoph Grunau) for stimulating scientific exchanges, and the members and coordinators (including S.M.) of the COST Actions Epigenetic mechanisms of Crop Adaptation To Climate cHange (EPICATCH, CA19125) and Innovative Woody Plant Cloning (COPYTREE, CA21157) for their constructive discussions. We thank Professor Steven H. Strauss and his team (Department of Forest Ecosystems and Society, Oregon State University, Corvallis, Oregon, USA) for providing the poplar RNAi-*ddm1* lines. We are grateful to Julien Vigneaud for valuable discussions related to the EPITREE project, and to Vanina Benoit and Corinne Buret (UMR BioForA, Orléans) for their assistance with sample collection.

## Author Contributions

S.M. and R.F. coordinated the research. The plant experimental design was established by V.S., I.A., S.S., R.F. and S.M. Plant growth and phenotypic assessments were performed by G.L., A.D., I.L.J., P.P under the supervision of R.F. Hormones analyses by LC-MS were done by Y-Q.F and B-D.C. Hydraulic analyses were done by R.F., G.L., I.L.J. and H.C. Histochemical and microscopic analysis were done by F.L. and R.F. Molecular analyses were performed by A.D., M.D.S. Statistics were performed by A.D. and R.F. Methylome sequencing was performed by J.T. and C.D. Bioinformatic analyses were done by A.D., A.T., O.R., and S.M.. The draft manuscript was conceived and written by A.D., R.F. and S.M., and further revised by H.D., V.J., L.S., J.T., V.S. All authors approved the final version of the article. We would like to pay a warm tribute to our co-author and friend Alain Delaunay, who passed away in August 2024. He joined the P2e lab (formerly LBLGC) of the University of Orléans (FRANCE) in 2000 and did outstanding work on epigenetic analysis in trees for several (inter)national projects and publications.

## Data availability

The data that support the findings of this study are openly available: The WGBS data have been deposited in the Sequence Read Archive (SRA) at https://www.ncbi.nlm.nih.gov/sra/PRJNA913022/ reference no. PRJNA913022; The RNA-Seq data have been deposited in the SRA at https://www.ncbi.nlm.nih.gov/sra/PRJNA913022/ reference no. PRJNA913022. The other data described in this Data note can be freely and openly accessed on the Recherche Data Gouv repository under: https://doi.org/10.57745/QZZ2WQ.

---

**Fig. S1.**
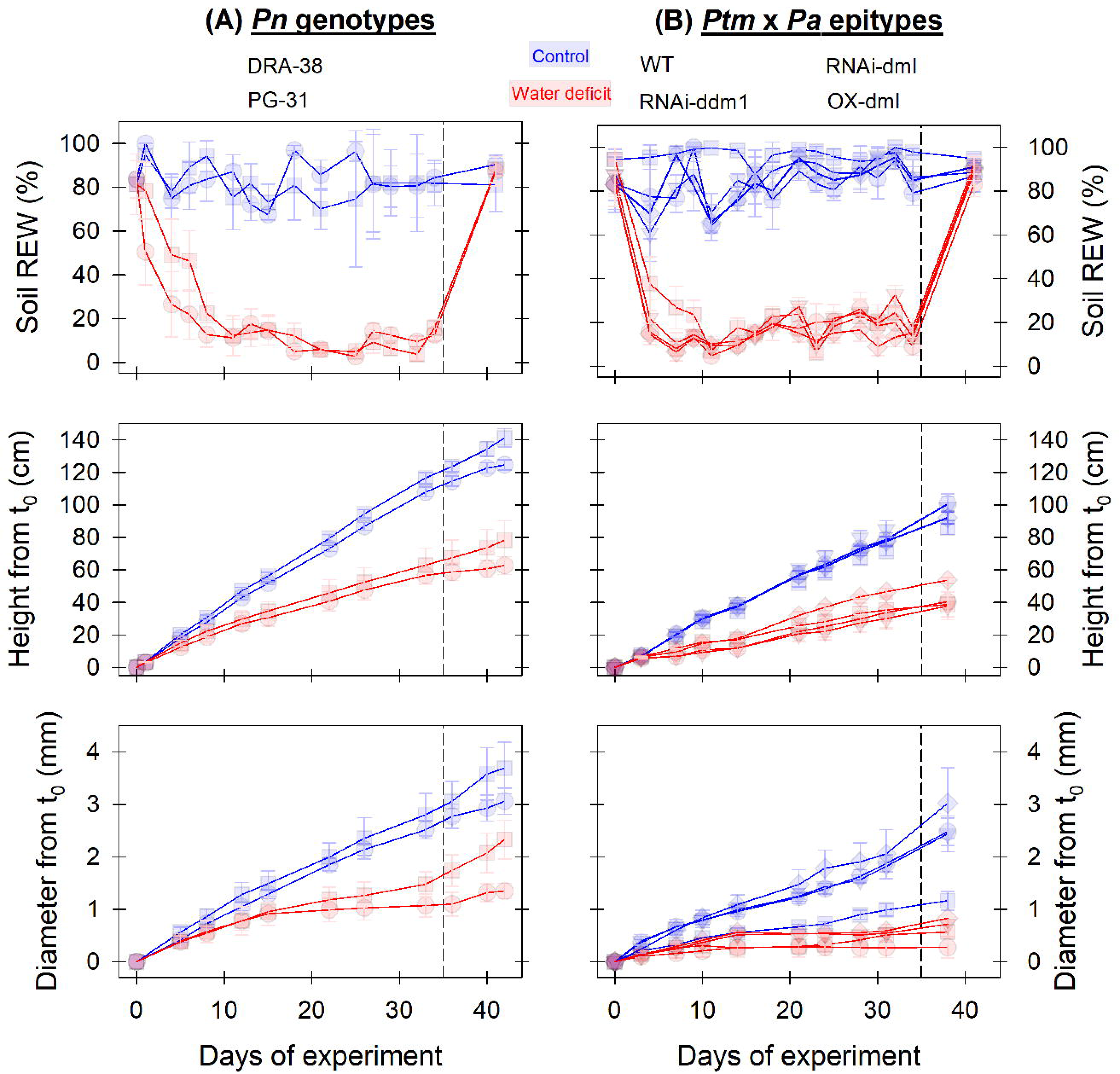
Soil water status and growth dynamics during drought treatment.

**Fig. S2.**
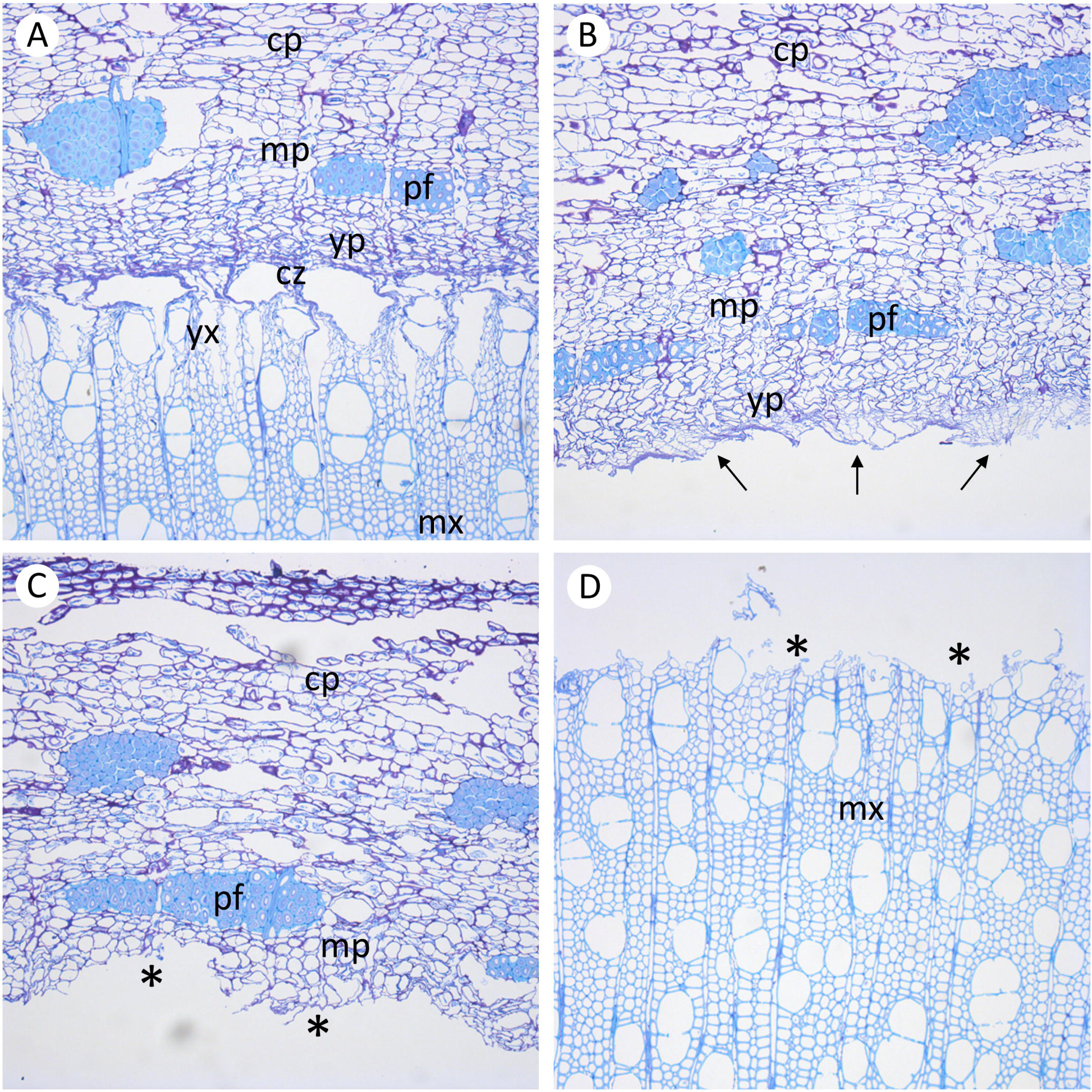
Histological characterization of cambial sampling.

**Fig. S3.**
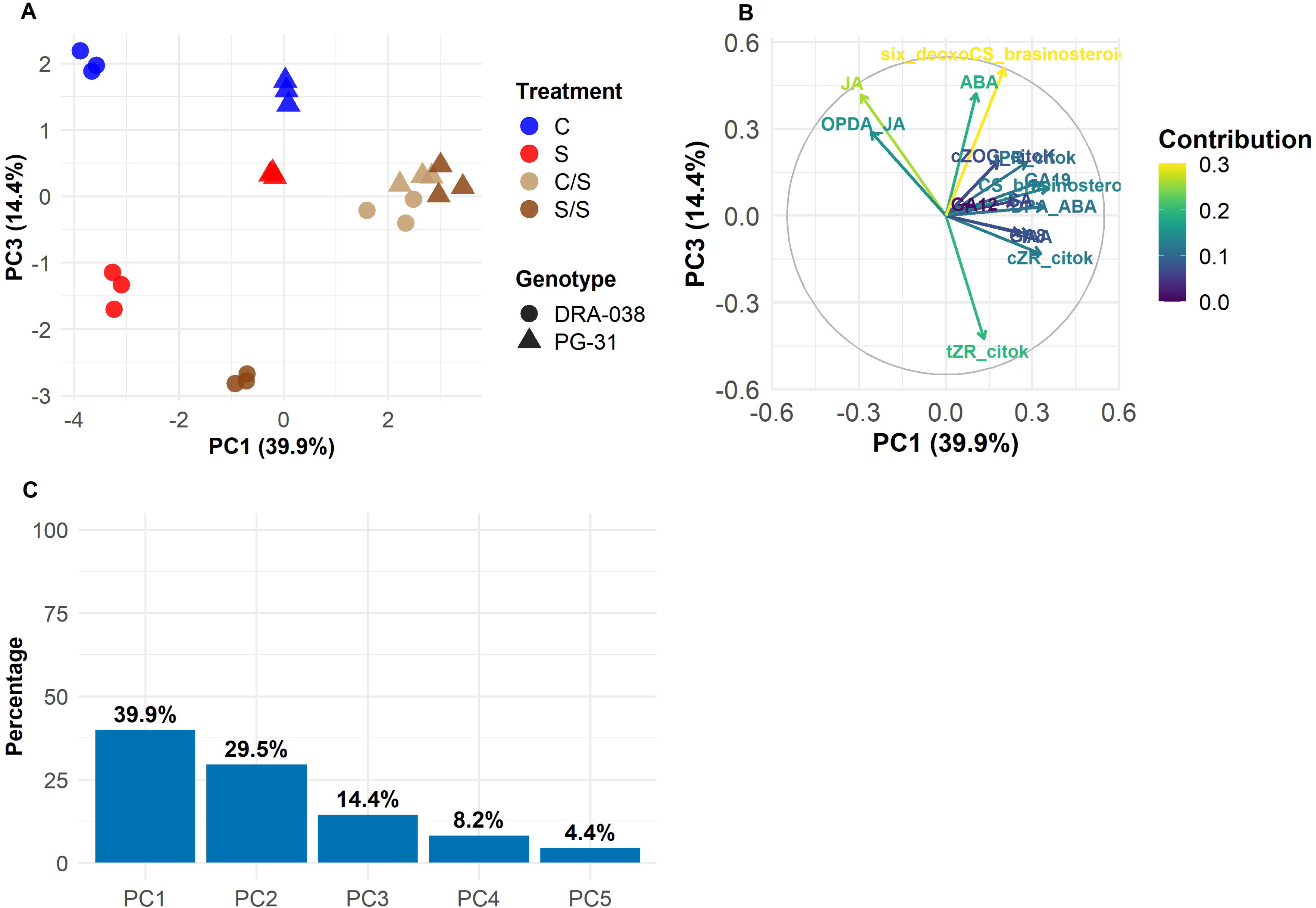
PCA of phytohormones in P. nigra genotypes.

**Fig. S4.**
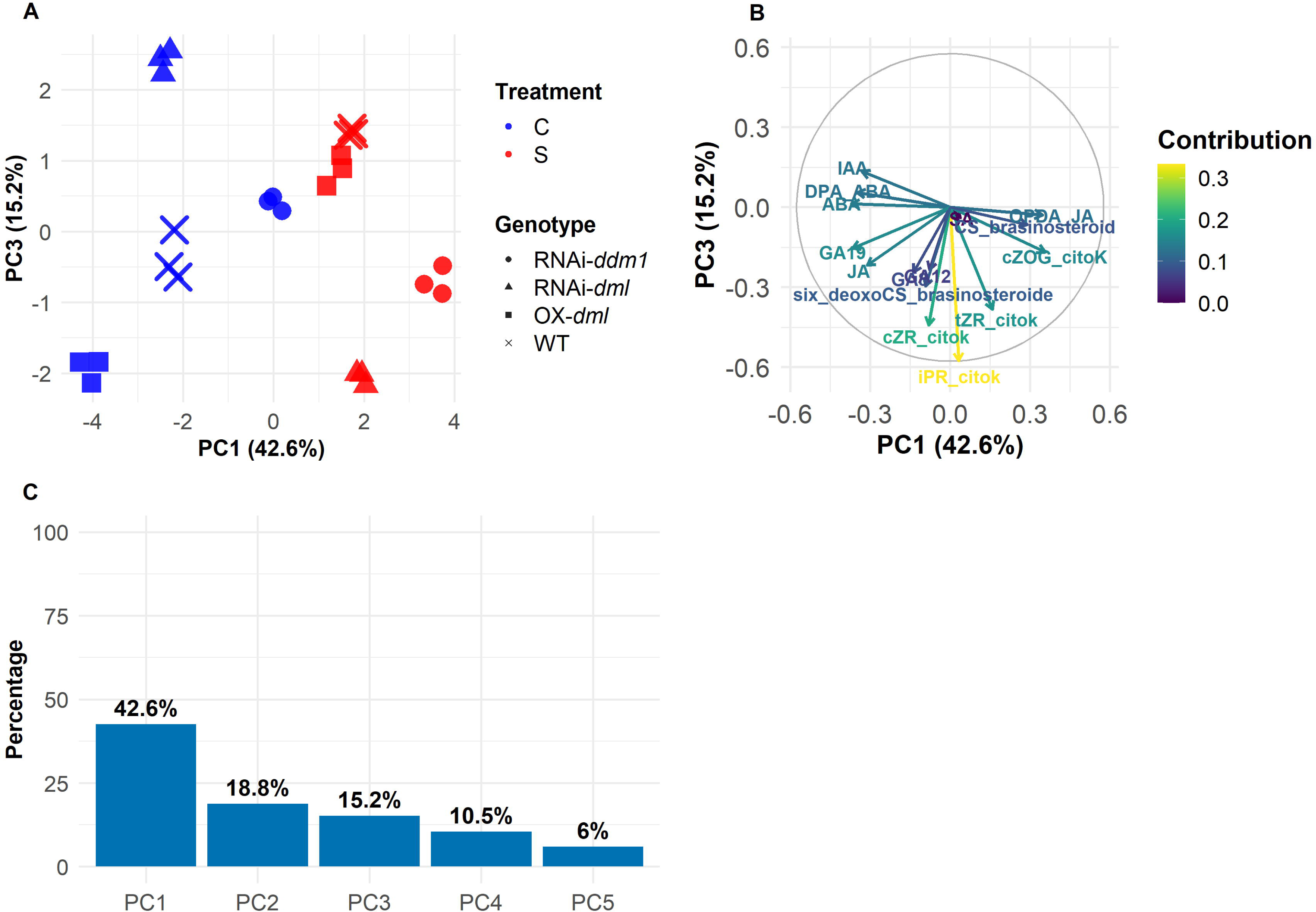
PCA of phytohormones in P. tremula × P. alba epitypes.

**Fig. S5.**
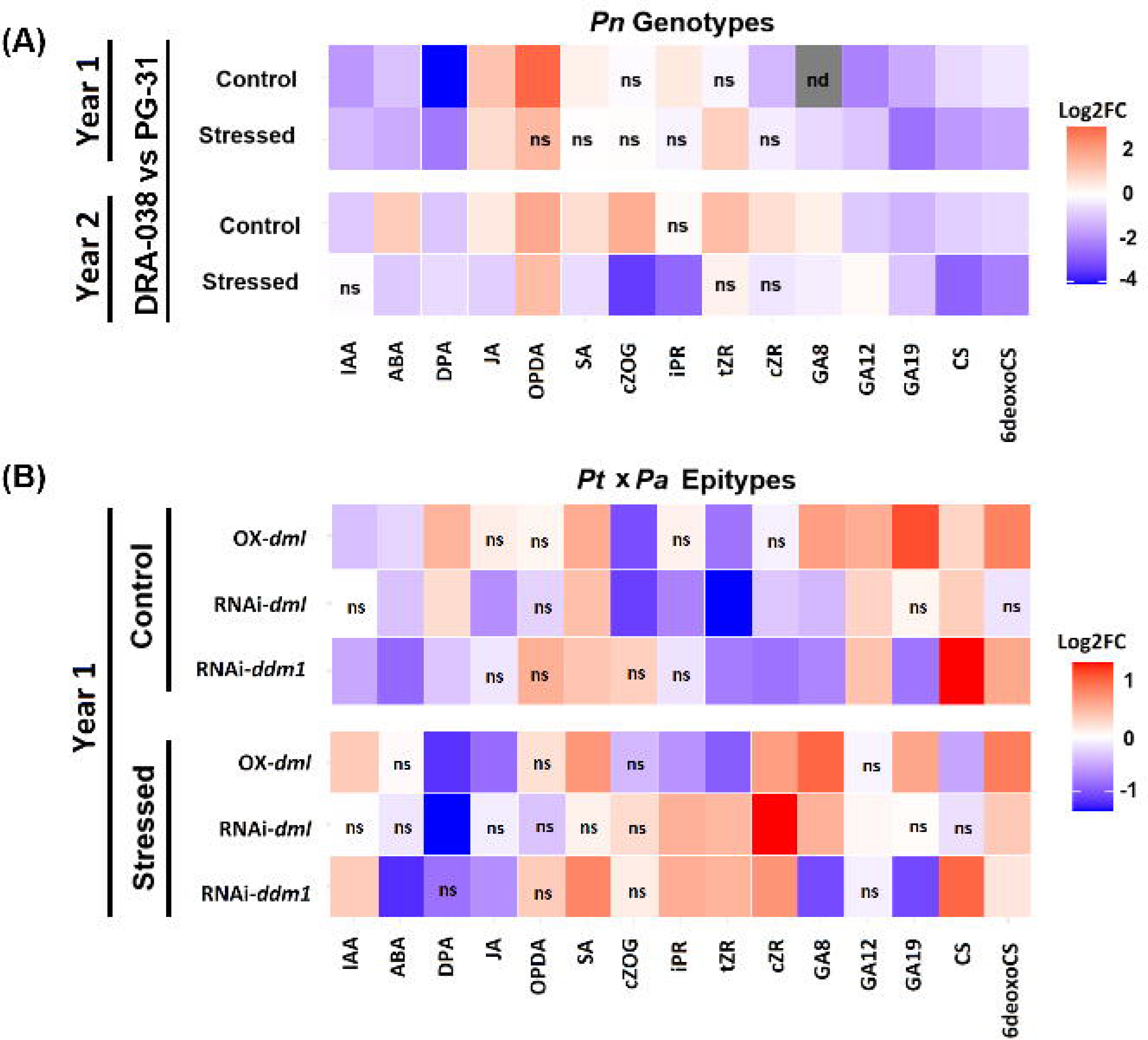
Hormone variation heatmap.

**Fig. S6.**
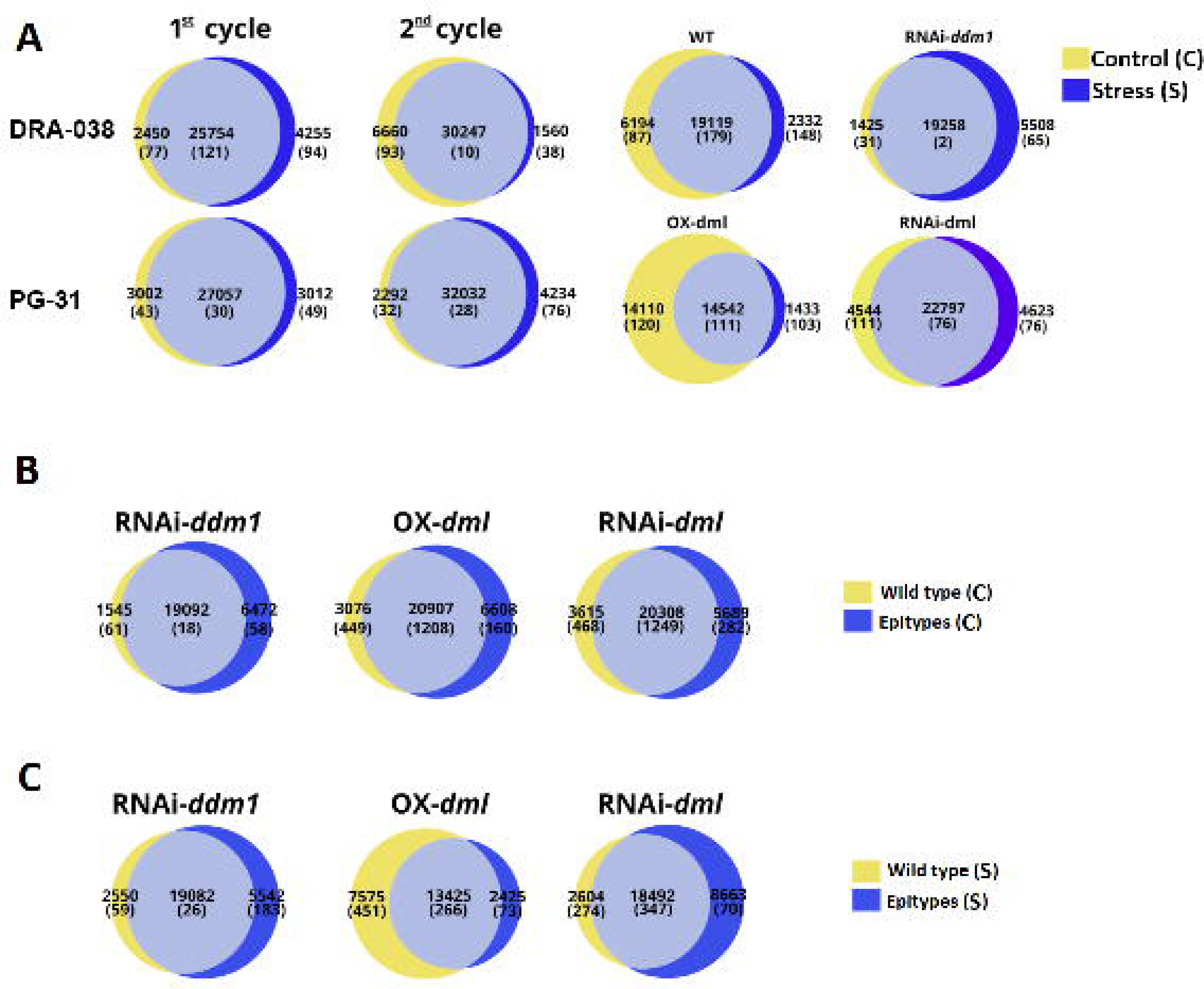
Transcript detection and differential expression overview.

**Fig. S7.**
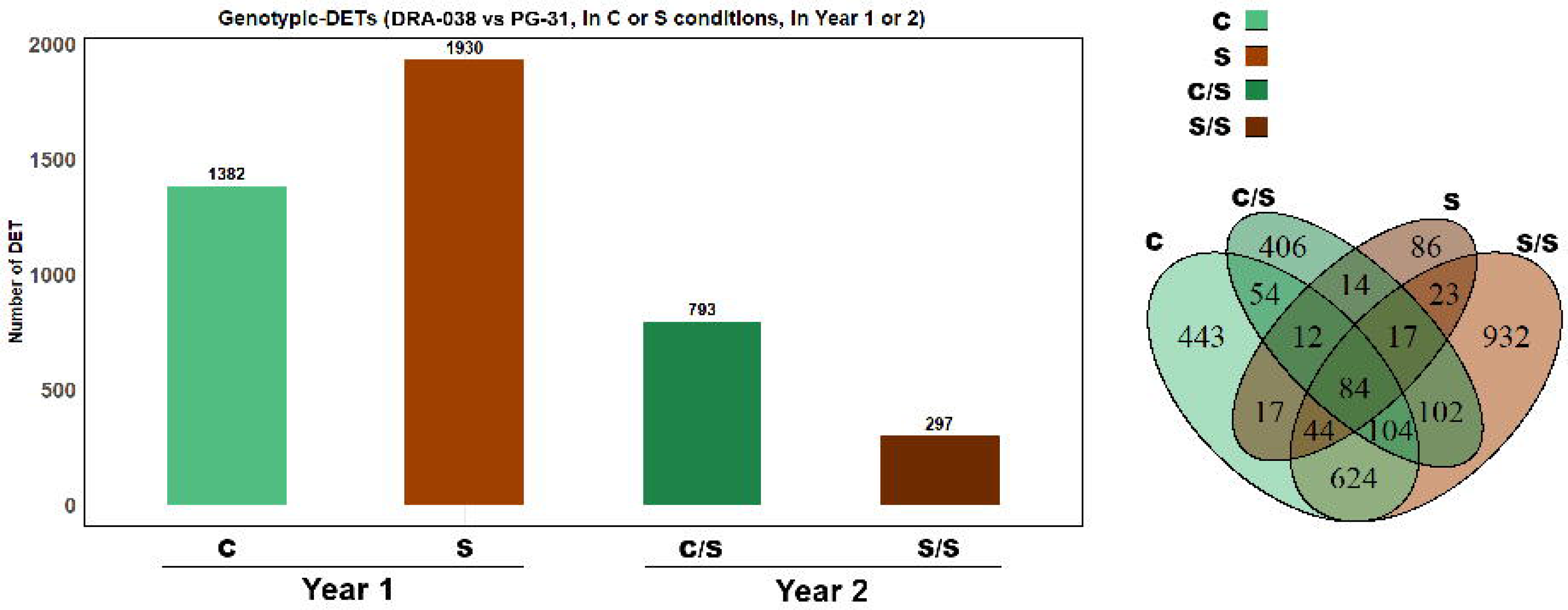
DET overlap across years in P. nigra.

**Fig. S8.**
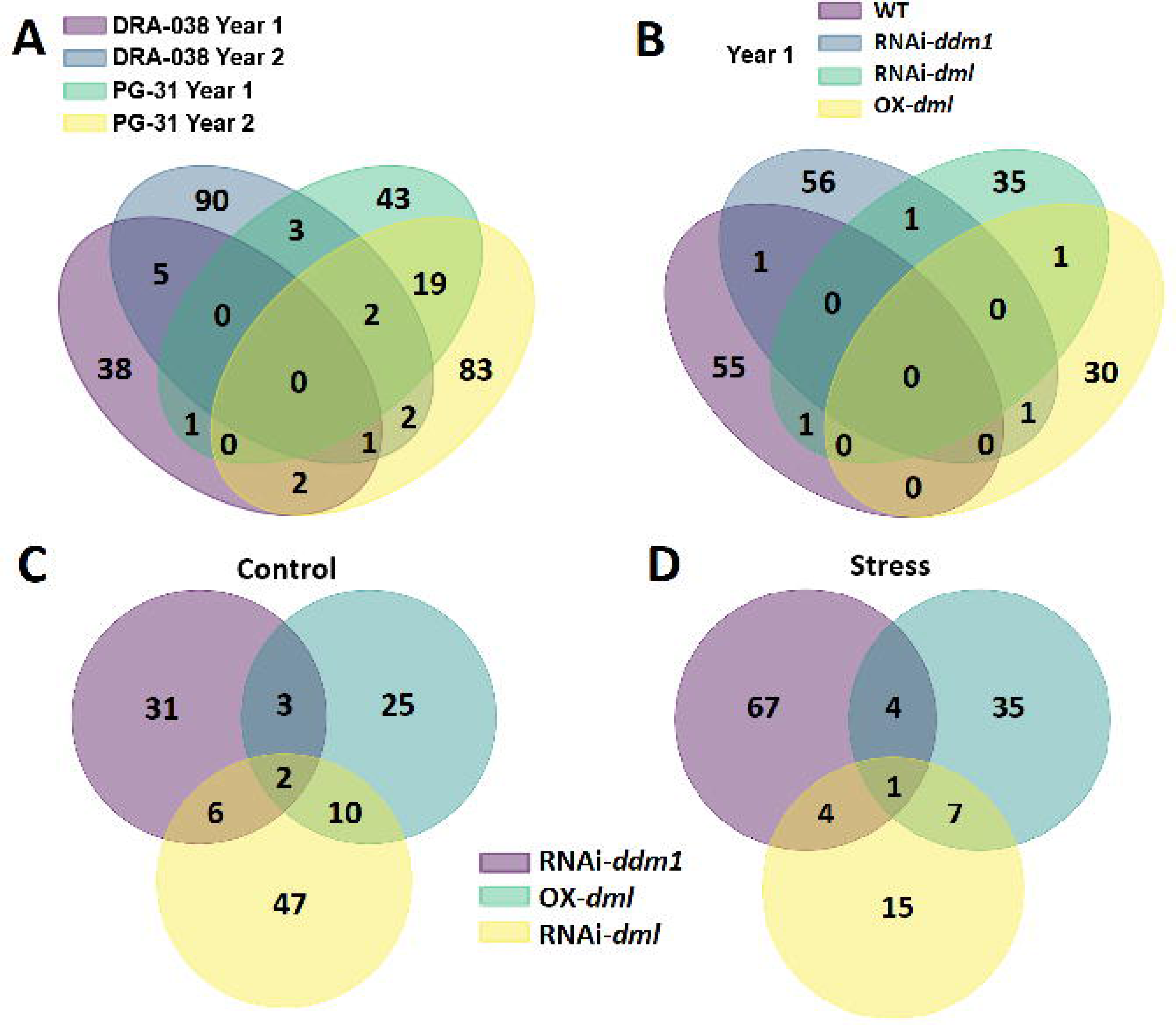
DET overlap across genotypes and epitypes.

**Fig. S9.**
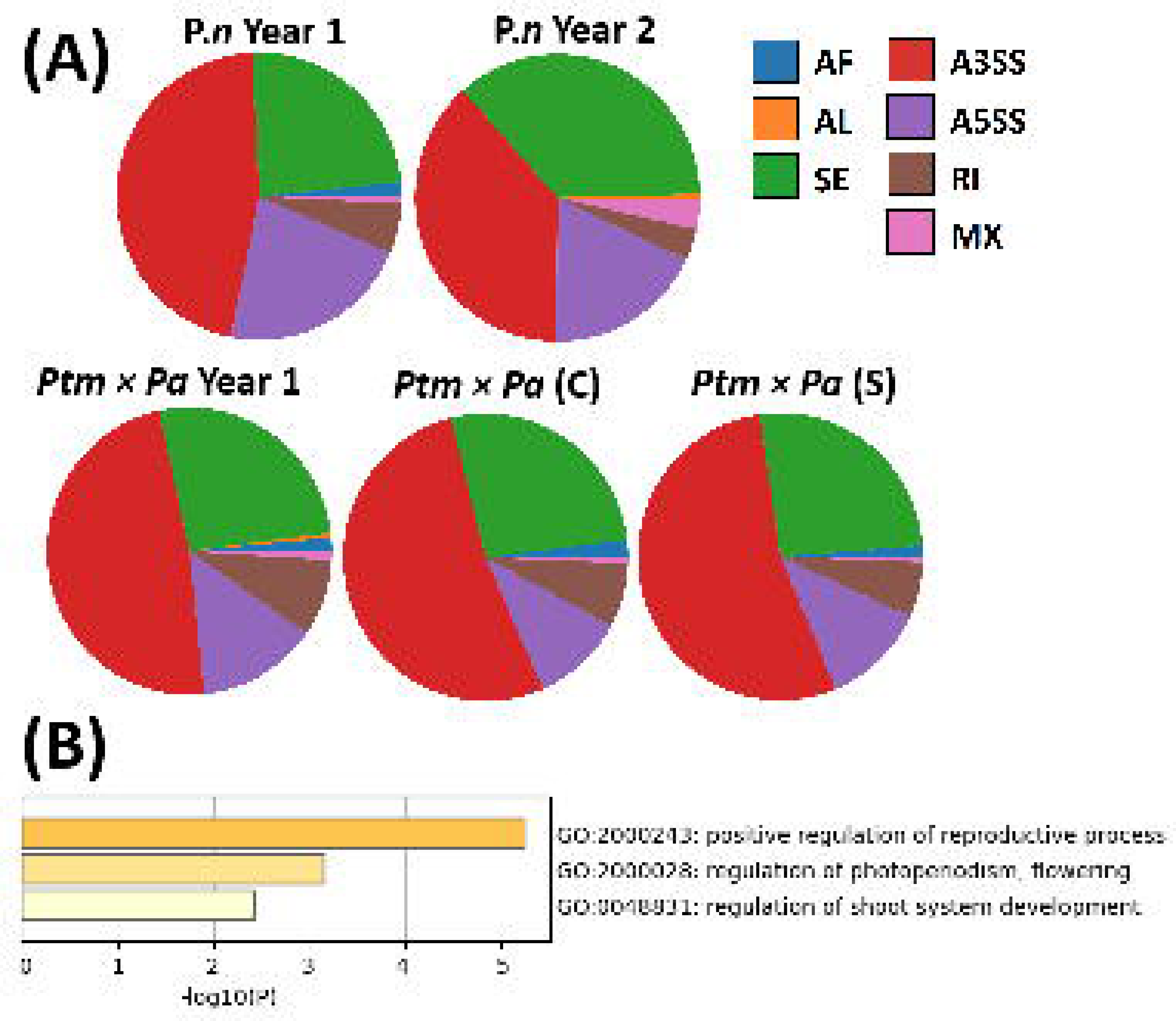
Distribution and functional enrichment of differential splicing events.

**Fig. S10.**
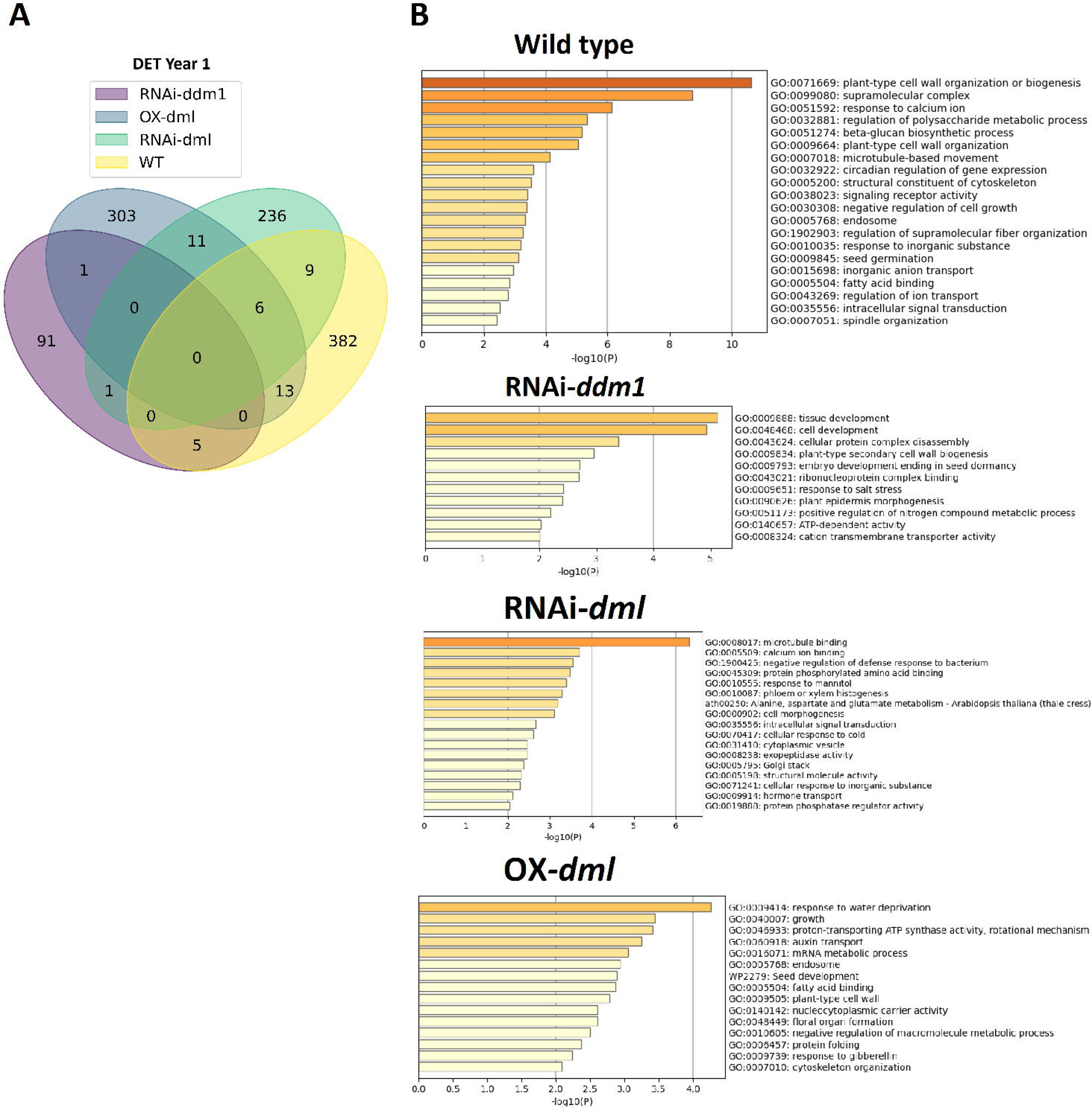
DETs and GO enrichment in epitypes (Year 1).

**Fig. S11.**
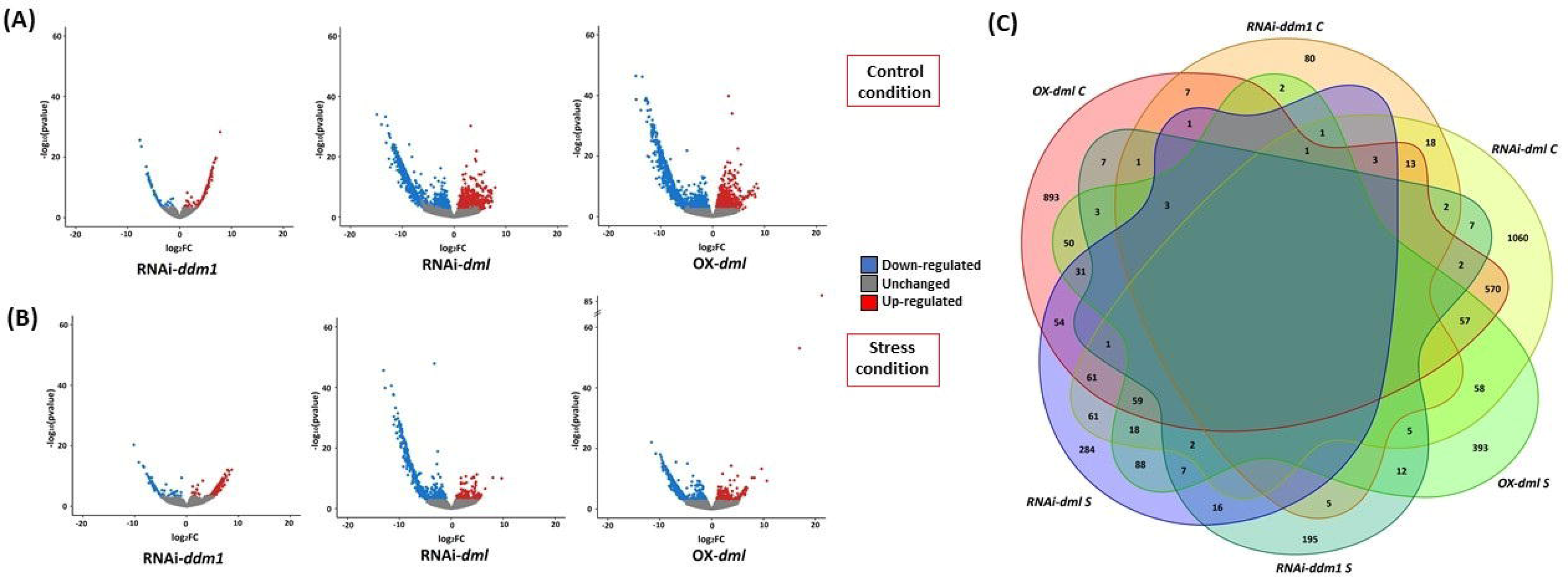
Differential gene expression in epitypes.

**Fig. S12.**
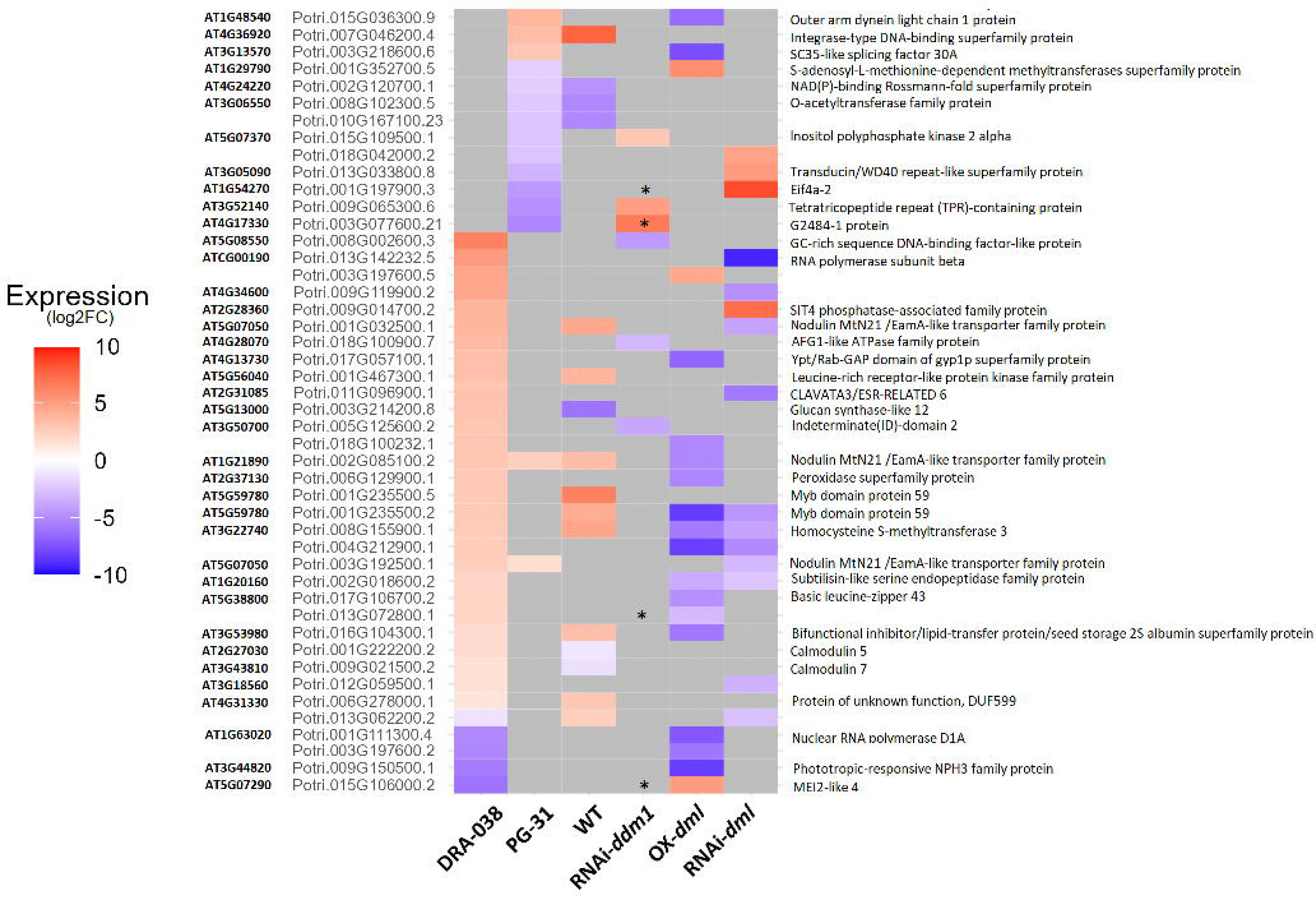
Heatmap of DETs across genotypes and epitypes.

**Fig. S13.**
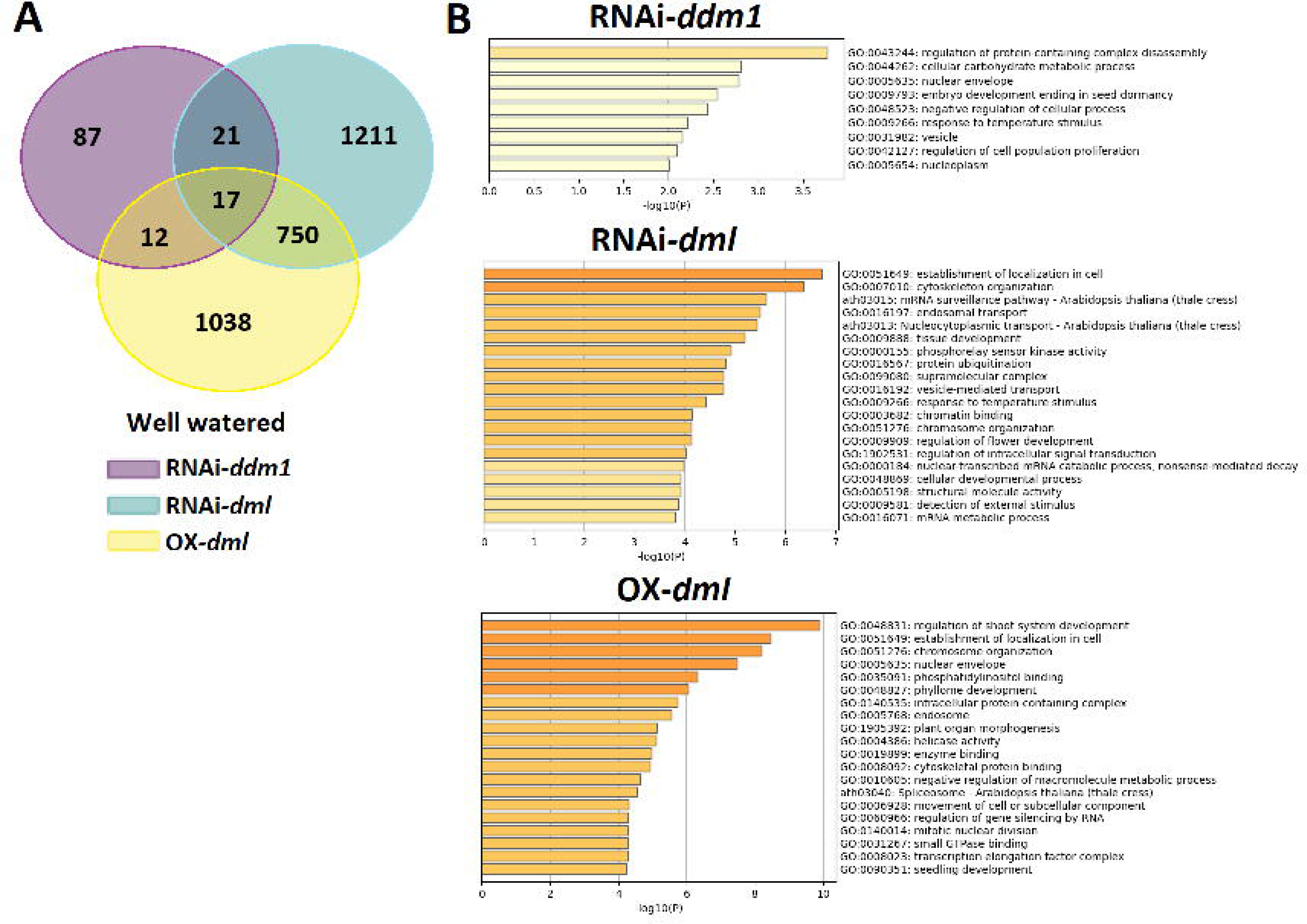
DETs and GO enrichment in epitypes under control conditions.

**Fig. S14.**
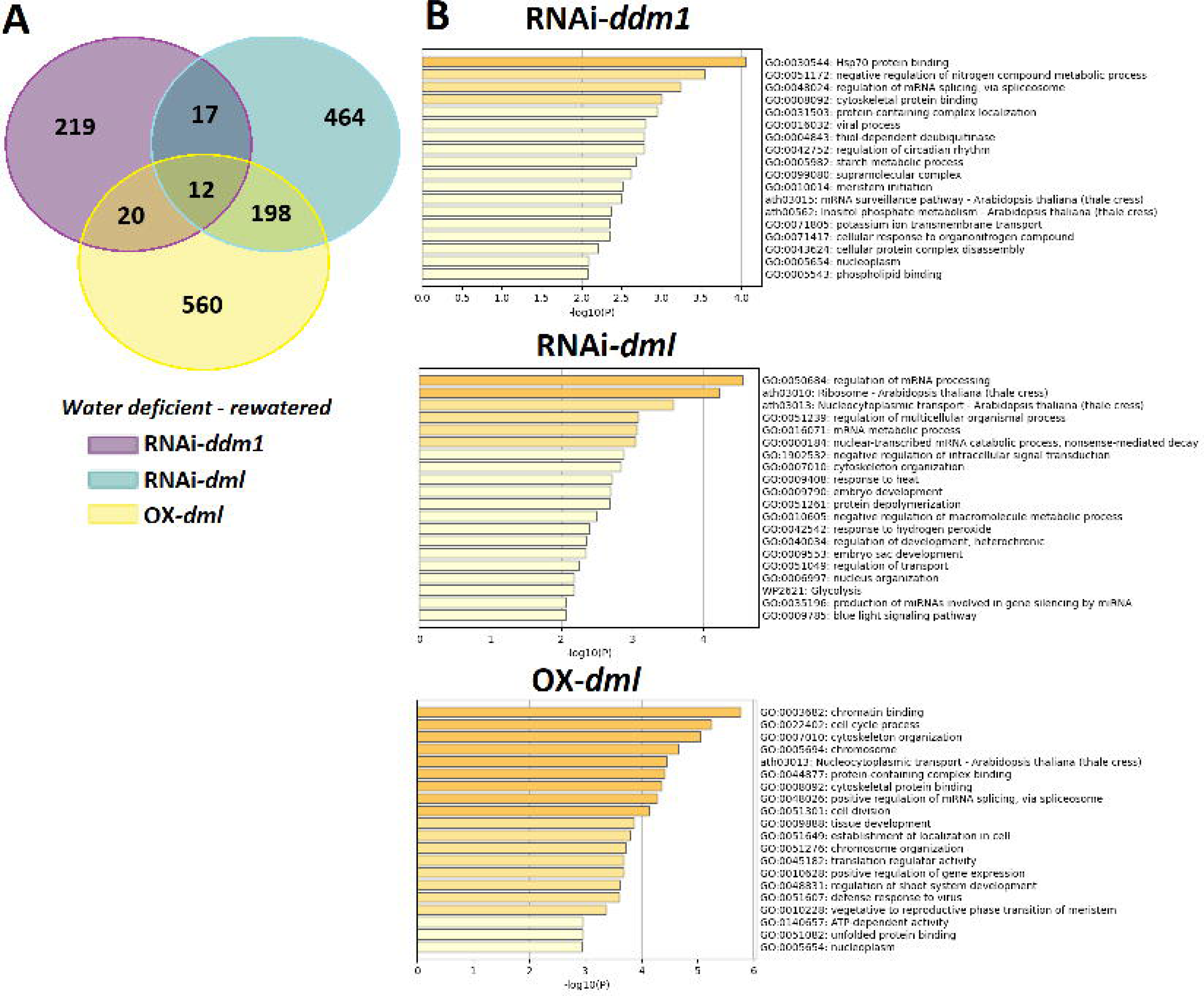
DETs and GO enrichment in epitypes under stress conditions.

**Fig. S15.**
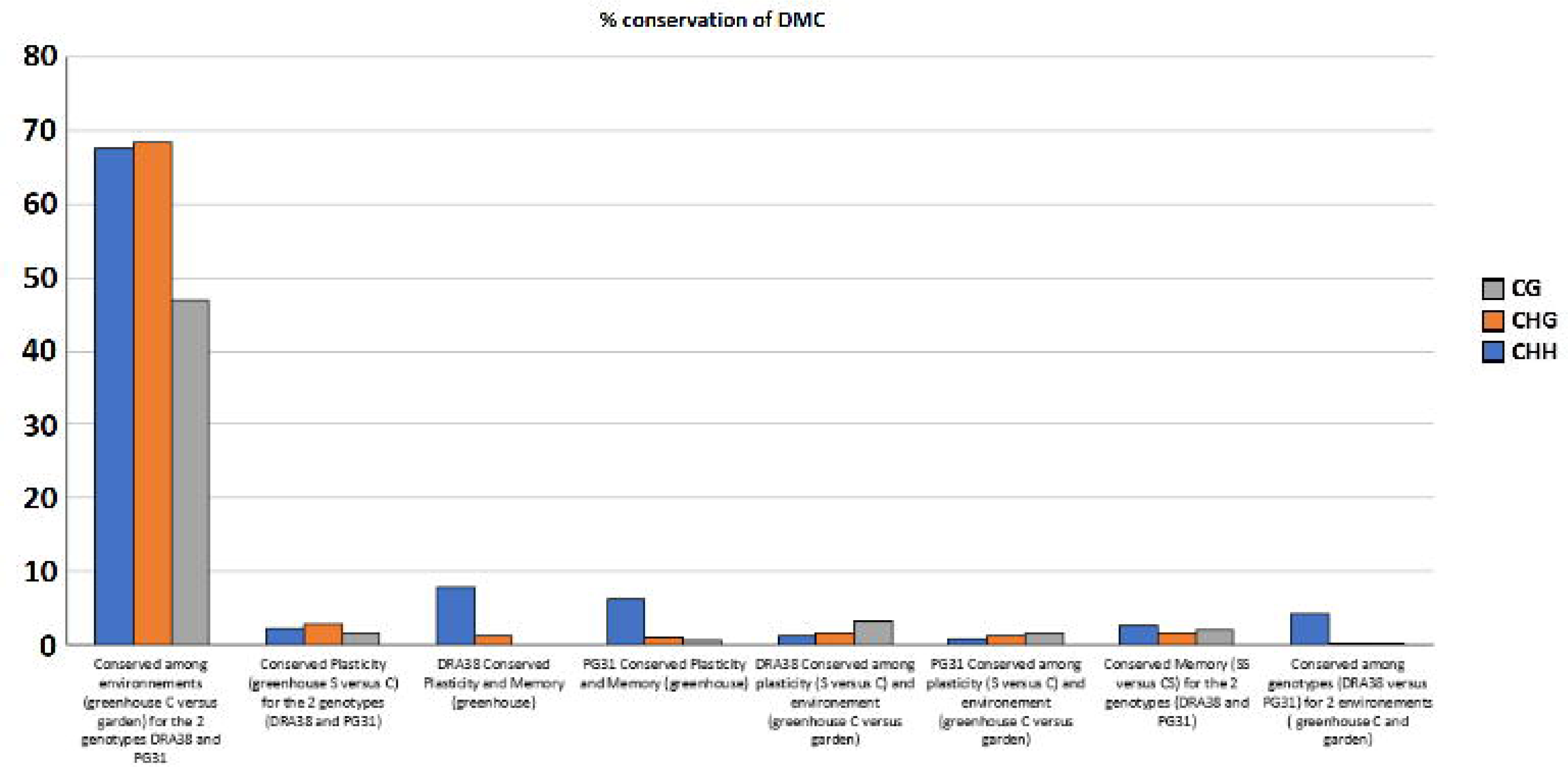
Conservation of DMCs across contexts and environments.

**Fig. S16.**
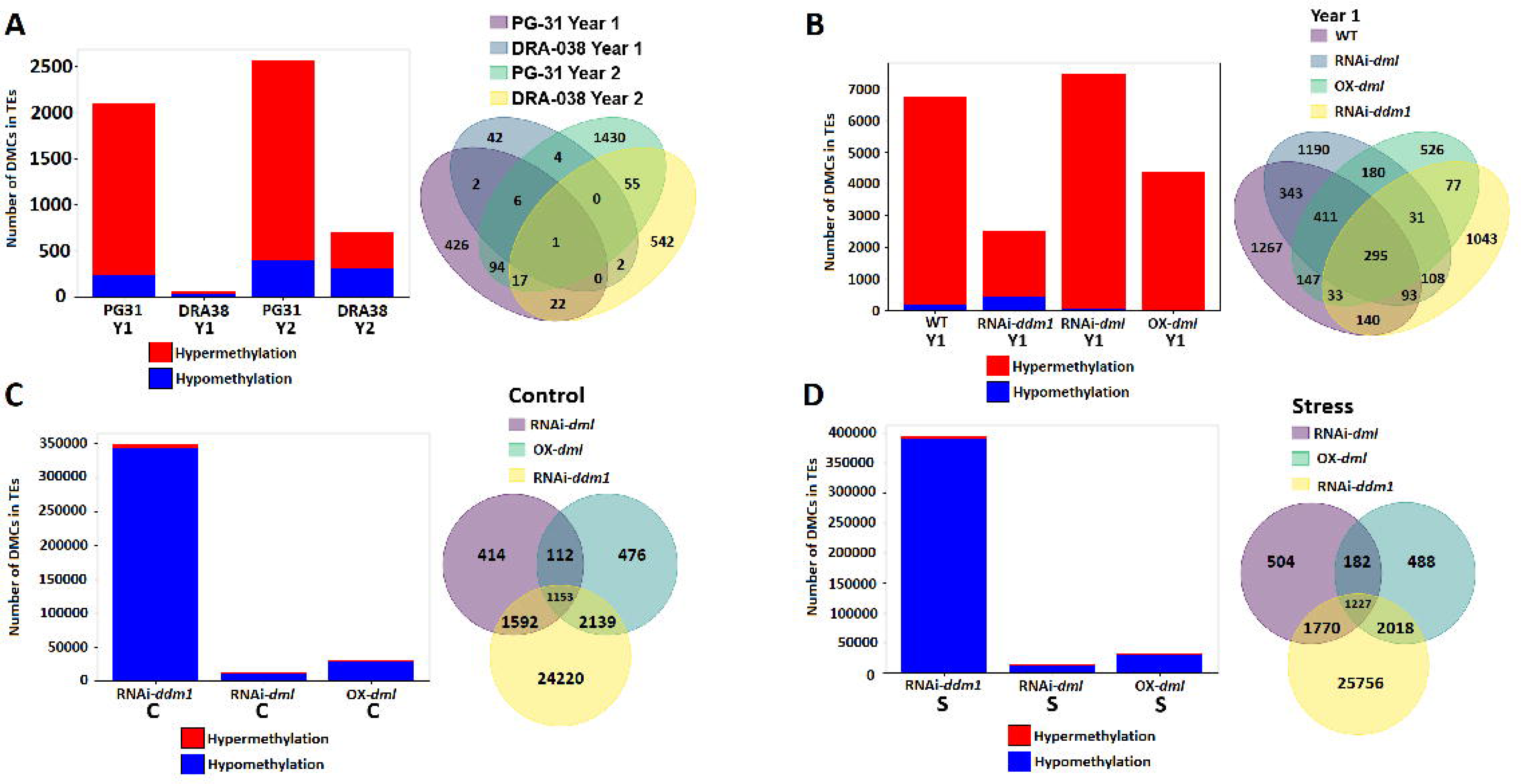
Differentially methylated transposable elements.

**Fig. S17.**
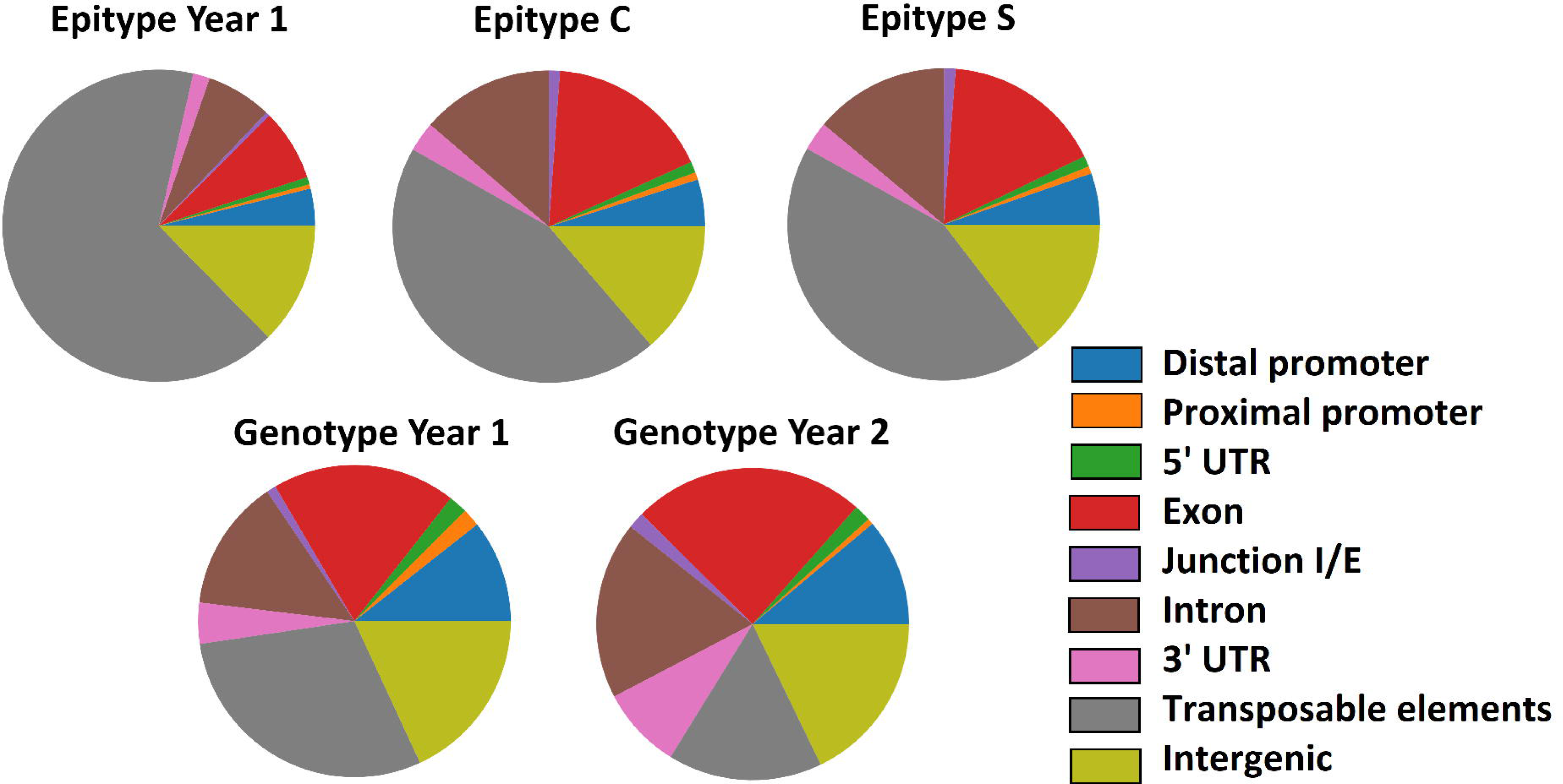
Distribution of DMTEs by transposable element class.

**Fig. S18.**
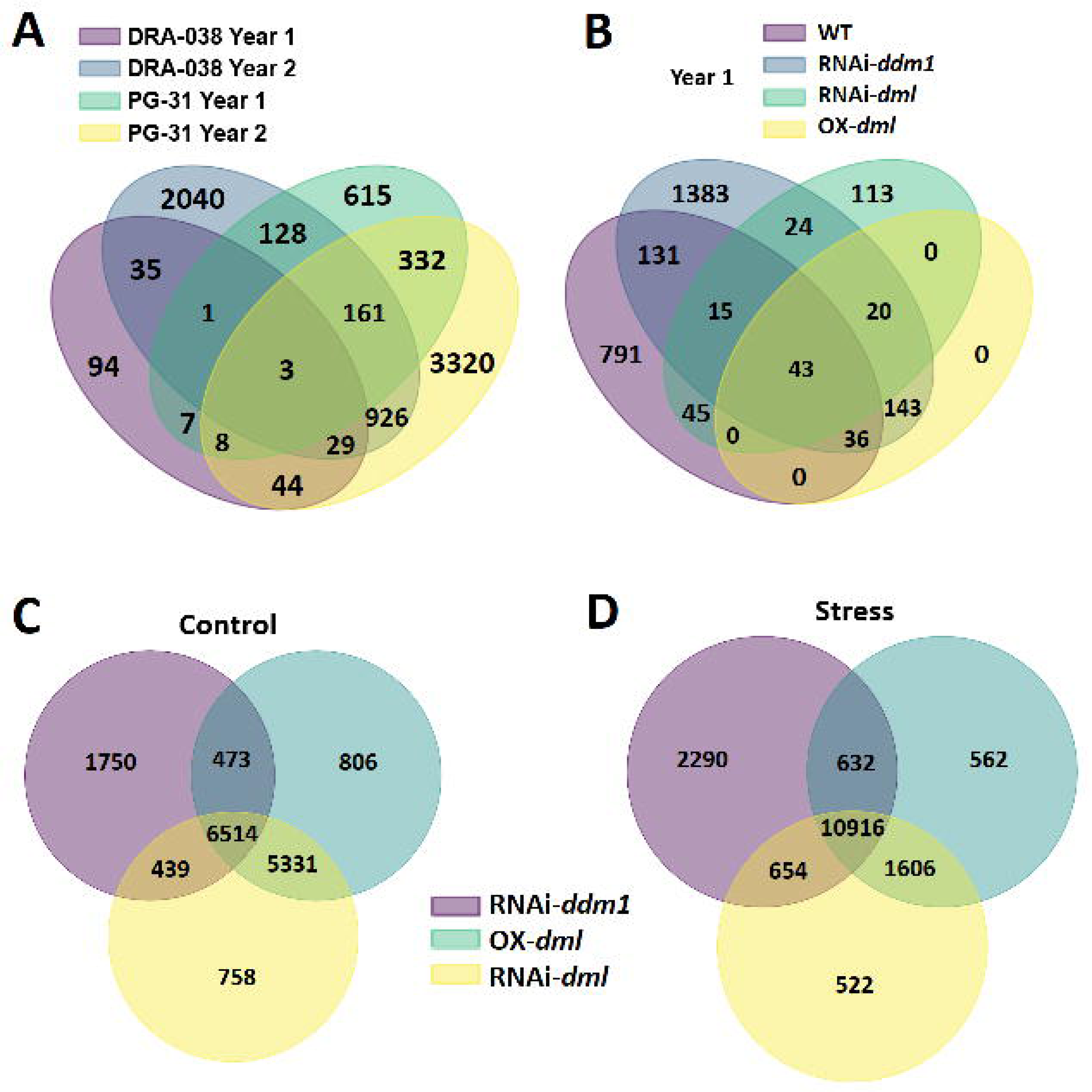
Overlap of differentially methylated genes.

**Fig. S19.**
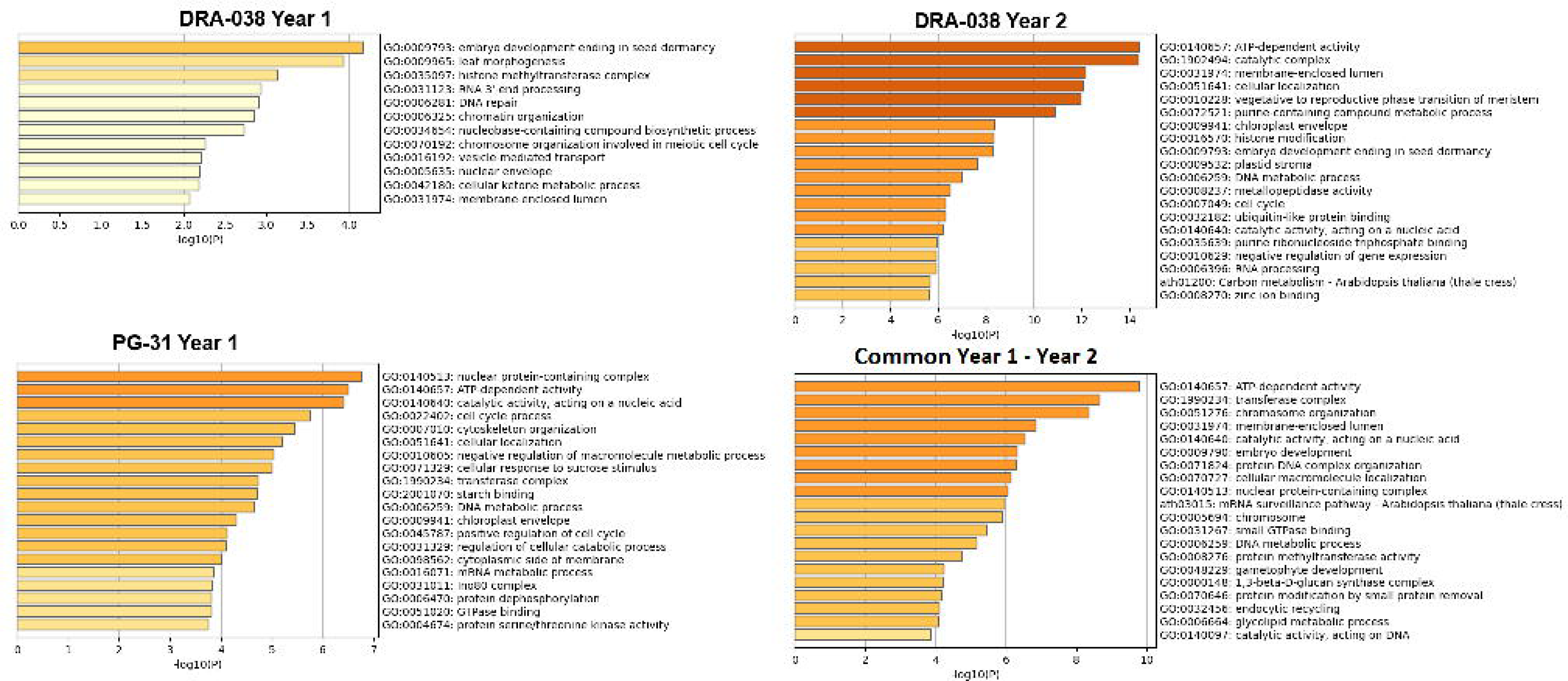
GO enrichment of DMGs in P. nigra. **Fig. S20** TE class distribution among DMTEs. **Fig. S21** GO enrichment of DMGs in epitypes.

**Fig. S20.**
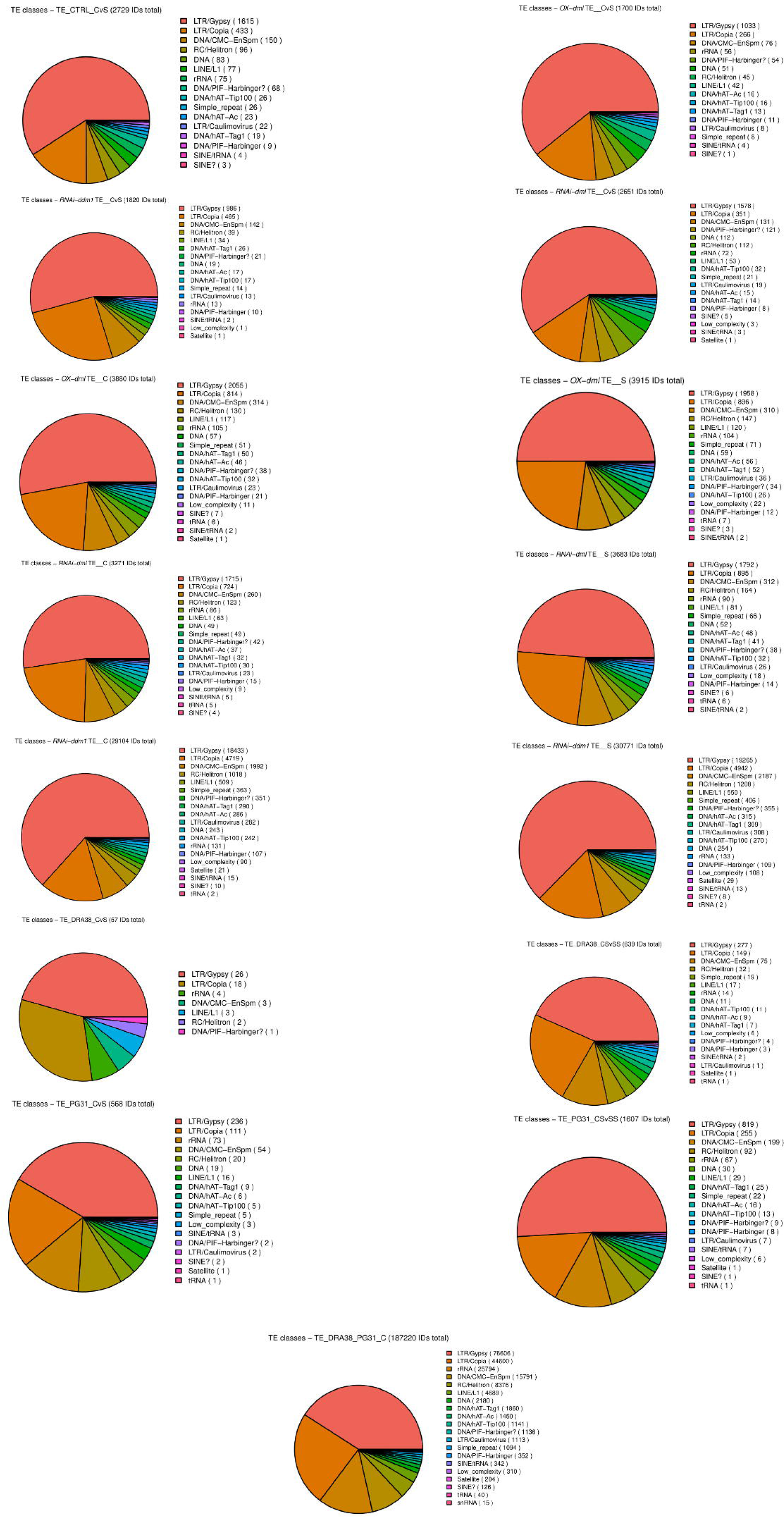
TE class distribution among DMTEs.

**Fig. S21.**
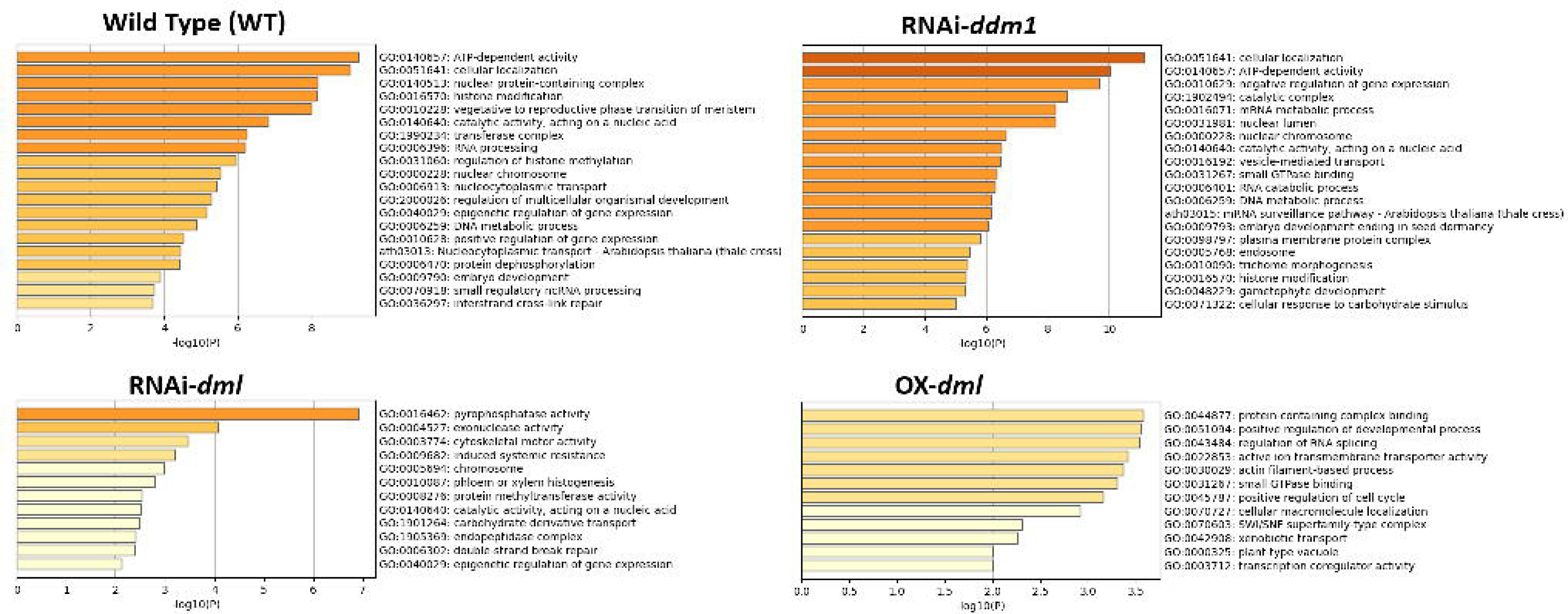
GO enrichment of DMGs in epitypes.

**Fig. S22.**
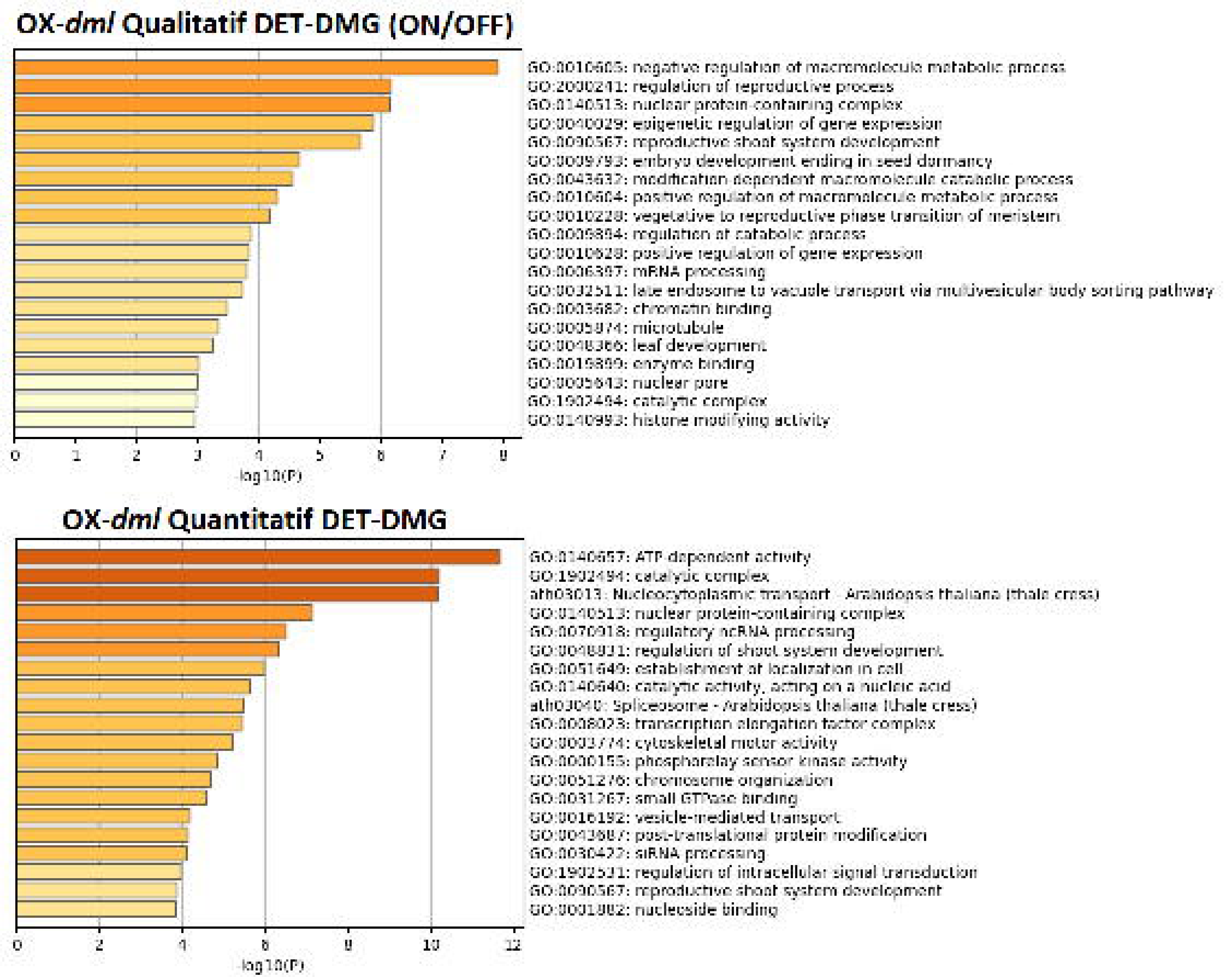
GO enrichment of DET–DMG genes in OX-*dml*.

**Fig. S23.**
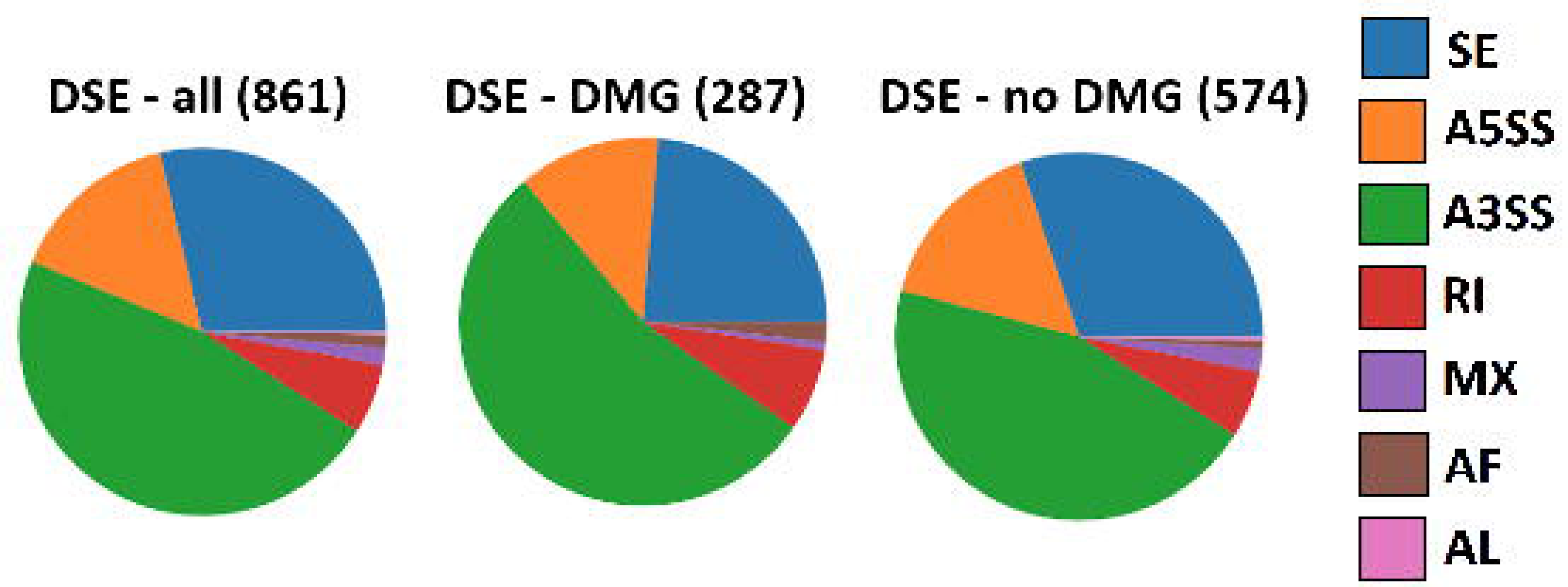
Distribution of differential splicing events.

**Fig. S24.**
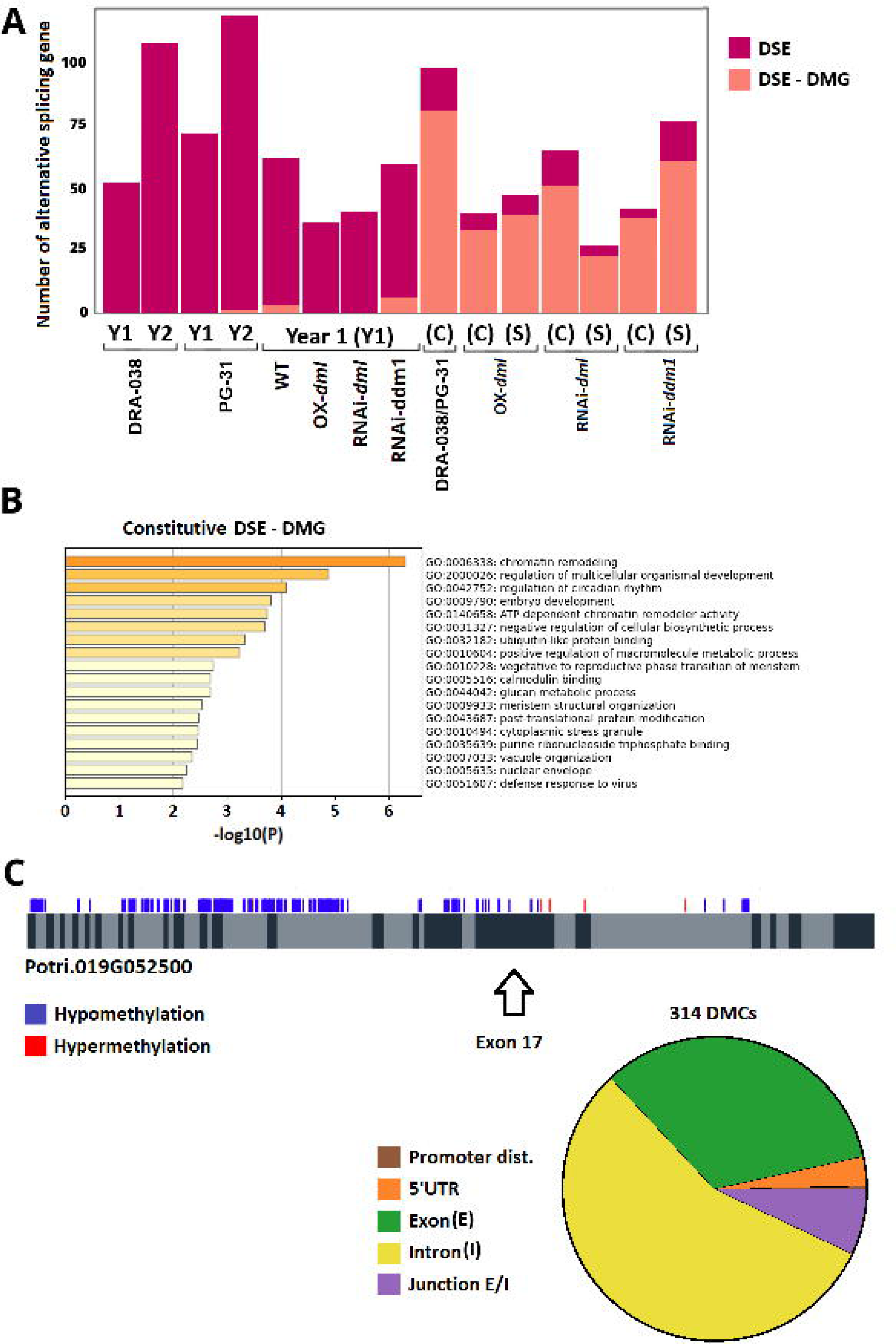
Overlap between DSE and DMG.

**Fig. S25.**
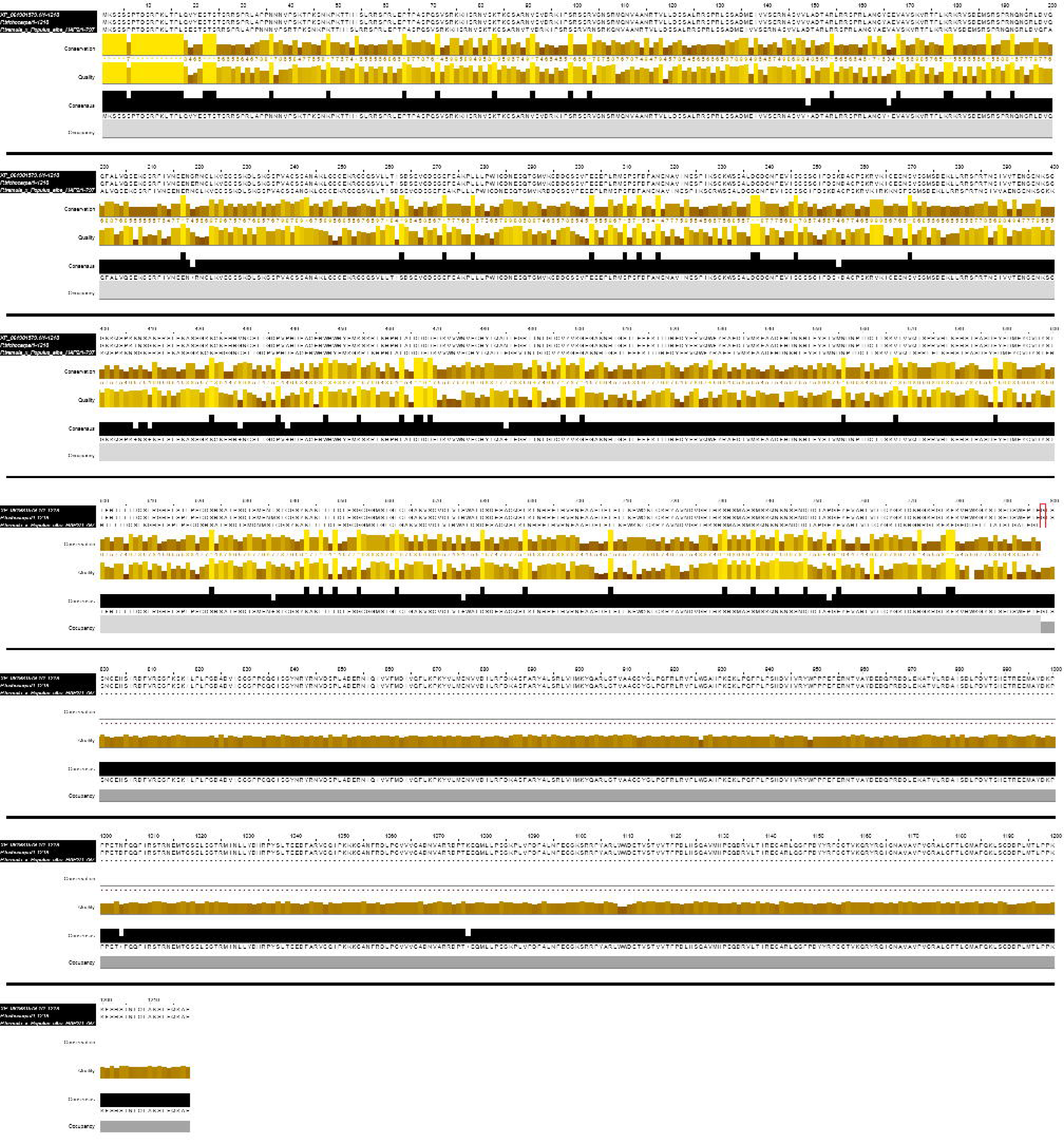
Sequence alignment of PtaDML6 homologs.

**Fig. S26.**
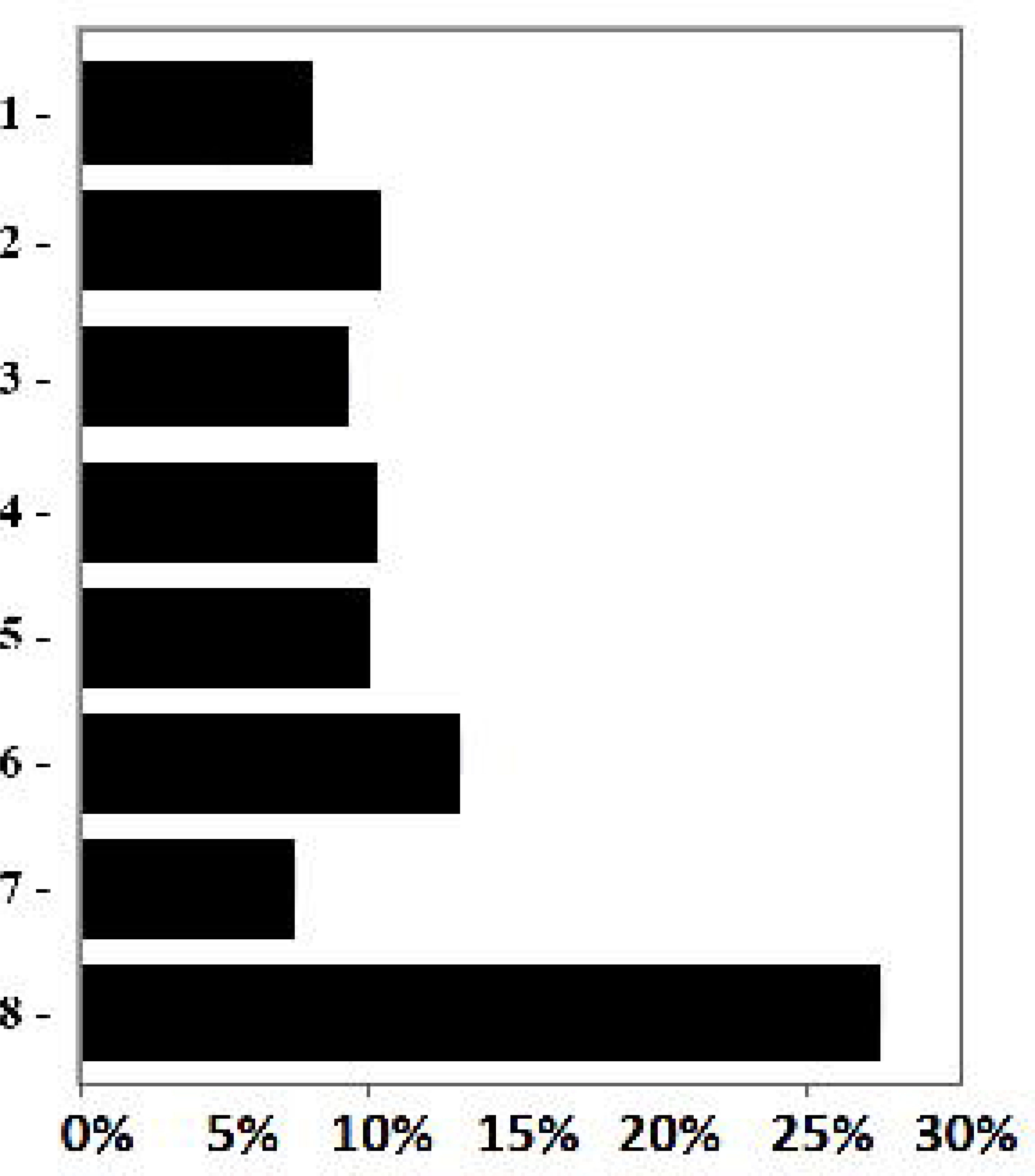
Overlap of differential expression and methylation across published studies.

## Supporting information

**Table S1** Phytohormone profiling in cambium-derived tissues.

**Table S2** Genes of interest identified in P. nigra genotypes.

**Table S3** Shared stress-responsive DETs in P. tremula × P. alba epitypes.

**Table S4** Genes of interest identified in P. tremula × P. alba epitypes.

**Table S5** Memory-related and epitype-specific DETs with DSE classification.

**Table S6** Genes containing stress-induced DMCs.

**Table S7** Differentially expressed transcripts and associated methylation status.

**Supplementary Methods** RNA-seq and WGBS Bioinformatic Workflows.

## References

Akalin, A., Kormaksson, M., Li, S., Garrett-Bakelman, F. E., Figueroa, M. E., Melnick, A., & Mason, C. E. (2012). methylKit : A comprehensive R package for the analysis of genome-wide DNA methylation profiles. Genome Biology, 13(10), R87. 10.1186/gb-2012-13-10-r87

Alves, M. S., Fontes, E. P. B., & Fietto, L. G. (2011). EARLY RESPONSIVE to DEHYDRATION 15, a new transcription factor that integrates stress signaling pathways. Plant Signaling & Behavior, 6(12), 1993–1996. 10.4161/psb.6.12.18268

Amaral, J., Ribeyre, Z., Vigneaud, J., Sow, M. D., Fichot, R., Messier, C., Pinto, G., Nolet, P., & Maury, S. (2020). Advances and Promises of Epigenetics for Forest Trees. Forests, 11(9), Article 9. 10.3390/f11090976

Backhaus, S., Kreyling, J., Grant, K., Beierkuhnlein, C., Walter, J., & Jentsch, A. (2014). Recurrent Mild Drought Events Increase Resistance Toward Extreme Drought Stress. Ecosystems, 17(6), 1068–1081. 10.1007/s10021-014-9781-5

Bäurle, I., & Trindade, I. (2020). Chromatin regulation of somatic abiotic stress memory. Journal of Experimental Botany, 71(17), 5269–5279. 10.1093/jxb/eraa098

Bouyer, L., Vincent-Barbaroux, C., Le Jan, I., Delaunay, A., Marchand, L., Feinard-Duranceau, M., Sallé, A., Chassagnaud, D., Barigah, T. S., Cochard, H., Brignolas, F., & Fichot, R. (2023). Concurrent time course of xylem hydraulic dysfunction and non-structural carbohydrates under contrasting water deficits and nitrogen supplies in poplar. Environmental and Experimental Botany, 206, 105173. 10.1016/j.envexpbot.2022.105173

Brenya, E., Pervin, M., Chen, Z.-H., Tissue, D. T., Johnson, S., Braam, J., & Cazzonelli, C. I. (2022). Mechanical stress acclimation in plants : Linking hormones and somatic memory to thigmomorphogenesis. *Plant*, Cell & Environment, 45(4), 989–1010. 10.1111/pce.14252

Cai, B.-D., Yin, J., Hao, Y.-H., Li, Y.-N., Yuan, B.-F., & Feng, Y.-Q. (2015). Profiling of phytohormones in rice under elevated cadmium concentration levels by magnetic solid-phase extraction coupled with liquid chromatography tandem mass spectrometry. Journal of Chromatography A, 1406, 78–86. 10.1016/j.chroma.2015.06.046

Chateigner, A., Lesage-Descauses, M.-C., Rogier, O., Jorge, V., Leplé, J.-C., Brunaud, V., Roux, C. P.-L., Soubigou-Taconnat, L., Martin-Magniette, M.-L., Sanchez, L., & Segura, V. (2020). Gene expression predictions and networks in natural populations supports the omnigenic theory. BMC Genomics, 21(1), 416. 10.1186/s12864-020-06809-2

Chen, M.-L., Fu, X.-M., Liu, J.-Q., Ye, T.-T., Hou, S.-Y., Huang, Y.-Q., Yuan, B.-F., Wu, Y., & Feng, Y.-Q. (2012). Highly sensitive and quantitative profiling of acidic phytohormones using derivatization approach coupled with nano-LC–ESI-Q-TOF-MS analysis. Journal of Chromatography B, 905, 67–74. 10.1016/j.jchromb.2012.08.005

Claeys, H., & Inzé, D. (2013). The Agony of Choice : How Plants Balance Growth and Survival under Water-Limiting Conditions1. Plant Physiology, 162(4), 1768–1779. 10.1104/pp.113.220921

Cochard, H., Damour, G., Bodet, C., Tharwat, I., Poirier, M., & Améglio, T. (2005). Evaluation of a new centrifuge technique for rapid generation of xylem vulnerability curves. Physiologia Plantarum, 124(4), 410–418. 10.1111/j.1399-3054.2005.00526.x

Cohen, D., Bogeat-Triboulot, M.-B., Tisserant, E., Balzergue, S., Martin-Magniette, M.-L., Lelandais, G., Ningre, N., Renou, J.-P., Tamby, J.-P., Le Thiec, D., & Hummel, I. (2010). Comparative transcriptomics of drought responses in Populus : A meta-analysis of genome-wide expression profiling in mature leaves and root apices across two genotypes. BMC Genomics, 11, 630. 10.1186/1471-2164-11-630

Conde, D., Le Gac, A.-L., Perales, M., Dervinis, C., Kirst, M., Maury, S., González-Melendi, P., & Allona, I. (2017a). Chilling-responsive DEMETER-LIKE DNA demethylase mediates in poplar bud break. *Plant*, Cell & Environment, 40(10), 2236–2249. 10.1111/pce.13019

Conde, D., Moreno-Cortés, A., Dervinis, C., Ramos-Sánchez, J. M., Kirst, M., Perales, M., González-Melendi, P., & Allona, I. (2017b). Overexpression of DEMETER, a DNA demethylase, promotes early apical bud maturation in poplar. *Plant*, Cell & Environment, 40(11), 2806–2819. 10.1111/pce.13056

Crisp, P. A., Ganguly, D., Eichten, S. R., Borevitz, J. O., & Pogson, B. J. (2016). Reconsidering plant memory : Intersections between stress recovery, RNA turnover, and epigenetics. Science Advances, 2(2), e1501340. 10.1126/sciadv.1501340

Ding, J., Mao, L.-J., Wang, S.-T., Yuan, B.-F., & Feng, Y.-Q. (2013). Determination of Endogenous Brassinosteroids in Plant Tissues Using Solid-phase Extraction with Double Layered Cartridge Followed by High-performance Liquid Chromatography–Tandem Mass Spectrometry. Phytochemical Analysis, 24(4), 386–394. 10.1002/pca.2421

Ding, W., Wang, C., Mei, M., Li, X., Zhang, Y., Lin, H., Li, Y., Ma, Z., Han, J., Song, X., Wu, M., Zheng, C., Lin, J., & Zhao, Y. (2024). Phytohormones involved in vascular cambium activity in woods : Current progress and future challenges. Frontiers in Plant Science, 15. 10.3389/fpls.2024.1508242

Dowen, R. H., Pelizzola, M., Schmitz, R. J., Lister, R., Dowen, J. M., Nery, J. R., Dixon, J. E., & Ecker, J. R. (2012). Widespread dynamic DNA methylation in response to biotic stress. Proceedings of the National Academy of Sciences, 109(32), E2183–E2191. 10.1073/pnas.1209329109

Doyle, J. J., & Doyle, J. L. (Éds.). (1987). A rapid DNA isolation procedure for small quantities of fresh leaf tissue. PHYTOCHEMICAL BULLETIN.

Dubin, M. J., Zhang, P., Meng, D., Remigereau, M.-S., Osborne, E. J., Paolo Casale, F., Drewe, P., Kahles, A., Jean, G., Vilhjálmsson, B., Jagoda, J., Irez, S., Voronin, V., Song, Q., Long, Q., Rätsch, G., Stegle, O., Clark, R. M., & Nordborg, M. (2015). DNA methylation in Arabidopsis has a genetic basis and shows evidence of local adaptation. eLife, 4, e05255. 10.7554/eLife.05255

Dugé de Bernonville, T., Daviaud, C., Chaparro, C., Tost, J., & Maury, S. (2022). From Methylome to Integrative Analysis of Tissue Specificity. In V. Courdavault & S. Besseau (Éds.), Catharanthus roseus : Methods and Protocols (p. 223–240). Springer US. 10.1007/978-1-0716-2349-7_16

Duursma, R., & Choat, B. (2017). fitplc—An R package to fit hydraulic vulnerability curves. Journal of Plant Hydraulics, 4, e002–e002. 10.20870/jph.2017.e002

Fàbregas, N., Yoshida, T., & Fernie, A. R. (2020). Role of Raf-like kinases in SnRK2 activation and osmotic stress response in plants. Nature Communications, 11(1), 6184. 10.1038/s41467-020-19977-2

Faivre-Rampant, P., Scalabrin, S., Guérin, V., Paoli, E. de, Aluome, C., Viger, M., Cattonaro, F., Payne, A., Paulstephenraj, P. S., Paslier, M.-C. L., Berard, A. A., Allwright, M., Villar, M. M., Taylor, G., Bastien, C., & Morgante, M. (2016). New resources for genetic studies in Populus nigra : Genome-wide SNP discovery and development of a 12k Infinium array. Molecular Ecology Resources, 16(4), 1023. 10.1111/1755-0998.12513

Farkas, D., & Dobránszki, J. (2024). Vegetal memory through the lens of transcriptomic changes—Recent progress and future practical prospects for exploiting plant transcriptional memory. Plant Signaling & Behavior, 19(1), 2383515. 10.1080/15592324.2024.2383515

Fichot, R., Brignolas, F., Cochard, H., & Ceulemans, R. (2015). Vulnerability to drought-induced cavitation in poplars : Synthesis and future opportunities. Plant, Cell & Environment, 38(7), 1233–1251. 10.1111/pce.12491

Fukuda, H. (2004). Signals that control plant vascular cell differentiation. Nature Reviews Molecular Cell Biology, 5(5), 379–391. 10.1038/nrm1364

Gallusci, P., Agius, D. R., Moschou, P. N., Dobránszki, J., Kaiserli, E., & Martinelli, F. (2023). Deep inside the epigenetic memories of stressed plants. Trends in Plant Science, 28(2), 142–153. 10.1016/j.tplants.2022.09.004

Galviz, Y. C. F., Ribeiro, R. V., & Souza, G. M. (2020). Yes, plants do have memory. Theoretical and Experimental Plant Physiology, 32(3), 195–202. 10.1007/s40626-020-00181-y

Gebreselassie, M. N., Ader, K., Boizot, N., Millier, F., Charpentier, J.-P., Alves, A., Simões, R., Rodrigues, J. C., Bodineau, G., Fabbrini, F., Sabatti, M., Bastien, C., & Segura, V. (2017). Near-infrared spectroscopy enables the genetic analysis of chemical properties in a large set of wood samples from *Populus nigra* (L.) natural populations. Industrial Crops and Products, 107, 159–171. 10.1016/j.indcrop.2017.05.013

Gessler, A., Bottero, A., Marshall, J., & Arend, M. (2020). The way back : Recovery of trees from drought and its implication for acclimation. New Phytologist, 228(6), 1704–1709. 10.1111/nph.16703

Groover, A. (2025). Roles for epigenetics in wood formation and stress response intrees–from basic biology to forest management. Frontiers in Epigenetics and Epigenomics, 3. 10.3389/freae.2025.1476499

Guet, J., Fichot, R., Lédée, C., Laurans, F., Cochard, H., Delzon, S., Bastien, C., & Brignolas, F. (2015). Stem xylem resistance to cavitation is related to xylem structure but not to growth and water-use efficiency at the within-population level in *Populus nigra* L. Journal of Experimental Botany, 66(15), 4643–4652. 10.1093/jxb/erv232

Hammond, W. M., Williams, A. P., Abatzoglou, J. T., Adams, H. D., Klein, T., López, R., Sáenz-Romero, C., Hartmann, H., Breshears, D. D., & Allen, C. D. (2022). Global field observations of tree die-off reveal hotter-drought fingerprint for Earth’s forests. Nature Communications, 13(1), 1761. 10.1038/s41467-022-29289-2

Harris, C. J., Amtmann, A., & Ton, J. (2023). Epigenetic processes in plant stress priming : Open questions and new approaches. Current Opinion in Plant Biology, 75, 102432. 10.1016/j.pbi.2023.102432

Hilker, M., Schwachtje, J., Baier, M., Balazadeh, S., Bäurle, I., Geiselhardt, S., Hincha, D. K., Kunze, R., Mueller-Roeber, B., Rillig, M. C., Rolff, J., Romeis, T., Schmülling, T., Steppuhn, A., van Dongen, J., Whitcomb, S. J., Wurst, S., Zuther, E., & Kopka, J. (2016). Priming and memory of stress responses in organisms lacking a nervous system. Biological Reviews of the Cambridge Philosophical Society, 91(4), 1118–1133. 10.1111/brv.12215

Holeski, L. M., Jander, G., & Agrawal, A. A. (2012). Transgenerational defense induction and epigenetic inheritance in plants. Trends in Ecology & Evolution, 27(11), 618–626. 10.1016/j.tree.2012.07.011

Inácio, V., Santos, R., Prazeres, R., Graça, J., Miguel, C. M., & Morais-Cecílio, L. (2022). Epigenetics at the crossroads of secondary growth regulation. Frontiers in Plant Science, 13. 10.3389/fpls.2022.970342

Jaspers, P., Overmyer, K., Wrzaczek, M., Vainonen, J. P., Blomster, T., Salojärvi, J., Reddy, R. A., & Kangasjärvi, J. (2010). The RST and PARP-like domain containing SRO protein family : Analysis of protein structure, function and conservation in land plants. BMC Genomics, 11(1), 170. 10.1186/1471-2164-11-170

Kakoulidou, I., Avramidou, E. V., Baránek, M., Brunel-Muguet, S., Farrona, S., Johannes, F., Kaiserli, E., Lieberman-Lazarovich, M., Martinelli, F., Mladenov, V., Testillano, P. S., Vassileva, V., & Maury, S. (2021). Epigenetics for Crop Improvement in Times of Global Change. Biology, 10(8), Article 8. 10.3390/biology10080766

Kim, M.-H., Cho, J.-S., Jeon, H.-W., Sangsawang, K., Shim, D., Choi, Y.-I., Park, E.-J., Lee, H., & Ko, J.-H. (2019). Wood Transcriptome Profiling Identifies Critical Pathway Genes of Secondary Wall Biosynthesis and Novel Regulators for Vascular Cambium Development in Populus. Genes, 10(9), 690. 10.3390/genes10090690

Krueger, F., James, F., Ewels, P., Afyounian, E., Weinstein, M., Schuster-Boeckler, B., Hulselmans, G., & sclamons. (2023). FelixKrueger/TrimGalore : V0.6.10 - add default decompression path (Version 0.6.10) [Logiciel]. Zenodo. 10.5281/zenodo.7598955

Kuromori, T., Seo, M., & Shinozaki, K. (2018). ABA Transport and Plant Water Stress Responses. Trends in Plant Science, 23(6), 513–522. 10.1016/j.tplants.2018.04.001

Lafon-Placette, C., Faivre-Rampant, P., Delaunay, A., Street, N., Brignolas, F., & Maury, S. (2013). Methylome of DNase I sensitive chromatin in Populus trichocarpa shoot apical meristematic cells : A simplified approach revealing characteristics of gene-body DNA methylation in open chromatin state. New Phytologist, 197(2), 416–430. 10.1111/nph.12026

Lafon-Placette, C., Le Gac, A.-L., Chauveau, D., Segura, V., Delaunay, A., Lesage-Descauses, M.-C., Hummel, I., Cohen, D., Jesson, B., Le Thiec, D., Bogeat-Triboulot, M.-B., Brignolas, F., & Maury, S. (2018). Changes in the epigenome and transcriptome of the poplar shoot apical meristem in response to water availability affect preferentially hormone pathways. Journal of Experimental Botany, 69(3), 537–551. 10.1093/jxb/erx409

Lämke, J., & Bäurle, I. (2017). Epigenetic and chromatin-based mechanisms in environmental stress adaptation and stress memory in plants. Genome Biology, 18(1), 124. 10.1186/s13059-017-1263-6

Langmead, B., & Salzberg, S. L. (2012). Fast gapped-read alignment with Bowtie 2. Nature Methods, 9(4). 10.1038/nmeth.1923

Le Gac, A. L., Lafon-Placette, C., Delaunay, A., & Maury, S. (2019). Developmental, genetic and environmental variations of global DNA methylation in the first leaves emerging from the shoot apical meristem in poplar trees. Plant Signaling & Behavior, 14(6), 1596717. 10.1080/15592324.2019.1596717

Le Gac, A.-L., Lafon-Placette, C., Chauveau, D., Segura, V., Delaunay, A., Fichot, R., Marron, N., Le Jan, I., Berthelot, A., Bodineau, G., Bastien, J.-C., Brignolas, F., & Maury, S. (2018). Winter-dormant shoot apical meristem in poplar trees shows environmental epigenetic memory. Journal of Experimental Botany, 69(20), 4821–4837. 10.1093/jxb/ery271

Leple, J. C., Brasileiro, A. C. M., Michel, M. F., Delmotte, F., & Jouanin, L. (1992). Transgenic poplars : Expression of chimeric genes using four different constructs. Plant Cell Reports, 11(3), 137–141. 10.1007/BF00232166

Liang, D., Zhang, Z., Wu, H., Huang, C., Shuai, P., Ye, C.-Y., Tang, S., Wang, Y., Yang, L., Wang, J., Yin, W., & Xia, X. (2014). Single-base-resolution methylomes of populus trichocarpa reveal the association between DNA methylation and drought stress. BMC Genetics, 15(Suppl 1), S9. 10.1186/1471-2156-15-S1-S9

Lin, Z., Li, Y., Zhang, Z., Liu, X., Hsu, C.-C., Du, Y., Sang, T., Zhu, C., Wang, Y., Satheesh, V., Pratibha, P., Zhao, Y., Song, C.-P., Tao, W. A., Zhu, J.-K., & Wang, P. (2020). A RAF-SnRK2 kinase cascade mediates early osmotic stress signaling in higher plants. Nature Communications, 11(1), 613. 10.1038/s41467-020-14477-9

Liu, X., Wei, X., Sheng, Z., Jiao, G., Tang, S., Luo, J., & Hu, P. (2016). Polycomb Protein OsFIE2 Affects Plant Height and Grain Yield in Rice. PLOS ONE, 11(10), e0164748. 10.1371/journal.pone.0164748

Lloyd, J. P. B., & Lister, R. (2022). Epigenome plasticity in plants. Nature Reviews. Genetics, 23(1), 55–68. 10.1038/s41576-021-00407-y

López Sánchez, A., Stassen, J. H. M., Furci, L., Smith, L. M., & Ton, J. (2016). The role of DNA (de)methylation in immune responsiveness of Arabidopsis. The Plant Journal: For Cell and Molecular Biology, 88(3), 361–374. 10.1111/tpj.13252

Love, M. I., Huber, W., & Anders, S. (2014). Moderated estimation of fold change and dispersion for RNA-seq data with DESeq2. Genome Biology, 15(12), 550. 10.1186/s13059-014-0550-8

Luo, Y., Takagi, J., Claus, L. A. N., Zhang, C., Yasuda, S., Hasegawa, Y., Yamaguchi, J., Shan, L., Russinova, E., & Sato, T. (2022). Deubiquitinating enzymes UBP12 and UBP13 stabilize the brassinosteroid receptor BRI1. EMBO reports, 23(4), e53354. 10.15252/embr.202153354

Mader, M., Le Paslier, M.-C., Bounon, R., Bérard, A., Faivre-Rampant, P., Fladung, M., Leplé, J.-C., & Kersten, B. (2016). Whole-genome draft assembly of *Populus tremula* x *P. alba* clone INRA 717-1B4. Silvae Genetica, 65(2), 74–79. 10.1515/sg-2016-0019

Mähönen, A. P., Bishopp, A., Higuchi, M., Nieminen, K. M., Kinoshita, K., Törmäkangas, K., Ikeda, Y., Oka, A., Kakimoto, T., & Helariutta, Y. (2006). Cytokinin Signaling and Its Inhibitor AHP6 Regulate Cell Fate During Vascular Development. Science, 311(5757), 94–98. 10.1126/science.1118875

Mauch-Mani, B., Baccelli, I., Luna, E., & Flors, V. (2017). Defense Priming : An Adaptive Part of Induced Resistance. Annual Review of Plant Biology, 68(Volume 68, 2017), 485–512. 10.1146/annurev-arplant-042916-041132

Maury, S., Sow, M. D., Le Gac, A.-L., Genitoni, J., Lafon-Placette, C., & Mozgova, I. (2019). Phytohormone and Chromatin Crosstalk : The Missing Link For Developmental Plasticity? Frontiers in Plant Science, 10, 395. 10.3389/fpls.2019.00395

McDowell, N. G., Sapes, G., Pivovaroff, A., Adams, H. D., Allen, C. D., Anderegg, W. R. L., Arend, M., Breshears, D. D., Brodribb, T., Choat, B., Cochard, H., De Cáceres, M., De Kauwe, M. G., Grossiord, C., Hammond, W. M., Hartmann, H., Hoch, G., Kahmen, A., Klein, T., … Xu, C. (2022). Mechanisms of woody-plant mortality under rising drought, CO2 and vapour pressure deficit. Nature Reviews Earth & Environment, 3(5), 294–308. 10.1038/s43017-022-00272-1

Mhiri, C., Borges, F., & Grandbastien, M.-A. (2022). Specificities and Dynamics of Transposable Elements in Land Plants. Biology, 11(4), 488. 10.3390/biology11040488

Muyle, A. M., Seymour, D. K., Lv, Y., Huettel, B., & Gaut, B. S. (2022). Gene Body Methylation in Plants : Mechanisms, Functions, and Important Implications for Understanding Evolutionary Processes. Genome Biology and Evolution, 14(4), evac038. 10.1093/gbe/evac038

Nicotra, A. B., Atkin, O. K., Bonser, S. P., Davidson, A. M., Finnegan, E. J., Mathesius, U., Poot, P., Purugganan, M. D., Richards, C. L., Valladares, F., & van Kleunen, M. (2010). Plant phenotypic plasticity in a changing climate. Trends in Plant Science, 15(12), 684–692. 10.1016/j.tplants.2010.09.008

Niederhuth, C. E., Bewick, A. J., Ji, L., Alabady, M. S., Kim, K. D., Li, Q., Rohr, N. A., Rambani, A., Burke, J. M., Udall, J. A., Egesi, C., Schmutz, J., Grimwood, J., Jackson, S. A., Springer, N. M., & Schmitz, R. J. (2016). Widespread natural variation of DNA methylation within angiosperms. Genome Biology, 17(1), 194. 10.1186/s13059-016-1059-0

Nieminen, K., Blomster, T., Helariutta, Y., & Mähönen, A. P. (2015). Vascular Cambium Development. The Arabidopsis Book / American Society of Plant Biologists, 13, e0177. 10.1199/tab.0177

Nieminen, K., Immanen, J., Laxell, M., Kauppinen, L., Tarkowski, P., Dolezal, K., Tähtiharju, S., Elo, A., Decourteix, M., Ljung, K., Bhalerao, R., Keinonen, K., Albert, V. A., & Helariutta, Y. (2008). Cytokinin signaling regulates cambial development in poplar. Proceedings of the National Academy of Sciences, 105(50), 20032–20037. 10.1073/pnas.0805617106

Ojolo, S. P., Cao, S., Priyadarshani, S. V. G. N., Li, W., Yan, M., Aslam, M., Zhao, H., & Qin, Y. (2018). Regulation of Plant Growth and Development : A Review From a Chromatin Remodeling Perspective. Frontiers in Plant Science, 9, 1232. 10.3389/fpls.2018.01232

Park, S.-H., Jeong, J. S., Zhou, Y., Binte Mustafa, N. F., & Chua, N.-H. (2022). Deubiquitination of BES1 by UBP12/UBP13 promotes brassinosteroid signaling and plant growth. Plant Communications, 3(5), 100348. 10.1016/j.xplc.2022.100348

Patro, R., Duggal, G., Love, M. I., Irizarry, R. A., & Kingsford, C. (2017). Salmon provides fast and bias-aware quantification of transcript expression. Nature Methods, 14(4), 417–419. 10.1038/nmeth.4197

Peirats-Llobet, M., Han, S.-K., Gonzalez-Guzman, M., Jeong, C. W., Rodriguez, L., Belda-Palazon, B., Wagner, D., & Rodriguez, P. L. (2016). A Direct Link between Abscisic Acid Sensing and the Chromatin-Remodeling ATPase BRAHMA via Core ABA Signaling Pathway Components. Molecular Plant, 9(1), 136–147. 10.1016/j.molp.2015.10.003

Peña-Ponton, C., Diez-Rodriguez, B., Perez-Bello, P., Becker, C., McIntyre, L. M., van der Putten, W. H., De Paoli, E., Heer, K., Opgenoorth, L., & Verhoeven, K. J. F. (2024). High-resolution methylome analysis uncovers stress-responsive genomic hotspots and drought-sensitive transposable element superfamilies in the clonal Lombardy poplar. Journal of Experimental Botany, 75(18), 5839–5856. 10.1093/jxb/erae262

Planas-Riverola, A., Gupta, A., Betegón-Putze, I., Bosch, N., Ibañes, M., & Caño-Delgado, A. I. (2019). Brassinosteroid signaling in plant development and adaptation to stress. *Development (Cambridge*, England), 146(5), dev151894. 10.1242/dev.151894

Plomion, C., Aury, J.-M., Amselem, J., Leroy, T., Murat, F., Duplessis, S., Faye, S., Francillonne, N., Labadie, K., Le Provost, G., Lesur, I., Bartholomé, J., Faivre-Rampant, P., Kohler, A., Leplé, J.-C., Chantret, N., Chen, J., Diévart, A., Alaeitabar, T., … Salse, J. (2018). Oak genome reveals facets of long lifespan. Nature Plants, 4(7), 440–452. 10.1038/s41477-018-0172-3

Ramakrishnan, M., Satish, L., Kalendar, R., Narayanan, M., Kandasamy, S., Sharma, A., Emamverdian, A., Wei, Q., & Zhou, M. (2022). Ramakrishnan et al. The Dynamism of Transposon Methylation for Plant Development and Stress Adaptation. Int. J. Mol. Sci. 2021, 22, 11387. International Journal of Molecular Sciences, 23(22), 14107. 10.3390/ijms232214107

Richards, E. J. (2006). Inherited epigenetic variation—Revisiting soft inheritance. Nature Reviews Genetics, 7(5), 395–401. 10.1038/nrg1834

Rogier, O., Chateigner, A., Lesage-Descauses, M.-C., Mandin, C., Brunaud, V., Caius, J., Soubigou-Taconnat, L., Almeida-Falcon, J., Bastien, C., Benoit, V., Bodineau, G., Boizot, N., Buret, C., Charpentier, J.-P., Déjardin, A., Delaunay, A., Fichot, R., Laine Prade, V., Laurans, F., … Segura, V. (2023). RNAseq based variant dataset in a black poplar association panel. BMC Research Notes, 16(1), 248. 10.1186/s13104-023-06521-w

Ruzicka, K., Ursache, R., & Hejátko, J. (2015, mars 23). Xylem development – from the cradle to the grave. https://nph.onlinelibrary.wiley.com/doi/10.1111/nph.13383

Shen, S., Park, J. W., Lu, Z., Lin, L., Henry, M. D., Wu, Y. N., Zhou, Q., & Xing, Y. (2014a). rMATS : Robust and flexible detection of differential alternative splicing from replicate RNA-Seq data. Proceedings of the National Academy of Sciences of the United States of America, 111(51), E5593–5601. 10.1073/pnas.1419161111

Shen, X., De Jonge, J., Forsberg, S. K. G., Pettersson, M. E., Sheng, Z., Hennig, L., & Carlborg, Ö. (2014b). Natural CMT2 variation is associated with genome-wide methylation changes and temperature seasonality. PLoS Genetics, 10(12), e1004842. 10.1371/journal.pgen.1004842

Smetana, O., Mäkilä, R., Lyu, M., Amiryousefi, A., Sánchez Rodríguez, F., Wu, M.-F., Solé-Gil, A., Leal Gavarrón, M., Siligato, R., Miyashima, S., Roszak, P., Blomster, T., Reed, J. W., Broholm, S., & Mähönen, A. P. (2019). High levels of auxin signalling define the stem-cell organizer of the vascular cambium. Nature, 565(7740), 485–489. 10.1038/s41586-018-0837-0

Sow, M. D., Le Gac, A.-L., Fichot, R., Lanciano, S., Delaunay, A., Le Jan, I., Lesage-Descauses, M.-C., Citerne, S., Caius, J., Brunaud, V., Soubigou-Taconnat, L., Cochard, H., Segura, V., Chaparro, C., Grunau, C., Daviaud, C., Tost, J., Brignolas, F., Strauss, S. H., … Maury, S. (2021). RNAi suppression of DNA methylation affects the drought stress response and genome integrity in transgenic poplar. The New Phytologist, 232(1), 80–97. 10.1111/nph.17555

Sow, M. D., Rogier, O., Lesur, I., Daviaud, C., Mardoc, E., Sanou, E., Duvaux, L., Civan, P., Delaunay, A., Descauses, M.-C. L.-, Benoit, V., Le-Jan, I., Buret, C., Besse, C., Durufle, H., Fichot, R., Le-Provost, G., Guichoux, E., Boury, C., … Salse, J. (2023). Epigenetic Variation in Tree Evolution : A case study in black poplar (Populus nigra) (p. 2023.07.16.549253). bioRxiv. 10.1101/2023.07.16.549253

Steinbach, D., Alaux, M., Amselem, J., Choisne, N., Durand, S., Flores, R., Keliet, A.-O., Kimmel, E., Lapalu, N., Luyten, I., Michotey, C., Mohellibi, N., Pommier, C., Reboux, S., Valdenaire, D., Verdelet, D., & Quesneville, H. (2013). GnpIS : An information system to integrate genetic and genomic data from plants and fungi. Database: The Journal of Biological Databases and Curation, 2013, bat058. 10.1093/database/bat058

Stroud, H., Do, T., Du, J., Zhong, X., Feng, S., Johnson, L., Patel, D. J., & Jacobsen, S. E. (2014). Non-CG methylation patterns shape the epigenetic landscape in Arabidopsis. Nature Structural & Molecular Biology, 21(1), 64–72. 10.1038/nsmb.2735

Talarico, E., Zambelli, A., Araniti, F., Greco, E., Chiappetta, A., & Bruno, L. (2024). Unravelling the Epigenetic Code : DNA Methylation in Plants and Its Role in Stress Response. Epigenomes, 8(3), 30. 10.3390/epigenomes8030030

Teotia, S., & Lamb, R. S. (2011). RCD1 and SRO1 are necessary to maintain meristematic fate in Arabidopsis thaliana. Journal of Experimental Botany, 62(3), 1271–1284. 10.1093/jxb/erq363

Teotia, S., Muthuswamy, S., & Lamb, R. (2009, octobre 21). Full article : Radical-induced cell death1 and similar to RCD one1 and the stress-induced morphogenetic response. https://www.tandfonline.com/doi/full/10.4161/psb.5.2.10400

Tong, H., Xiao, Y., Liu, D., Gao, S., Liu, L., Yin, Y., Jin, Y., Qian, Q., & Chu, C. (2014). Brassinosteroid Regulates Cell Elongation by Modulating Gibberellin Metabolism in Rice. The Plant Cell, 26(11), 4376–4393. 10.1105/tpc.114.132092

Trincado, J. L., Entizne, J. C., Hysenaj, G., Singh, B., Skalic, M., Elliott, D. J., & Eyras, E. (2018). SUPPA2 : Fast, accurate, and uncertainty-aware differential splicing analysis across multiple conditions. Genome Biology, 19(1), 40. 10.1186/s13059-018-1417-1

Trontin, J.-F., Raschke, J., & Rupps, A. (2021). Tree ‘memory’ : New insights on temperature-induced priming effects during early embryogenesis. Tree Physiology, 41(6), 906–911. 10.1093/treephys/tpaa150

Trontin, J.-F., Sow, M. D., Delaunay, A., Modesto, I., Teyssier, C., Reymond, I., Canlet, F., Boizot, N., Le Metté, C., Gibert, A., Chaparro, C., Daviaud, C., Tost, J., Miguel, C., Lelu-Walter, M.-A., & Maury, S. (2024). Epigenetic memory of temperature sensed during somatic embryo maturation in 2-year-old maritime pine trees. *Plant Physiology*, kiae600. 10.1093/plphys/kiae600

Vanden Broeck, A., Meese, T., Verschelde, P., Cox, K., Heinze, B., Deforce, D., De Meester, E., & Van Nieuwerburgh, F. (2024). Genome-wide methylome stability and parental effects in the worldwide distributed Lombardy poplar. BMC Biology, 22(1), 30. 10.1186/s12915-024-01816-1

Vigneaud, J., Kohler, A., Sow, M. D., Delaunay, A., Fauchery, L., Guinet, F., Daviaud, C., Barry, K. W., Keymanesh, K., Johnson, J., Singan, V., Grigoriev, I., Fichot, R., Conde, D., Perales, M., Tost, J., Martin, F. M., Allona, I., Strauss, S. H., … Maury, S. (2023). DNA hypomethylation of the host tree impairs interaction with mutualistic ectomycorrhizal fungus. New Phytologist, 238(6), 2561–2577. 10.1111/nph.18734

Vongs, A., Kakutani, T., Martienssen, R. A., & Richards, E. J. (1993). Arabidopsis thaliana DNA methylation mutants. Science (New York, N.Y.), 260(5116), 1926–1928. 10.1126/science.8316832

Wang, X., Hu, L., Wang, X., Li, N., Xu, C., Gong, L., & Liu, B. (2016). DNA Methylation Affects Gene Alternative Splicing in Plants : An Example from Rice. Molecular Plant, 9(2), 305–307. 10.1016/j.molp.2015.09.016

Westerband, A. C., Funk, J. L., & Barton, K. E. (2021). Intraspecific trait variation in plants : A renewed focus on its role in ecological processes. Annals of Botany, 127(4), 397–410. 10.1093/aob/mcab011

Wilkins, O., Waldron, L., Nahal, H., Provart, N. J., & Campbell, M. M. (2009). Genotype and time of day shape the Populus drought response. The Plant Journal, 60(4), 703–715. 10.1111/j.1365-313X.2009.03993.x

Wybouw, B., Zhang, X., & Mähönen, A. P. (2024). Vascular cambium stem cells : Past, present and future. New Phytologist, 243(3), 851–865. 10.1111/nph.19897

Xie, G., Du, X., Hu, H., & Du, J. (2025). Molecular Mechanisms Underlying the Establishment, Maintenance, and Removal of DNA Methylation in Plants. 10.1146/annurev-arplant-083123-054357

Xiong, M., Feng, G.-N., Gao, Q., Zhang, C.-Q., Li, Q.-F., Liu, Q.-Q., Xiong, M., Feng, G.-N., Gao, Q., Zhang, C.-Q., Li, Q.-F., & Liu, Q.-Q. (2022). Brassinosteroid regulation in rice seed biology. Seed Biology, 1(1), 1–9. 10.48130/SeedBio-2022-0002

Xu, G., & Law, J. A. (2024). Loops, crosstalk, and compartmentalization : It takes many layers to regulate DNA methylation. Current Opinion in Genetics & Development, 84, 102147. 10.1016/j.gde.2023.102147

Yamamuro, C., Zhu, J.-K., & Yang, Z. (2016). Epigenetic Modifications and Plant Hormone Action. Molecular Plant, 9(1), 57–70. 10.1016/j.molp.2015.10.008

Yang, J., & Johannes, F. (2025). DNA methylation dynamics in the shoot apical meristem. Journal of Experimental Botany, 76(9), 2478–2486. 10.1093/jxb/eraf024

Zhang, H., Lang, Z., & Zhu, J.-K. (2018). Dynamics and function of DNA methylation in plants. Nature Reviews. Molecular Cell Biology, 19(8), 489–506. 10.1038/s41580-018-0016-z

Zhang, S., Hu, N., & Yu, F. (2024). Insights into a functional model of key deubiquitinases UBP12/13 in plants. New Phytologist, 242(2), 424–430. 10.1111/nph.19639

Zhang, X., Yazaki, J., Sundaresan, A., Cokus, S., Chan, S. W.-L., Chen, H., Henderson, I. R., Shinn, P., Pellegrini, M., Jacobsen, S. E., & Ecker, J. R. (2006). Genome-wide high-resolution mapping and functional analysis of DNA methylation in arabidopsis. Cell, 126(6), 1189–1201. 10.1016/j.cell.2006.08.003

Zhang, Y.-Z., Yuan, J., Zhang, L., Chen, C., Wang, Y., Zhang, G., Peng, L., Xie, S.-S., Jiang, J., Zhu, J.-K., Du, J., & Duan, C.-G. (2020). Coupling of H3K27me3 recognition with transcriptional repression through the BAH-PHD-CPL2 complex in Arabidopsis. Nature Communications, 11, 6212. 10.1038/s41467-020-20089-0

Zhao, B., Liu, Q., Wang, B., & Yuan, F. (2021). Roles of Phytohormones and Their Signaling Pathways in Leaf Development and Stress Responses. Journal of Agricultural and Food Chemistry, 69(12), 3566–3584. 10.1021/acs.jafc.0c07908

Zhou, W., Wang, M., Wang, L., Liu, Y., Tian, Z., Xie, L., & Wang, Y. (2025). Epigenetics in Plant Response to Climate Change. Biology, 14(6), 631. 10.3390/biology14060631

Zhou, Y., Zhou, B., Pache, L., Chang, M., Khodabakhshi, A. H., Tanaseichuk, O., Benner, C., & Chanda, S. K. (2019). Metascape provides a biologist-oriented resource for the analysis of systems-level datasets. Nature Communications, 10(1), 1523. 10.1038/s41467-019-09234-6

Zhu, R., Shevchenko, O., Ma, C., Maury, S., Freitag, M., & Strauss, S. H. (2013). Poplars with a PtDDM1-RNAi transgene have reduced DNA methylation and show aberrant post-dormancy morphology. Planta, 237(6), 1483–1493. 10.1007/s00425-013-1858-4

Zhu, R., Xue, Y., & Qian, W. (2025). Molecular mechanisms and biological functions of active DNA demethylation in plants. Epigenetics & Chromatin, 18(1), 41. 10.1186/s13072-025-00605-6

