## Supplementary material for "Drought-Induced Epigenetic Memory in the Cambium of Poplar Trees persists and primes future stress responses": Table S1

Sup_Table_1 : Summary table of the main trends observed on the measurement of the 15 analyzed phytohormones in the cambium : IAA (Auxine), ABA (Abscisic Acid), DPA (4'-dihydrophaseic acid; ABA derivative), JA (Jasmonic acid), OPDA (oxylipin 12-oxo-phytodienoic acid, JA derivative), SA (Salicylic Acid), cZOG (cis-Zeatin-O-glucoside; Cytokinin), iPR (Isopentenyl adenine riboside; Cytokinin), tZR (Trans‐zeatin riboside; Cytokinin), cZR (cis‐Zeatin riboside; Cytokinin), GA8, GA12, GA19 (Gibberellins), CS (castasterone; Brassinosteroid), 6-deoxoCS (6-Deoxocastasterone; Brassinosteroid).

| **Year of experiment** | **Species and Genotypes** | **Increased concentrations** | **Decreased concentrations** | **Total number of phytohormones affected** |
| --- | --- | --- | --- | --- |
| **Year 1  (short-term somatic memory)** | *Both species*  *P. nigra and*  *P. tremula × P. alba*  *WD-RW versus WW* | CS , GA12 | GA19, IAA  ABA , JA | 6/15 |
|  | *P. nigra*  *DRA38 specific*  *WD-RW versus WW* | DPA, cZR | OPDA, IPR, 6-deoxoCS | 5/15 |
|  | *P. nigra*  *PG31 specific*  *WD-RW versus WW* | / | / | 0/15 |
|  | *Differences among*  *2 species*  *WD-RW versus WW* | OPDA , cZOG  (P. tremula × P. alba)  cZR and tZR  (P. nigra) | OPDA , cZOG  (P. nigra)  cZR and tZR  ( P. tremula × P. alba) | 4/15 |
|  | *P. tremula × P. alba*  *3 epitypes (RNAi-ddm1, RNAi-dml and OX-dml) versus WT (WW and/or WD-RW)* | CS, 6-deoxoCS, SA (RNAi-ddm1) | ABA, IAA, GA19  (RNAi-ddm1) | 6/15 |
|  |  | 6-deoxoCS  GA19, GA8, DPA, (OX-dml) | cZOG  (OX-dml) | 5/15 |
|  |  | /  (RNAi-dml) | cZOG, tZR, JA  (RNAi-dml) | 3/15 |
| **Year 2  (inter-annual somatic memory)** | *P. nigra*  both genotypes  *WD-RW1/WD-RW2 versus WW1/WD-RW2* | GA19 | iPR | 2/15 |
|  | *P. nigra*  DRA38 specific  *WD-RW1/WD-RW2 versus WW1/WD-RW2* | GA12, IAA, DPA | 10 other phytohormones | 13/15 |
|  | *P. nigra*  PG31 specific  *WD-RW1/WD-RW2 versus WW1/WD-RW2* | SA, cZOG, tZR | GA12 | 4/15 |
| **Note** : 1. JA, OPDA larger amounts in short-term (year 1) while cZR, GA8, CS larger amounts in interannual memories (year 2).              2. JA, OPDA  larger amounts in DRA38 than PG31 (year 1) while GA12, GA19, ABA, DPA with larger amounts in PG31 than DRA38 (year 1).              3. **DPA with 10 fold more amounts  in P. nigra than  P. tremula × P. alba (year 1)**. GA19, GA12, cZOG, JA and OPDA with larger amounts in  P. tremula × P. alba than  P. nigra (year 1) in control conditions. | | | | |
